# Characterization of Cancer Evolution Landscape Based on Accurate Detection of Somatic Mutations in Single Tumor Cells

**DOI:** 10.1101/2023.10.09.561356

**Authors:** Muchun Niu, Yang Zhang, Jiayi Luo, Jefferson C. Sinson, Alastair M. Thompson, Chenghang Zong

## Abstract

Accurate detection of somatic mutations in single tumor cells is greatly desired as it allows us to quantify the single-cell mutation burden and construct the mutation-based phylogenetic tree. Here we developed scNanoSeq chemistry and profiled 842 single cells from 21 human breast cancer samples. The majority of the mutation-based phylogenetic trees comprise a characteristic stem evolution followed by the clonal sweep. We observed the subtype-dependent lengths in the stem evolution. To explain this phenomenon, we propose that the differences are related to different reprogramming required for different subtypes of breast cancer. Furthermore, we reason that the time that the tumor-initiating cell took to acquire the critical clonal-sweep-initiating mutation by random chance set the time limit for the reprogramming process. We refer to this model as a reprogramming and critical mutation co-timing (RCMC) subtype model. Next, in the sweeping clone, we observed that tumor cells undergo a branched evolution with rapidly decreasing selection. In the most recent clades, effectively neutral evolution has been reached, resulting in a substantially large number of mutational heterogeneities. Integrative analysis with 522-713X ultra-deep bulk whole genome sequencing (WGS) further validated this evolution mode. Mutation-based phylogenetic trees also allow us to identify the early branched cells in a few samples, whose phylogenetic trees support the gradual evolution of copy number variations (CNVs). Overall, the development of scNanoSeq allows us to unveil novel insights into breast cancer evolution.

## Introduction

Characterizing tumor evolution is critically important for understanding the source of intra-tumor heterogeneity (ITH). Various models have been proposed to describe this complex process ^1–8^. Meanwhile, experimental endeavors in single-cell CNV profiling have provided the first single-cell resolved evolution tree ^9,10^. In comparison to the CNV-based approach, constructing the mutation-based phylogenetic tree is also highly desired, as it allows a straightforward determination of evolution distance. It is worth mentioning that previous mutation-based phylogenetic trees were mainly based on deep sequencing of bulk samples ^11,12^ or multi-region sequencing ^13^. Although the single-cell mutation-based approach has been attempted with single-cell multiple displacement amplification (scMDA)^14^, the error rate of 1-2 false positives per million bases has constrained its application on mutation-based phylogeny construction. Therefore, a single-cell WGA allowing accurate detection of somatic mutations is greatly desired.

To achieve accurate detection of somatic mutations in single cells, here we adapt the design principle of the NanoSeq method ^15^ to single-cell chemistry. We refer to this duplex-based single-cell genome sequencing method as scNanoSeq. By duplex sequencing of both Watson and Crick strands, we can efficiently filter out DNA damage and amplification-induced errors. In total, we profiled the somatic mutations of 842 single cells from 21 human breast cancer samples (7 ER+, 7 HER2+, and 7 triple-negative breast cancer (TNBC)). As a result, we accurately determined the tumor mutation burden carried by single tumor cells, which exhibit a significant degree of inter-tumor variations (ranging from 1,444 to 49,155 somatic mutations per tumor cell).

Next, based on somatic mutations detected in single cells, we constructed the single-cell phylogenetic tree to characterize tumor evolution. We observed the following three characteristic features of evolutionary dynamics in breast cancer. First, we observed the existence of a single dominant clone and the associated clonal sweep in almost all samples, with the exception of two samples where dual dominant clones were observed. Secondly, we observed a stem structure before the onset of the clonal sweep. We define it as the stem structure because we observed a limited number of or no branches in this central node before the clonal sweep. The lack of detection of branches in the stem evolution strongly suggests the single-sweep model, though the multiple-sweep model or selective sweep model ^3^ cannot be completely ruled out. Thirdly, in the sweeping clone, we observed a branched evolution followed by the rapid genesis of a few major subclones. We showed that selection within the dominant population rapidly dropped to a baseline level, as a result, effectively neutral evolution is achieved and dominates the clonal sweep dynamics.

To alternatively verify the phylogenetic structure and the effectively neutral evolution mode, we performed 522X-713X ultra-deep tumor bulk whole genome sequencing (WGS) on 6 tumor samples. By applying a bulk-guided single-cell evolutionary path-based fingerprint analysis, we not only verified the lineage relation represented by the phylogenetic trees but also the effectively neutral evolution. Based on the single cell unique mutation load, which is accumulated during the effectively neutral evolution phase, we then determined the whole-tumor mutation burden, which ranges from 3.74×10^8^ to 1.02×10^12^ (median: 4.01×10^10^) somatic mutations for the 21 samples we profiled in this study. So, for most of the samples, the genome-scale mutational heterogeneity has already been reached. We conclude that effectively neutral evolution is the critical process for producing such a large and diverse mutation pool, which underscores the challenges in cancer treatment.

In addition, scNanoSeq data also allowed us to determine the CNVs of the single cells. Interestingly, we observed a minor level of CNV differences between the subclones of the dominant clone. At last, we performed mutational signature analysis for the somatic mutations we detected in single cells, and six major mutational signatures were captured from our single-cell data, among which we discovered a novel signature that had not been annotated before.

## Results

### Accurate detection of somatic mutations in single cells by scNanoSeq

Based on the duplex-sequencing chemistry (NanoSeq) recently developed by Abascal et al. ^15^, here we develop the scNanoSeq assay to allow the accurate detection of somatic mutations at the single-cell resolution. The scheme of scNanoSeq is illustrated in **Fig. 1a**. We first used fluorescence-activated nucleus sorting (FANS) to sort single nuclei into individual tubes. After the protease-based lysis to release genomic DNA (gDNA), we fragmented the gDNA by restriction enzyme HpyCH4V. Following the heat inactivation of the restriction enzyme, we performed dATP and ddBTP tailing to prevent the extension initiated at nicks existing on the original DNA template. The DNA fragments with the dA protruding end were then ligated to Y-shape adapters containing unique molecular identifiers (UMI). After purification, the ligation product was amplified by PCR and quantified for next-generation sequencing.

**Fig. 1.**
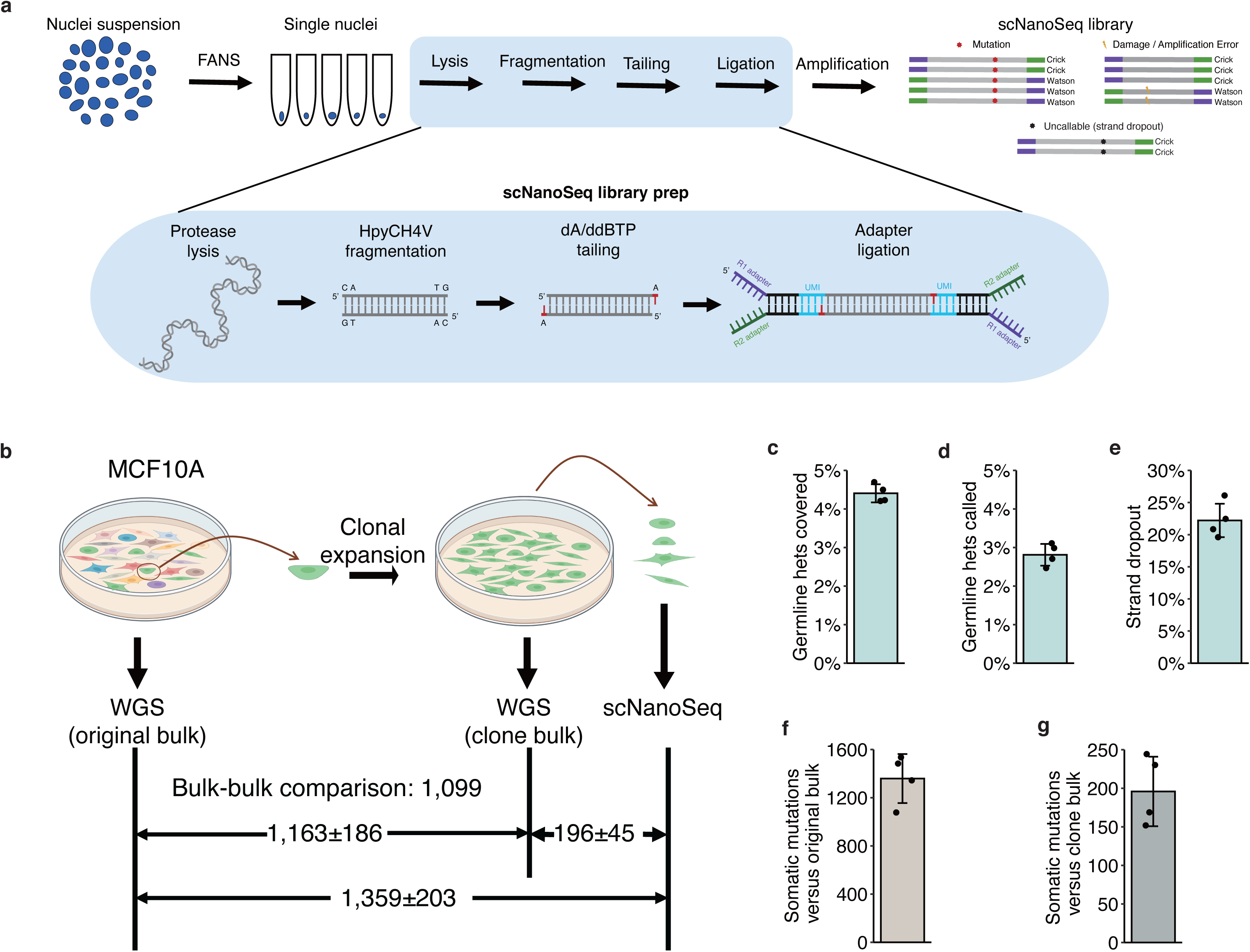
ScNanoSeq chemistry and validation. (**a**) The scheme of scNanoSeq chemistry and variant calling strategy. (**b**) The scheme of clonal expansion experiment. (**c**) Bar plot showing the percentage of germline heterozygous mutations covered by at least one strand. (**d**) Bar plot showing the percentage of germline heterozygous mutations called based on the detection of two strands. (**e**) Bar plot showing the strand dropout rate. (**f**) Bar plot showing the estimated single-cell somatic mutation numbers compared to the original bulk WGS. (**g**) Bar plot showing the estimated single-cell somatic mutation numbers compared to the clone bulk WGS. For panels (**c**)-(**g**), error bars indicate standard deviation.

The Y-shape adapter design allows us to specify the two strands of the same original gDNA fragment based on the direction of paired-end reads. True mutations are expected to be detected in the fragments with both Watson and Crick strands identified. On the other hand, since the templated DNA synthesis from nick translation has been avoided, single-base DNA damage or amplification errors will be detected in only one strand (**Fig. 1a**). Given that the single-base DNA damage rate of 1×10^-5^ per base ^16^ and Q5 DNA polymerase error rate of 5.3×10^-7^ per base ^17^, the theoretical probability of a random overlap of the same variants in both strands due to two independent DNA damage is (1×10^-5^)^2^×1/3 = 4×10^-11^ per base, corresponding to 0.26 false positive calls of somatic mutations per diploid human genome. Therefore, by requiring a candidate variant to be detected in both strands of the same fragment, the vast majority of false positives can be effectively filtered out, resulting in the accurate detection of somatic mutations in single cells.

To directly validate the accuracy of scNanoSeq, we performed a clonal expansion experiment with the human mammary epithelial cell line MCF10A (**Fig. 1b**). In brief, one single cell was picked from the original culture and then expanded for approximately 30 generations. We isolated four single diploid cells from the clonal-expanded culture to perform scNanoSeq. At first, we quantified the overall variant detection rate of scNanoseq. At the average depth of 16.8 million 2×150-bp paired-end reads from four single MCF10A cells, 4.41%±0.02% (mean ± sd) of germline heterozygous variants were covered in at least one strand, and 2.81%±0.03% (mean ± sd) of germline heterozygous variants were reliably called based on the two-strand-detection filtering criteria (**Fig. 1c-d**). Notably, the specific cutting sites of restriction enzyme HpyCH4V constrain the coverage to 28.7% region of the human genome. Therefore, the 2.81% germline variant detection rate corresponds to the effective detection rate of 9.8%. The other critical technical metric for duplex sequencing methods is the strand dropout rate since a DNA fragment is only informative when both strands are detected. Here, we determined that the strand dropout rate is 22.2%±2.6% (mean ± sd) for scNanoSeq (**Fig. 1e**). This feature shows that close to 80% of reads can be used for duplex calling and therefore warrants the cost-effectiveness in sequencing.

Next, we compared the four single cells to the original multiple-clone culture. The somatic mutations detected by this comparison comprised the somatic mutations pre-existing in the selected cell for clonal expansion and the newly accumulated mutations during the clonal expansion (**Fig. 1f**). And when we compared the four single cells to the single-cell expanded clone bulk, the somatic mutations only comprise the newly accumulated mutations in each cell during the clonal expansion (**Fig. 1f**). With the scNanoSeq data, we determined that the four single cells harbored on average 1,359±203 (mean ± sd) somatic mutations when compared to the original multiple-clone bulk (**Fig. 1f**) and 196±45 (mean ± sd) somatic mutations when compared to the single-cell expanded clone bulk (**Fig. 1g**). As a result, we determined that there were 1,163±186 (mean ± sd) somatic mutations existing in the ancestor cell selected for clonal expansion. The number of these somatic mutations can also be derived by directly comparing the clone bulk to the original bulk, which is 1,099 mutations. The consistent result between the single cell based analysis and bulk based analysis validates the accuracy of scNanoSeq in detecting mutations.

### Somatic mutation load in single breast cancer cells

With the validation of the accuracy of scNanoSeq in detecting single-cell somatic mutations, we next applied it to characterize the somatic mutations in single tumor cells of 21 frozen human breast cancer samples across three major subtypes (7 ER+, 7 HER2+, and 7 TNBC) (**Fig. 2a-b** and **Supplementary Table 1**). We indexed 7 ER+ samples as BTE1–7, 7 HER2+ samples as BTH1– 7, and 7 TNBC samples as BTT1–7.

**Fig. 2.**
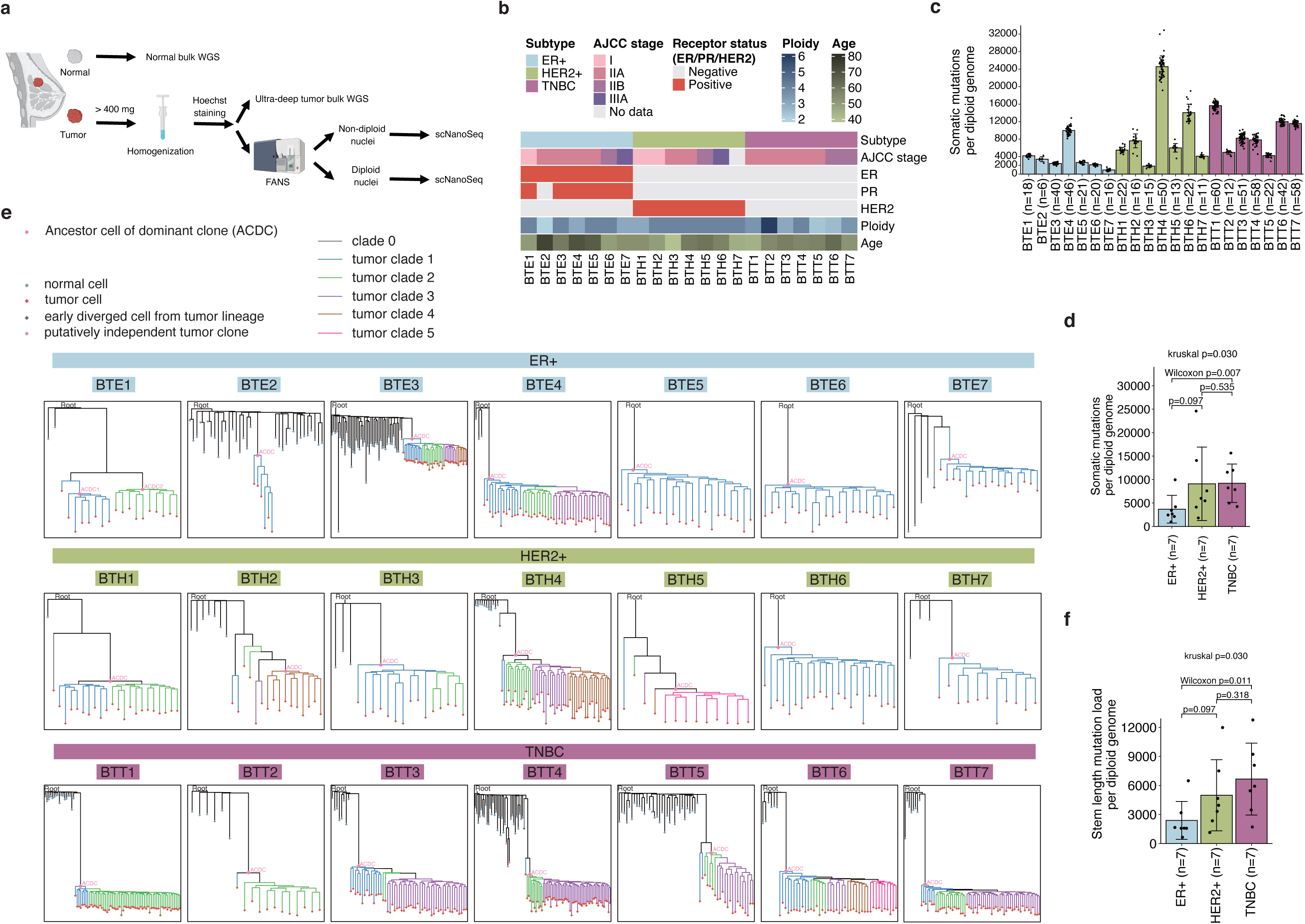
Somatic mutation load in single breast cancer cells and mutation-based evolution tree. (**a**) The experimental workflow for isolation, amplification, and sequencing of single tumor cells. (**b**) The summary of breast cancer sample information. (**c**) Bar plot showing the number of somatic mutations per diploid genome in single tumor cells. (**d**) Bar plot showing the average number of somatic mutations per diploid genome in tumor samples grouped by breast cancer subtypes. The Kruskal-Wallis test was performed among the three breast cancer subtypes. A two-sided Wilcoxon rank-sum test was performed for pairwise comparison. (**e**) Somatic mutation-based tumor evolution tree of the 21 breast cancers. (**f**) Bar plot demonstrating the stem length mutation load per diploid genome. Error bars indicate standard deviation. The Kruskal-Wallis test was performed among the three breast cancer subtypes. A two-sided Wilcoxon rank-sum test was performed for pairwise comparison.

To reduce potential regional bias, we homogenized tumor tissues with quantities of at least 400 mg. After Hoechst staining of nuclei, we performed FANS to quantify gDNA content in single nuclei, and then isolated single abnormal ploidy nuclei (candidate tumor nuclei) and diploid nuclei (putatively normal nuclei as the control) for scNanoSeq (**Supplementary Fig. 1a-y**). It is worth noting that in three samples: BTE2, BTH2, and BTT5, we observed a near-diploid population that partially overlaid with the diploid population in the histogram of Hoechst signal (**Supplementary Fig. 1f, m, w**). For these samples, we profiled single nuclei from the entire population composed of both normal and tumor nuclei.

In total, we profiled 842 single nuclei from the 21 breast cancer samples. Based on the evolution trees shown below, 619 out of the 758 candidate tumor nuclei were identified as tumor lineage. Meanwhile, none of the 84 diploid nuclei (putatively normal) were assigned to the tumor lineage (**Methods** and **Supplementary Table 2**), confirming that sorting abnormal ploidy nuclei is an effective way to sample the tumor cells.

Next, after filtering the germline mutations detected in the normal bulk samples, we detected the average number of somatic mutations in single tumor cells ranging from 44 to 1,222 (median: 278, **Supplementary Fig. 2a**) for the 21 tumor samples. When we normalized to the diploid genome, the mutation load corresponded to 963 – 24,578 somatic mutations (**Fig. 2c**). When we normalized to the ploidy numbers detected in the tumor cells, the mutation load corresponded to 1,444 – 49,155 somatic mutations per tumor cell (**Supplementary Fig. 2b**). This result shows a large degree of inter-tumor variations in the average single-cell somatic mutation load. Despite the large degree of inter-tumoral variations, the intra-tumoral variations in single-cell somatic mutation load are limited (**Fig. 2c**). We also noticed that the clinically more aggressive subtype: TNBC, exhibited significantly higher single-cell somatic mutation loads than ER+ breast cancers (**Fig. 2d** and **Supplementary Fig. 2c**). This mutation load differences among the three subtypes of breast cancers are consistent with the bulk sequencing-based measurement ^18^.

Besides the mutation load in the tumor cells, we also plotted the distribution of the mutation load in the nontumor cells (**Supplementary Fig. 3**)—the distribution peaks at 1500 mutations per diploid genome. Interestingly, we also detected 13 hypermutator cells (>3,500 somatic mutations per diploid genome) from the 217 normal lineage cells (**Supplementary Fig. 3**). This observation shows the existence of nonmalignant hypermutator cells in the tumor samples. Whether these hypermutator cells are precursor cells of cancer demand future investigation.

### Construction of phylogenetic trees based on somatic mutations detected in single cells

With the accurate detection of somatic mutations in single cells, next, we applied the probabilistic model-based algorithm CellPhy ^19^ to infer the tumor evolution tree (**Methods**). The phylogenetic trees of 21 breast cancer samples are shown in **Fig. 2e** (the tree heights are not scaled for cross-sample comparison) and **Supplementary Fig. 4** (the tree heights are scaled for cross-sample comparison). In the majority of the samples, we observed a single stem evolution followed by the rapid expansion of the dominant clone. The prevalent observation of the clonal sweep pictures confirmed the classical tumor evolution picture: the acquisition of a critical gatekeeper mutation (such as *TP53*) is likely the rate-limiting step in achieving the full scale of malignancy.

Interestingly, in two samples (Sample BTE1 and BTH1), we observed two major clones. And we call them dual-clonal sweeps. This observation shows it is possible to accommodate two major clones in one tumor. Since no single clone eventually takes over the whole population, we reason that the levels of fitness of the dual clones are very close to each other, as a result, no obvious clonal selection occurred in these two tumors.

Based on the phylogenetic trees, next we can readily obtain an approximation of the length from the root to the ancestor cells of the dominant clone (ACDC), which we refer to as stem lengths. We then use this length to estimate the mutation density at the onset of the clonal sweep (**Methods)**. Interestingly, we observed that three breast cancer subtypes demonstrated significantly different stem lengths: on average, 2,387, 4,985, and 6,657 mutations per diploid genome for ER+, HER2+, and TNBC, respectively (**Fig. 2f**). The difference in stem lengths suggests a subtype-dependent mutation load required for clonal sweep. We reason that this dependence is caused by different degrees of reprogramming required for different subtypes of breast cancer and we propose a model for explaining the relationship between reprogramming, clonal sweeping, and the subtype of tumor (please refer to the **Discussion** for details).

### Decreasing subclonal selection led to the emergence of effectively neutral evolution

Furthermore, following the ACDC in the phylogenetic trees, in most samples, we observed a series of branching events that resulted in multiple subclades in the dominant clone (**Fig. 2e**). These subclades demonstrated variations in their population sizes, which is indicative of subclonal selection. To quantify the level of subclonal selection, we introduced the unbalanced ratio between the major daughter lineage versus the minor daughter lineage at different nodes (**Methods**). Unbalanced splitting suggests selection occurred between two daughter lineages, while balanced splitting suggests neutral evolution, where no significant selection occurred (**Fig. 3a**). By applying this metric to the ACDC node and other major nodes on the deeper layers of the phylogenetic trees, we noticed the rapid decreasing trend of the unbalanced ratio across different tumor samples (**Fig. 3b-i)**. As the node goes deeper, the log_2_ of the unbalanced ratio is approaching a baseline level, and quickly we observe that zero selection consistently falls into its 95% confidence interval for all the samples examined.

**Fig. 3.**
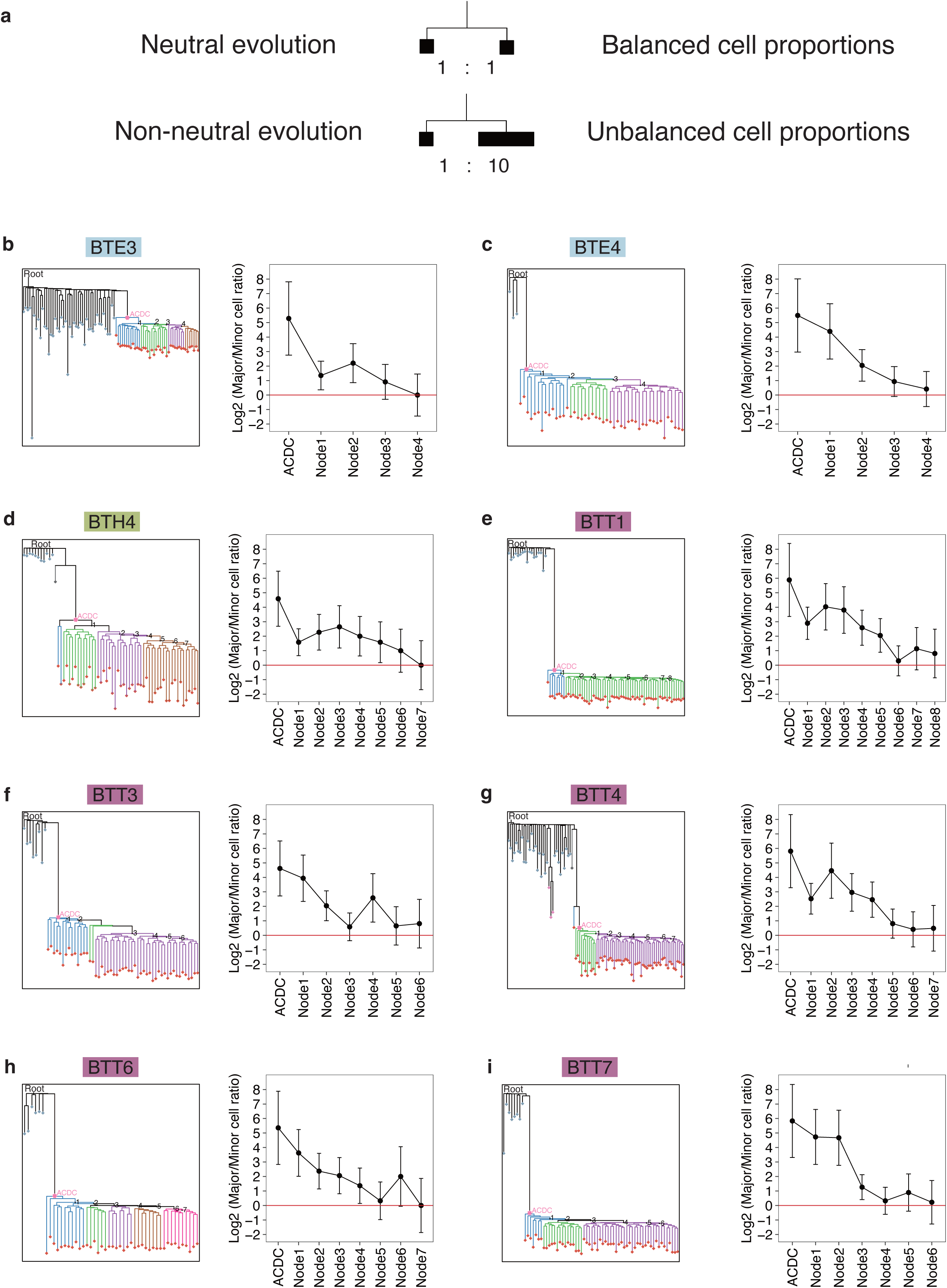
Identification of effectively neutral evolution based on single cell phylogenetic trees. (**a**) The scheme showing balanced cell proportions of both daughter lineages in neutral evolution, or unbalanced cell proportions of both daughter lineages in non-neutral evolution. (**b**-**i**) The trend of cell proportions of both daughter lineages at the ACDC node and other major nodes (labeled on the phylogenetic tree on the left) along the phylogenetic trees of tumor sample: BTE3, BTE4, BTH4, BTT1, BTT3, BTT4, BTT6, and BTT7. Error bars represent 95% confidence interval derived from the binomial proportion test. A red horizontal line at 0 stands for absolute neutral evolution.

Overall, the result above suggests that in the lineage under the ACDC, the subclonal selection decreased quickly and approached a minimal level. The minimal subclonal selection in the most recent subclade indicates that the fitness could have reached or is very close to the upper biological limit for proliferation. As a result, no significant leap in fitness can be acquired anymore. Thereby, in the end, the tumor follows effectively neutral evolution. The ultimate emergence of effectively neutral evolution follows the premise of the “big bang” evolution model as observed in colorectal cancer, in which many subclones are generated in a short period of time and grow simultaneously with extremely rare selective sweeps ^6,20^.

### Validation of the length of stem evolution in phylogenetic trees by ultra-deep sequencing

To verify the accuracy of the lineage construction and the quantification of evolution distance by the CellPhy algorithm (**Supplementary Note 1**), we performed the ultra-deep WGS for 6 breast cancer tumor bulk samples: BTE1, BTE4, BTH4, BTH5, BTT4, and BTT7 (522-713X genome coverage per sample, **Supplementary Table 3**) on the Ultima sequencing platform. We subsampled these bulk WGS datasets at different sequencing depths for somatic mutation calling using the LoFreq package^21^. As the depth increased, we observed an increased number of somatic mutations being called, but the number started to reach an upper plateau or a plateau with a small slope (**Supplementary Fig. 5a-f**). This curvature change reflects that the stem mutations and the branched part of mutations in the major subclones of the dominant clone have likely all been captured by the ultra-deep sequencing. As a result, we reach the effectively neutral evolution part of the phylogenetic tree in terms of detecting somatic mutations. And the newly detected mutations will essentially come from the doubling of the cell population driven by neutral evolution when we double the sequencing depth. In detail, the number of new detectable mutations is the product of the number of major subclones and the effective mutation rate of these subclones.

Next, the ultra-deep bulk WGS warranted clear separation of different peaks in the variant allele frequency (VAF) distribution, therefore allowing us to accurately quantify the stem length mutation density corresponding to the ACDC (**Methods**, **Supplementary Figs. 6-11, Supplementary Table 4**). As a result, we observed that the approximation of stem length estimated by CellPhy aligned very well with the bulk WGS-based quantification (**Fig. 4a**, R^2^ = 0.99). The differences between both methods were 67-324 somatic mutations (median: 305) per diploid genome. This result verified the robustness of stem length inferred by CellPhy.

**Fig. 4.**
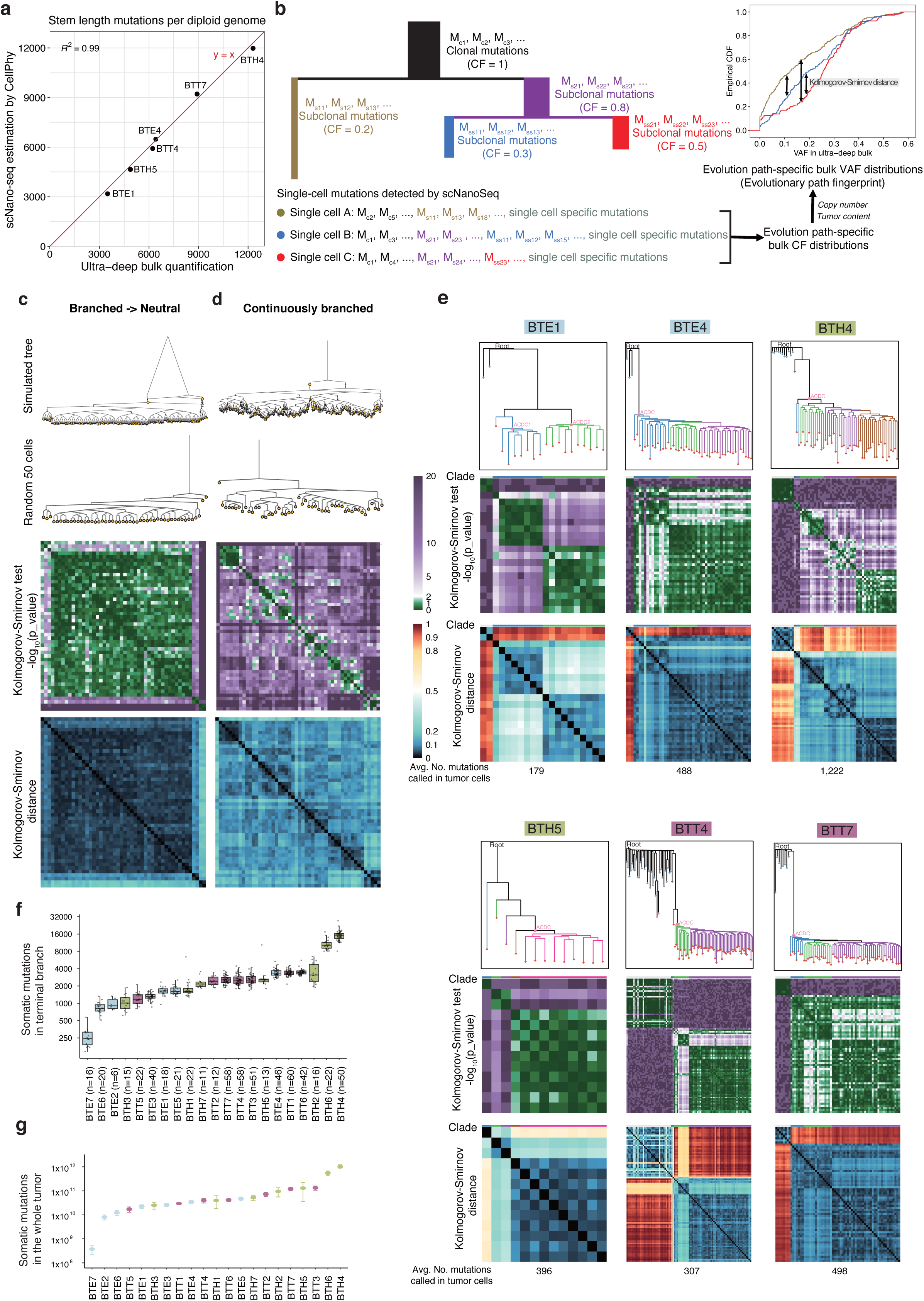
Validation of tree structure and effectively neutral evolution mode by ultra-deep bulk WGS. (**a**) The scatter plot comparing the stem length mutation density quantified by ultra-deep bulk WGS versus estimated based on CellPhy-inferred phylogenetic tree. The red line represents the diagonal line y=x. (**b**) An example scheme demonstrating the bulk-guided single-cell evolutionary path analysis framework. Mutations accumulated during stages of tumorigenesis affect different cell fractions (CF): for example, clonal mutations affect all tumor cells (CF=1), while subclonal mutations affect only part of the cells (0<CF<1), and the CF values depend on the evolution structure. As a result, for the somatic mutations detected in each single cell, their CF value distribution represented a unique evolutionary path of the cell. When the bulk copy number and tumor content is taken into account, the CF value for each variant can be uniquely mapped to a specific VAF value that is observable in the bulk WGS. Therefore, the bulk VAF distribution can be used as a fingerprint of single cell evolutionary path, and different cell lineages could be identified by the Kolmogorov-Smirnov test. (**c-d**) Evaluation of the bulk-guided single-cell evolutionary path analysis framework on (**c**) a simulated dataset assuming branched evolution followed by neutral evolution, or (**d**) a simulated dataset assuming continuously branched evolution. The upper two panels demonstrate the simulated full evolution tree structures, or evolution tree structures with 50 randomly sampled cells. For each single cell, an average of 1,000 mutations were randomly sampled for pairwise Kolmogorov-Smirnov test. The p values of the pairwise Kolmogorov-Smirnov test are displayed in the heatmaps as the third panel. The pairwise Kolmogorov-Smirnov distances are displayed in the heatmaps as the lower panel. (**e**) Pairwise Kolmogorov-Smirnov test statistics of single cell VAF fingerprints in breast cancer samples. The CellPhy-inferred evolutionary trees (same as Fig. 2e) for different tumor samples are displayed as the upper panel. The p values of the pairwise Kolmogorov-Smirnov test are displayed in the heatmaps as the middle panel. The pairwise Kolmogorov-Smirnov distances are displayed in the heatmaps as the lower panel. For all the heatmaps, the single cells were ordered the same as the phylogenetic trees. The average number of mutations called per tumor cell is labeled at the bottom. (**f**) Approximation of somatic mutation load per tumor cell in the terminal branch. (**g**) Approximation of whole tumor mutation burden. For (**f**) and (**g**), error bars stand for standard deviations derived across single-cell estimations.

### Validation of neutral evolution by VAF fingerprint derived from ultra-deep sequencing data

To further verify the lineage structure of the phylogenetic tree inferred by CellPhy, we introduced a bulk-guided single-cell evolutionary path analysis framework (**Fig. 4b**). Since the detection of somatic mutations by scNanoSeq is on the single molecule basis, the mutation detection efficiency does not depend on its VAF in the bulk sample, thereby allowing unbiased detection of somatic mutations accumulated throughout the evolutionary path of each single cell. We reasoned that when we trace the somatic mutations detected in each single cell back to the ultra-deep bulk WGS data, the VAFs of these mutations are expected to follow a specific distribution, which can be used as the fingerprint of an evolutionary path that describes the lineage history of this cell. For convenience, we refer to the VAF distribution of the scNanoSeq-called somatic mutations in ultra-deep bulk sequencing data as the VAF fingerprint of single cells.

To examine this VAF fingerprint framework, we first simulated a phylogenetic dataset with the assumption of branched evolution followed by neutral evolution (**Methods**, **Fig. 4c**) and the continuously branched evolution model with notable selection (**Methods**, **Fig. 4d**). We randomly sampled 50 single cells and performed the pairwise Kolmogorov-Smirnov (KS) test to characterize the differences in their VAF fingerprints. We observed that in the clades where absolute neutral evolution is achieved (**Fig. 4c**), cell pairs within the clade consistently display statistically undistinguishable VAF fingerprints, as indicated by insignificant p values (> 0.01) and relatively small Kolmogorov-Smirnov distances. In contrast, the cells from different clades display different VAF fingerprints, resulting in multiple blocks along the diagonal (**Fig. 4d**). We further tested the performance of KS statistics under different numbers of somatic mutations detected per cell (**Supplementary Fig. 12a-b**). Although fewer sampling of somatic mutations resulted in decreased statistical power, a notable portion of inter-branch comparisons remain statistically significant in the continuously branched evolution mode, therefore rejecting neutral evolution under this simulation (**Supplementary Fig. 12b**). These simulation data provide a direct validation of the statistical power of VAF fingerprint analysis in identifying unbalanced branched evolution resulting from noticeable selection.

Having validated the statistical power to characterize neutral evolution versus branched evolution, next, we applied the VAF fingerprint framework to the 6 tumor samples with ultra-deep bulk WGS (**Fig. 4e**). As a result, only a few blocks of VAF fingerprints were observed on the diagonal, and the single cell pairs within the same lineage display the same VAF fingerprints at large. The same VAF fingerprints within the same lineages suggest minimal selection in the most recent clade, in other words, effectively neutral evolution is reached.

In contrast, the VAF fingerprints between different CellPhy-defined clades were largely distinguishable from each other. It is worth pointing out that such a separation between subclades matched with phylogenies assigned by CellPhy. For example, within the Clade 2 and 3 of BTH4 sample (**Fig. 4e**), the fingerprint analysis resulted in a clear separation of subclade structures attributed to branched evolution, as smaller blocks can be seen along the diagonal line in the heatmaps, and the blocks are consistent with the lineage assignment. As a result, the VAF fingerprint provided an independent validation of the accuracy of lineage construction by CellPhy algorithm.

### Estimation of whole tumor mutation burden on the basis of neutral evolution

As effective neutral evolution is achieved, we are able to quantify the whole tumor mutation burden on the basis of an effective exponential expansion model. Based on the averaged single-cell unique mutation load *L* and effective cell doubling number *n*, we can calculate the effective mutation rate per cell doubling as 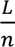, and thereby the total number of mutations acquired during the exponential growth phase as 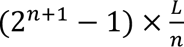 (**Methods**). As a result, we determined the whole-tumor mutation burden, which ranges from 3.47×10^8^ to 1.02×10^12^ (median: 4.01×10^10^) mutations across the tumor samples (**Fig. 4f-g, Supplementary Table 5**). Therefore, we conclude that the whole-tumor mutation burden already reached the magnitude of the human genome size. This result shows that effectively neutral evolution is critical in generating genome-wide mutational heterogeneity.

From the effective neutral evolution, we also estimated the effective mutation rate, which ranges from 11 to 507 mutations (median: 87) per division (**Supplementary Table 5**). These mutation rates are on the same scale as previous experimental measurements on cell lines: immortalized cell line RPE1 acquired 39 mutations per division while cancer cell line HT115 could acquire as many as 173 mutations per division ^22^. With such a high mutation rate, it is worth pointing out that even in a single-cell expanded clone, a large number of mutations (on the scale of tens of millions) could be accumulated through effective neutral evolution, therefore, cautions need to be taken when using a single-cell expanded clone as an isogenic clone.

Overall, the genetic diversity (in both tumor samples and cell line samples) will provide substantial survival advantages to tumor cells when facing drug treatment by either directly blocking the drug-binding pocket (drug resistance), or bypassing the signaling pathways for growth (drug tolerance)^23,24^.

### Mutation-based phylogenetic trees identify early-branched cells

With the scNanoSeq data, we can also determine the CNVs in the same single cells. We performed the CNV analysis followed by the hierarchical clustering using the Ginkgo algorithm ^25^ (**Supplementary Table 6**, **Supplementary Fig. 13-33**). For a few samples in which we detected early branches, we noticed that the CNV profiles of the early branched cells have significant differences from the dominant clones. For example, in the phylogenetic tree of BTT4 (**Fig. 5a**), an early-branched cell (sc304) and a middle-late-stage cell (sc033) were detected. In **Fig. 5b**, we include a randomly selected normal cell (sc307) and one cell from the dominant clone (sc343) for comparison. We observed significant differences between the CNVs of the early branched cell sc304 and sc343 from the dominant clone. We reason that with the large degree of differences in CNVs, this cell may not be recognized as the cell in the tumor lineage by the CNV-based clustering. Indeed, in the clustering analysis by Ginkgo, both early-branched sc304 and later diverged sc033 were assigned to the nontumor cell cluster (**Fig. 5c**).

**Fig. 5.**
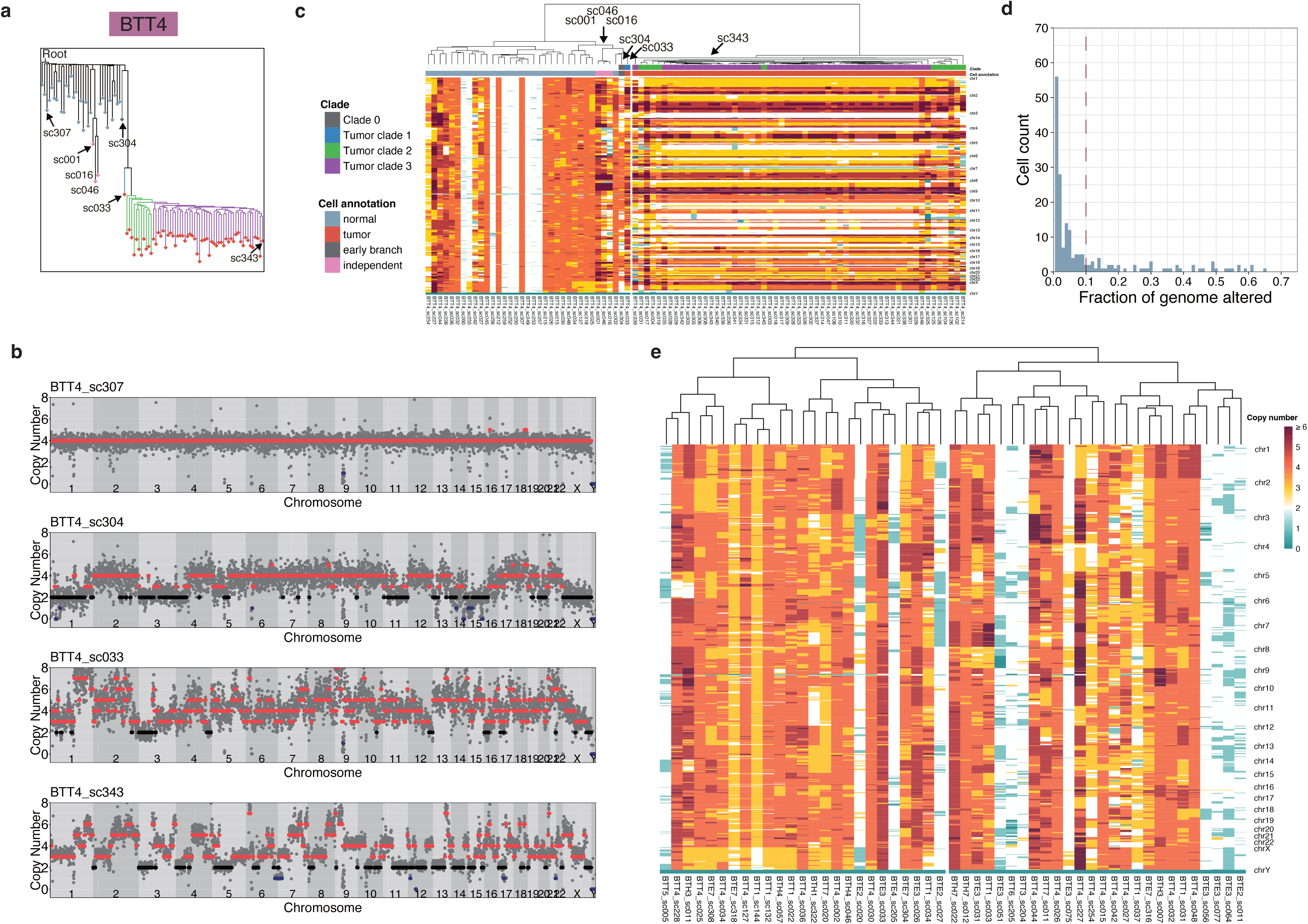
Gradual CNV evolution in stem evolution and CNVs in nontumor cells. (**a**) The evolution tree of BTT4 based on mutations, with selected cells labeled. (**b**) Copy number profiles of the selected cells. (**c**) The CNV heatmap and the lineage relationship between cells based on their CNVs. **(d)** Histogram showing the fraction of genome altered by CNV in normal lineage cells. A cell is considered with major CNV if >10% of its genome is altered. **(e)** The CNV patterns in the non-tumor cells with major CNV changes across different samples.

In an ER+ sample with early branched cells, BTE7, we also observed that the early diverged cell, sc321, only developed a few regional amplifications (**Supplementary Fig. 34**). In comparison, sc307 at the top layer of the clonal sweep (**Supplementary Fig. 34**) shows much more progressed CNV patterns and highly consistent with the dominant clone (**Supplementary Fig. 34**). In a HER2+ sample with early branched cells, BTH4 (**Supplementary Fig. 35**), we were able to sample one early branched cell: sc005, and its CNVs are also significantly different from the dominant clone (**Supplementary Fig. 35**).

These results suggest that different clones could be generated along the stem evolution but with likely very limited proliferation compared to the dominant clone. As a result, we did not capture early branched cells in most of the samples. However, based on the few samples from which we have captured their early branched cells, we reason that it is possible that copy number changes follow the gradual evolution along the stem evolution instead of a single punctuated event ^9,10^.

### Detection of non-tumor lineage lesion cells

Among the cells isolated from the abnormal ploidy population by FANS, we frequently detected non-tumor lineage lesion cells with clear CNVs. In 51 out of 217 non-tumor lineage cells, we observed clear CNVs with more than 10% of genome altered (**Fig. 5d**). The CNVs of these cells across different samples are shown in **Fig. 5e**. Interestingly, we observed noticeable deletion of chromosome X across different cells from different samples.

About the origin of these nontumor prelesion cells, it is possible that they are induced by the stressful tumor microenvironment. On the other hand, this high percentage of nontumor prelesion cells could also correspond to the benign precursor cells existing before the tumor development. It is possible that the tumor lineage originates from one of these precursor cells. Interestingly, in sample BTT4, we observed an extra development of a nontumor prelesion clone represented by sc001, sc016, and sc046 (pink cells in **Fig. 5a, c**), indicating that these precursor cells can definitely expand once the appropriate mutations and CNVs were acquired to promote the proliferation. Future investigation of whether these nontumor high-ploidy cells are the precursor lesions before tumor development and how these different clones are initiated is greatly desired.

### Mutational signatures analysis identifies characteristic signatures in tumor samples

With the somatic mutations detected in single cells, we further explore the underlying mutagenesis mechanisms in our samples by mutational signature analysis of the somatic mutations detected in single tumor cells. We first observed clear intertumoral differences in both the 6-type mutation spectrum (**Supplementary Fig. 36**) and 96-type single base substitution (SBS) mutational profiles (**Supplementary Fig. 37**). More importantly, the large number of somatic mutations we detected in single tumor cells allows robust SBS mutational signature analysis at the single-cell resolution. Through non-negative matrix factorization (NMF) implemented in the MutationalPatterns R package ^26^, we successfully deconvoluted 6 SBS signature components: Sig1-6 (**Fig. 6a** and **Supplementary Table 7)**.

**Fig. 6.**
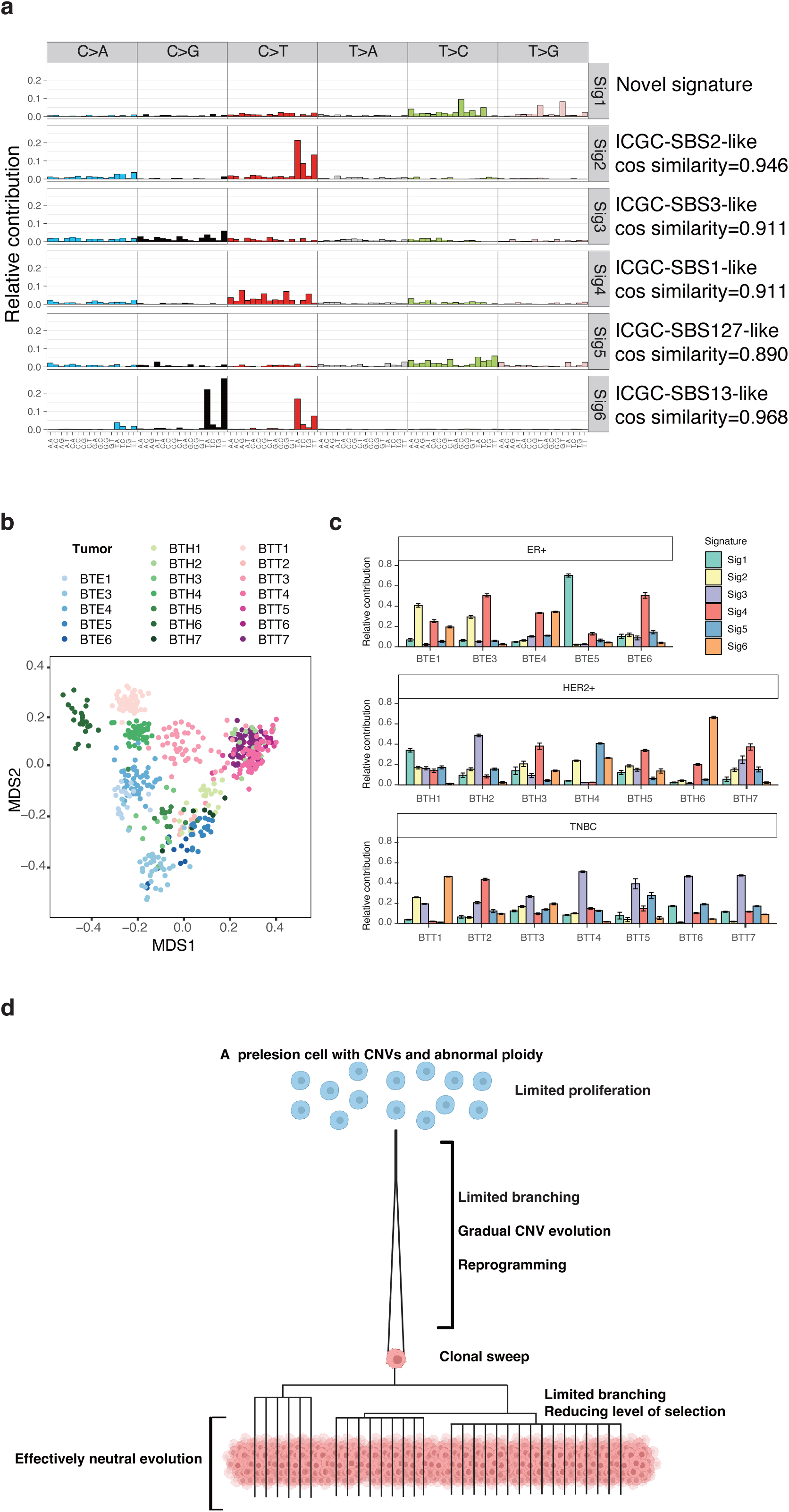
Mutational signatures and scheme of tumor evolution in breast cancer. (**a**) Six mutational signature components extracted by NMF. (**b**) The MDS plot for visualization of mutational signature relative contributions in single tumor cells. (**c**) Contributions of the six extracted signatures in the 21 breast cancers. Error bars indicate the standard error of the mean. (**d**) The scheme of tumor evolution in breast cancer.

By projecting these signatures onto the Signal cancer signature database generated based on three independent cancer cohorts (ICGC, GEL, and Hartwig) ^27^, we connected five signatures (Sig2-6) to the known signatures with cosine similarity greater than 0.85 (**Fig. 6a**, **Supplementary Fig. 38a-c**, and **Supplementary Table 8**). In summary, Sig2 and Sig 6 were connected to SBS2-like and SBS13-like signatures, respectively, and both SBS signatures were attributed to the activity of the APOBEC family of cytidine deaminases ^28,29^. Sig3 was connected to SBS3-like signatures, a signature representing defective homologous recombination-based DNA repair; Sig4 was connected to SBS1-like signatures, which results from spontaneous deamination of 5- methylcytosine. Sig5 was related to SBS127-like signatures with an unknown etiology. Finally, Sig1 is a novel signature to which we did not identify a similar signature in the Signal cancer signature database.

Through the linear combinations of Sig1-6, we then reconstructed the mutational signatures in single tumor cells (**Supplementary Fig. 39a-b, Supplementary Table 9**). The cosine similarity of reconstructed signatures versus the observed signatures reached beyond 0.8 in 90.0% of tumor cells (557 out of 619). The degree of discrepancy is less than 0.2 of cosine similarity, primarily due to the noise caused by the limited number of somatic mutations detected in the cells from the samples with low mutation burden (**Supplementary Fig. 39a**). After filtering out the cells with less accurate signature reconstruction (**Supplementary Fig. 39c**), we performed multidimensional scaling (MDS) analysis on the single-cell signature relative contribution matrix for visualization (**Fig. 6b**). Overall, while we can observe some separation between ER+ samples and TNBC samples, the samples within the same subtype are quite spread-out, indicating there are no characteristic subtype-specific mutational signatures.

Next, we plotted the relative contributions of the six signatures to the mutation burden for each tumor in **Fig. 6c**. We observed that the tumor samples often have 1 to 2 characteristic mutational signatures. For example, Sig6 (related to the activity of the APOBEC family of cytidine deaminases) is the major signature for BTT1, Sig4 (related to spontaneous deamination of 5- methylcytosine) is the major signature for BTT2, and Sig3 (related to defective homologous recombination-based DNA repair) is the major signature for BTT4-7.

## Discussion

In recent single-cell studies of tumor evolution, the CNV-based approach predominately captures a long stem node followed by many short branches of clones – the so-called “palm tree” like structure ^9,10^. Compared to the CNV-based single-cell approach, constructing the single-cell lineage tree based on somatic mutations is greatly desired since mutation-based phylogenetic analysis allows a more accurate construction of the evolution tree and a more straightforward estimation of evolution distance. With the development of the single-cell version of NanoSeq chemistry, we achieved accurate detection of somatic mutations in single cells and successfully constructed phylogenetic trees based on their somatic mutations.

From the phylogenetic trees, we observed “palm tree” structures similar to the tree observed in the CNV-based approach. The long stem indicates that one critical mutation (e.g. a critical gatekeeper mutation) is required for the genesis of sweeping clones. With the quantification of evolution distance, we observed that the stem evolution length of the TNBC subtype is significantly longer than the ER+ subtype. This phenomenon is likely dependent on the reprogramming processes required for the different subtype formation.

Previous studies have shown that TNBC cancer originates from luminal cells (rather than basal stem cells), the same as ER+ and HER2+ subtypes ^30–32^. However, different from ER+ and HER2+ subtypes, these luminal cells are reprogrammed into basal-like cells in TNBC. As a result, the long stem lengths observed in TNBC trees (**Fig. 2e**) suggest that it takes a long time to reprogram luminal cells into basal-like cells. In other words, because the reprogramming process in TNBC takes a long time, a large mutation load will be accumulated in its stem evolution. In contrast, in most of the ER+ samples, we observed that the mutation loads accumulated in the stem evolution were similar to the somatic mutation loads observed in the nontumor cells (**Fig. 2e**), indicating a limited or no reprogramming is needed for the ER+ subtype. It is worth pointing out that newly acquired CNVs and mutations can also be responsible for boosting phenotypic plasticity to facilitate reprogramming and influence the reprogramming outcome ^33,34^. In addition, the lack of major branches in our phylogenetic trees further indicates that the mutations and CNVs that occurred on the stem evolution could contribute more to reprogramming than to proliferation.

The next important question is how the clonal sweep is integrated with the reprogramming process in the stem evolution. First, we reason that the time that the tumor-initiating cell used to acquire the critical clonal-sweep-initiating mutation sets the time limit for the reprogramming process. For example, by random chances, the tumor-initiating cell could acquire the critical mutation before the cell has the time to be transdifferentiated from luminal-like to basal-like cells, as a result, an ER+ or HER2+ subtype tumor emerges. On the other hand, it is also possible that the tumor-initiating cell acquires a critical mutation at a rather late stage. During the long stem evolution, the acquisition of mutations/CNVs will promote cell plasticity, dedifferentiation, and transdifferentiation. And by the time of the late acquisition of critical gatekeeper mutations, the cell has already transformed into a basal-like cell, as a result, a TNBC subtype tumor emerges.

To summarize, we speculated that the subtype-dependent stem evolution is associated with various degrees of reprogramming for different breast cancer subtypes. We further proposed that the tumor subtypes closely depend on when the critical gatekeeper mutation was acquired. We refer to this model as a reprogramming and critical mutation co-timing (RCMC) subtype model. Furthermore, when we examined tree structures within the dominant clone, we observed evidence for neutral evolution ^35^. Recently Williams *et al.* have explored the bulk sequencing data for the evidence of neutral tumor evolution ^7^, however, the approach used in their study is under debate ^36–42^. In our study, by integrative analysis of both single-cell sequencing and ultra-deep bulk sequencing data, we provided a new insight that reconciled the debate: in the clonal sweep, branched evolution occurred first; but after that, the selection rapidly dropped, as a result, effectively neutral evolution is reached within the major subclones.

Next, based on neutral evolution, we estimated that the whole-tumor mutation burden could reach as many as 10^8^-10^12^ mutations per sample. As a result, we conclude that a full-blown tumor is intrinsically a comprehensive genetic pool in which mutations of the vast majority of genes have been sampled. This quantification underscores the critical role of neutral evolution, instead of selective evolution, in creating genome-scale mutational heterogeneity, which underscores the source of the preexisting drug-resistant variants.

Here we further speculate that the underlying reason for the prevalent observation of effectively neutral evolution is the potential existence of an upper limit of fitness, which is physically constrained by the cellular system itself. With the continuous accumulation of mutations and CNVs, the fitness of the tumor cells will gradually approach the upper limit of fitness. As the fitness of the tumor cells gets close to the upper limit, all the cells and their represented clones will expand at a similar rate, which essentially results in effective neutral evolution. Interestingly, we observed two tumor samples with dual clones. The observation of the co-existence of dual clones in one tumor further suggests the existence of a biological upper limit of fitness. In the parallel evolution of early diverged branches, two cells from two distal branches likely have acquired similar close-to-limit fitness. As a result, two independent clones began to sweep the total population together.

Besides drug resistance, the mutations that boost drug tolerance could also be sampled in the large mutation pool of tumor cells, A recent study by Razavi *et al.* has shown that at least tens of genes have been identified to be capable of endocrine therapy tolerance ^24^, therefore we can expect that the number of mutations that grant drug tolerance will be over one hundred (**Supplementary Note 2**). Due to the neutral evolution and the diverse pool of mutations existing in tumors as well as cultured cells (even in single-cell expanded clones^43^), we would expect that the cells with various degrees of drug tolerance have already emerged from neutral evolution.

With scNanoSeq data, we can also derive the CNV profiles for the same single cells. In the early branched cells, we observed a relatively limited degree of CNVs compared to the high-degree CNVs observed in the dominant clones, indicating a gradual acquisition of copy number changes along the stem evolution. Surprisingly, besides the tumor cells, nontumor lesion cells with abnormal ploidy and CNVs were also frequently observed. Those cells could correspond to the preexisting precursor cells, and the tumor can develop from one of them.

Overall, our study suggests the following the tumor evolution landscape of breast cancer (**Fig. 6d**). Along the observed long-stem evolution, the tumor-initiating cell continuously acquires CNVs and mutations, which promote not only growth but more importantly, the reprogramming process. This process is slow and has different branches. Eventually, one of the tumor-initiating cells acquires the critical mutations such as *TP53* or other mountain mutations^5^ to ignite the clonal sweep. Following the RCMC subtype model, the subtypes of tumors are essentially determined by the timing of critical mutations for clonal sweep and the degree of reprogramming at the time.

During the clonal expansion, multiple subclones with close-to-biological-limit fitness (i.e. close-to-saturated) are formed in a short period of time (i.e. the diverging nodes are very close to each other in phylogenetic trees). This process is similar to what is described in the big-bang evolution model ^6^. Because of the similar fitness of these subclones, the selection power between them is minimal, which essentially represents the effectively neutral evolution. Following the progress of the neutral evolution, the total number of somatic mutations of a full-blown tumor could reach the whole genome scale.

To summarize, supported by our study of breast cancer, we believe that the scNanoSeq method and the analytical approach developed in this study can be readily applied to other types of cancer to shed new light on cancer evolution.

## Acknowledgments

We are grateful to the McNair family for the support through McNair Scholarship. We thank other Zong lab members for their help in this project. We would like to thank Carol Chenault, Bryant Lee McCue, and Susan G. Hilsenbeck for assisting in the tissue request. We also would like to thank Pengfei Liu, Becky Maywald, and Kim Worley for their support in Ultima sequencing. We also would like to thank Susan Rosenberg and Brendan Lee for their kindly support. FANS experiments were performed in the Cytometry and Cell Sorting Core at Baylor College of Medicine with funding from the CPRIT Core Facility Support Award (CPRIT-RP180672), the NIH (P30 CA125123, S10 RR024574, and S10 OD025251), and the assistance of Joel M. Sederstrom.

## Author contributions

C.Z. and M.N. designed the project. M.N. developed the scNanoSeq chemistry and performed the major experiments. J.L. performed the FANS. J.C.S. performed the Ultima WGS. A.M.T. provided the breast cancer samples. M.N., Y.Z., and C.Z. performed the bioinformatic analysis. M.N. and C.Z. wrote the manuscript. C.Z. supervised the project.

## Data and materials availability

The raw sequencing data of scNanoSeq and matched normal bulk WGS have been deposited in NCBI Sequence Read Archive (SRA) with BioProject accession code PRJNA958124. The raw sequencing data of ultra-deep bulk WGS data have been deposited in NCBI Sequence Read Archive (SRA) with BioProject accession code PRJNA962101.

## Supplementary Materials

Materials and Methods

Supplementary Notes 1 to 2

Supplementary Figs. 1 to 39

Supplementary Tables 1 to 9

