## Supplementary Materials for "Characterization of Cancer Evolution Landscape Based on Accurate Detection of Somatic Mutations in Single Tumor Cells"

**This file includes:**

**Materials and Methods**
**Supplementary Notes 1 to 2**
**Supplementary Figs. 1 to 39**
**Supplementary Tables 1 to 9**

### Materials and Methods

scNanoSeq protocol

Single cells/nuclei were FANS sorted into individual PCR tubes containing 2.5 µL protease lysis buffer [30 mM Tris-HCl (pH 8.0), 2 mM EDTA, 20 mM KCl, 0.1% Triton X-100, 0.36% NP-40, 0.8 mg/mL protease (Qiagen, Cat. 19157)]. The reaction was incubated at 50°C for 2 h, 75°C for 20 min, and 85°C for 5 min for cell lysis. Next, 2.5 µL fragmentation mix [0.5 µL CutSmart Buffer (New England BioLabs, Cat. B7204), 0.15 µL HpyCH4V (New England BioLabs, Cat. R0620L), 1.85 µL nuclease-free water] was added to individual tubes, and the reaction is incubated at 37°C for 15 min followed by heat inactivation of the restriction enzyme at 65°C for 20 min. Next, dA/ddBTP tailing was performed by adding 10 µL tailing mix [1 µL NEBuffer4 (New England BioLabs, Cat. B7004S), 7.35 µL nuclease-free water, 1.5 µL dATP/ddBTP mix (1 mM each, Trilink Biotechnologies), 0.15 µL Klenow fragment 3' exo- (New England BioLabs, Cat. M0212L)] and incubated at 37°C for 30 min. Immediately after tailing, 22.4 µL ligation mix [2.24 µL NEBuffer4 (New England BioLabs, Cat. B7004S), 15.53 µL nuclease-free water, 3.74 µL 10 mM ATP, 0.33 µL 15 µM xGen CS adapter (IDT, Cat. 1080799), 0.56 µL 400 U/µL T4 DNA ligase (New England BioLabs, Cat. M0202L)] was added, and ligation was performed at 20°C for 22 min. The ligation product was immediately purified using 0.95X Ampure XP beads (Beckman Coulter, Cat. A63881), and eluted in 15.5 µL nuclease-free water. To avoid sample loss, we flicked the tubes for all the mixing steps instead of pipetting.

0.5 µL of the product was used for quantitative PCR (qPCR) to quantify the library complexity. An Illumina Truseq library with known concentration was used as calibration standards. The qPCR was prepared by mixing 5 µL iTaq Universal SYBR Green Supermix (Biorad, Cat. 1725125), 0.25 µL 10 µM Truseq5 primer (ACACTCTTTCCCTACACGAC), 0.25 µL 10 µM Truseq7 primer (GTGACTGGAGTTCAGACGTGT), 4 µL nuclease-free water, and 0.5 µL purified product. The qPCR was performed on a Roche LightCycler 96 machine following the program: 94°C for 2 min, 30 cycles of [94°C for 20 s, 58°C for 20 s, 72°C for 1 min], followed by the high-resolution melting curve.

The remaining 15 µL product was used for amplification, and single-cell indices were introduced by indexed PCR primers. 5 µL 10 µM P5_index5_Truseq5 primer (AATGATACGGCGACCACCGAGATCTACAC[8-base-index]ACACTCTTTCCCTACACGACGCTCTTCCGATCT), 5 µL 10 µM P7_index7_Truseq7 primer (CAAGCAGAAGACGGCATACGAGAT[8-base-index]GTGACTGGAGTTCAG ACGTGTGCTCTTCCGATCT), and 25 µL NEBNext Ultra II Q5 master mix (New England BioLabs, Cat. M0544X) was added. The PCR amplification was performed following the program: 98°C for 30 s, 17 cycles of [98°C for 10 s, 73°C for 75 s], and 73°C for 5 min. The PCR product was purified twice with 0.7X Ampure XP beads and eluted in 20 µL nuclease-free water. Single-cell libraries were pooled together and sequenced on an Illumina NovaSeq 6000 platform S4 flowcell with 2×150 bp paired-end mode. The target depth is 5~6 reads per fragment based on pre-amplification qPCR quantification.

Cell culture

MCF10A cells were obtained from American Type Culture Collection (Cat. ATCC-CRL-10317), which is referred to as the original culture. The cells were cultured in Dulbecco's modified Eagle's medium/F12 (Invitrogen, Cat. 11330-032) with 5% horse serum (Invitrogen, Cat. 16050-122), 20 ng/mL epidermal growth factor (PeproTech), 0.5 µg/mL hydrocortisone (Sigma-Aldrich, Cat. H-0888), 100 ng/mL cholera toxin (Sigma-Aldrich, Cat. C-8052) and 10 µg/mL insulin

(Sigma-Aldrich, Cat. I-1882), 100 IU/mL penicillin, and 100 μg/mL streptomycin at 37°C and 5% $CO_2$. Cell culture was passaged every two to three days. The single-cell expansion experiment is performed by diluting the cells to 10 cells/mL in the conditioned media. 100 μL of diluted cells were aliquoted to individual wells of a 96-well plate. Four hours later, each well was checked by microscopy to exclude the empty wells and doublet wells. Only the wells containing exactly one single cell were kept for further expansion. After approximately 25 generations of expansion, the cells were harvested for gDNA extraction using the Zymo Quick-DNA Miniprep kit (Zymo, Cat. D3024) following the manufacturer's instructions. Single cells were isolated at approximately the $30^{th}$ generation.

Frozen human breast cancer samples

Frozen human breast cancer samples and matched normal tissues were obtained from Tissue Bank at BCM Lester and Sue Smith Breast Cancer Center. The sample information is summarized in **Fig. 2b** and **Supplementary Table 1**.

Tissue processing

Whenever available, > 400 mg of frozen tumor tissue was chopped into small pieces with a blade. The tissue was resuspended in 1.5 mL homogenization buffer [250 mM sucrose, 25 mM KCl, 5 mM $MgCl_2$, 10 mM Tris-HCl (pH 8.0), 0.1% Triton X-100] and transferred to a Dounce homogenizer (Wheaton). Homogenization was performed by 6 strokes of the loose pestle and 12 strokes of the tight pestle. The homogenate was passed through a 70-μm cell strainer and centrifuged at 1,500 g for 6 min at 4°C. The supernatant was discarded, and the pellet containing nuclei was resuspended in 1 mL phosphate-buffered saline (PBS) containing 1 μg/mL Hoechst 33342 (Invitrogen, Cat. H3570). Hoechst staining was performed at room temperature for 10 min. One aliquot of the sample was used for gDNA extraction using the Zymo Quick-DNA Miniprep kit (Zymo, Cat. D3024). The samples were proceeded for FANS using a BD Aria II or a BD Aria III flow cytometer. The gating strategy is demonstrated in **Supplementary Fig. 1**. The diploid single nuclei and non-diploid single nuclei were sorted into individual PCR tubes with single-cell precision. To extract the matched normal tissue gDNA, we followed the same procedure to generate the single nucleus suspension for gDNA extraction.

Illumina bulk WGS

Bulk gDNA was fragmented with the hyperactive Tn5 transposase. We assembled the transposition mix: 7.5 μL tagmentation DNA buffer (Illumina, Cat. 15027866), 0.7 μL tagment DNA enzyme (Illumina, Cat. 15027865), 50 ng gDNA, and nuclease-free water to bring the volume to 15 μL. Transposition was performed at 55°C for 16 min. 3 μL 0.2 M EDTA was added to quench the reaction, followed by 50°C heating for 30 min to release the transposase. Next, 32 μL amplification mix [3 μL 0.2 M $MgCl_2$, 2 μL 10 μM N5XX index primer (AATGATACGGCGACCACCGAGATCTACAC[8-base-index]TCGTCGGCAGCGTC), 2 μL 10 μM N7XX index primer (CAAGCAGAAGACGGCATACGAGAT[8-base-index] GTCTCGTGGGCTCGG), 25 μL NEBNext Ultra II Q5 master mix (New England BioLabs, Cat. M0544X)] was added, and PCR amplification was performed following the program: 72℃ for 3 min, 98℃ for 30 s, 7 cycles of [98℃ for 10 s, 58℃ for 15 s, 72℃ for 1 min], 72℃ for 3 min. The amplified product underwent 0.55X/0.7X two-sided Ampure XP bead size selection, and the library was sequenced on an Illumina NovaSeq 6000 platform S4 flowcell with 2×150 bp paired-end mode targeting 30X whole genome coverage per sample.

Ultima ultra-deep bulk WGS

2 μg tumor bulk gDNA was diluted in 100 μL elution buffer [10 mM Tris-HCl (pH 8.5)], and fragmented to a target peak size of 350 bp by sonication using a Covaris S220 ultrasonicator. End repair was performed using the NEBNext Ultra II End-Repair/dA-Tailing Module following the manufacturer's instructions. 0.5X/1.0X two-sided Ampure XP bead size selection was performed, and the end repair product was eluted in 42 μL nuclease-free water. Next, the ligation mix was assembled as: 42 μL end repair product, 50 μL Quick Ligase Reaction Buffer (New England BioLabs, Cat. B2200S), 6 μL 15 μM xGen CS adapter (IDT, Cat. 1080799), and 2 μL Quick Ligase (New England BioLabs, Cat. M2200S). Ligation was performed at 20°C for 20 min, followed by 1X Ampure XP bead cleanup. A qPCR was performed to estimate the complexity. For each PCR tube, we aliquoted the purified ligation product corresponding to 1,000~2,000X human genome complexity. Next, we added 5 μL 10 μM P5_index5_Truseq5 primer (AATGATACGGCGACCACCGAGATCTACAC[8-base-index]ACACTCTTTCCCTACACGACGCTCTTCCGATCT), 5 μL 10 μM P7_index7_Truseq7 primer (CAAGCAGAAGACGGCATACGAGAT[8-base-index]GTGACTGGAGTTCAG ACGTGTGCTCTTCCGATCT), 25 μL NEBNext Ultra II Q5 master mix (New England BioLabs, Cat. M0544X), and nuclease-free water to bring the volume to 50 μL. PCR was performed as: 98°C for 30 s, 6 cycles of [98°C for 10 s, 73°C for 75 s], 73°C for 5 min. The PCR product was purified by 0.9X Ampure XP beads, resulting in the Illumina Truseq library. Based on our experience, 2 μg input DNA was sufficient to generate 24 independent libraries with 1,000~2,000X complexity each. The Illumina Truseq library was converted to the Ultima Genomics-compatible library using the UG Library Amplification Kit v3.0 (Ultima Genomics) following the manufacturer's instructions. For each tumor sample, we split the 24 independent libraries into two sequening runs on the Ultima Genomics UG100 platform, with a targeted sequencing depth of 25~30X genome coverage per library, equivalent to 600~720X genome coverage per sample.

Germline heterozygous variant calling from Illumina bulk WGS

The 3' Nextera adapter sequences were trimmed from paired-end sequencing reads using cutadapt v3.4 [1] with the argument -a CTGTCTCTTATACACATCT -A CTGTCTCTTATACACATCT -m 50. The reads were then mapped to the hg19 reference genome using bwa-mem v0.7.13-r1126 [2]. PCR duplicates were marked using samtools v1.3.1 [3] markdup command, and the germline variant was called using samtools mpileup -q 50 -Q 20 followed by bcftools call. We only kept the heterozygous single nucleotide variants with a calling quality score higher than 50. Tandem regions, centromere regions, and homopolymer regions were filtered out.

Somatic mutation calling from scNanoSeq

The 3-base UMI at the beginning of both reads was extracted using a previously published Python script [4]. Next, 3' Truseq adapter sequences were trimmed using cutadapt v3.4 with the argument -a AGATCGGAAGAGCACACGTCTGAACTCCAGTCA -A AGATCGGAAGAGCGTCGTGTAGGGAAAGAGTGT -m 50. The reads were mapped to the hg19 reference genome using bwa-mem v0.7.13-r1126. Appropriately mapped read pairs with mapping quality no less than 50 were kept, and the resulting bam file was split by UMI. For each split, raw variant calling was performed by samtools mpileup followed by bcftools call -mv. All called variants were retained regardless of the variant-calling quality score. Tandem regions, centromere regions, and homopolymer regions were filtered out, resulting in a list of raw variants.

A custom Python script was used to traverse each raw variant, a variant would be called a mutation if it fulfilled the three criteria: 1) covered in both paired-read directions, 2) at least two reads for each direction with sequencing quality no less than 30, and 3) all of the q>=30 bases support the variant base. By dividing the number of called germline heterozygous mutations (without deduplication of the same mutations called from different original DNA fragments) by the total number of germline heterozygous mutations, we can determine the mutation detection efficiency. A mutation would be called a somatic mutation if 1) the locus was covered by at least 10 reads in the matched normal bulk WGS with sequencing quality >=30, but zero read supports the variant call, and 2) the variant is not annotated in the dbSNP database. Somatic mutation load was calculated by dividing the number of somatic mutations detected by the mutation detection efficiency.

Quantification of strand dropout rate

The strand dropout rate was calculated based on the number of germline heterozygous mutations detected in both strands or only one single strand of the template DNA fragments. We denote the strand dropout rate as *sd*, total germline heterozygous mutation as *T*, and the allele coverage rate as *ac*. Then, we expected to call $T*ac*(1-sd)^2$ germline heterozygous mutations in both strands, while $T*ac*2*sd*(1-sd)$ germline heterozygous mutations are detected in only one strand. Therefore, the ratio between germline heterozygous mutations detected in both strands versus only one strand $Ratio = \frac{Germline\_hets\_both\_strands}{Germline\_hets\_single\_strand} = \frac{1-sd}{2*sd}$. Based on the observed ratio, the strand dropout can be calculated as $sd = \frac{1}{2*Ratio+1}$.

Mutation-based tumor evolution tree construction

For each breast cancer sample, we generated a list of genomic loci that covers the somatic mutations detected in at least one single cell. Using this list and the bam files of all single cells, we performed multi-sample samtools mpileup followed by bcftools call to generate a genotype likelihood matrix. The matched normal bulk sample was included as well to locate the root of the tree. The matrix was passed into CellPhy v0.9.2 [5] for tumor evolution tree inference. The tumor lineage was determined based on the existence of a long root edge in the tree structure.

Estimation of stem length mutation density based on CellPhy phylogenetic tree

For each single tumor cell, we first quantified its distance to the root ($D_{total}$) on the phylogenetic tree inferred by CellPhy. Next, on the same phylogenetic tree, we quantified the distance between the single cell to the closest node on the next layer of the ACDC ($D_{post\_clonal\_sweep}$). It is worth noting that since one or two early branchings from the ACDC were frequently observesed in the tree structures, direct calculation of the single cell distance to the ACDC node itself is prone to the variation of the branching point we sampled. Therefore, to alleviate this variation, we quantified the $D_{post\_clonal\_sweep}$ as the minimal distance between single cells to the second layer of nodes followed by the ACDC. As a result, the stem length mutation load is calculated as single cell mutation load multiplied by (1- $D_{post\_clonal\_sweep}/D_{total}$). The stem length mutation load of each tumor sample is calculated as an average of single tumor cell-based estimations.

Cell proportion test for neutral evolution on the CellPhy phylogenetic tree

For each sample with no fewer than 40 single tumor cells profiled, we examine the major nodes (with >=8 cells) followed by the ACDC on the CellPhy inferred phylogenetic tree. For each major node, we quantify the cell numbers in both daughter lineages defined by the node. The unbalance ratio of the node is defined as the cell number in the major daughter lineage divided by the cell number in the minor daughter lineage, and its 95% confidence interval is derived based on the binomial proportion test implemented by the prop.test function in R.

#### CNV analysis for scNanoSeq

Starting with the single-cell bam files, we marked the PCR duplicated reads using Picard v3.0.0 MarkDuplicates. Duplicated reads were then removed by samtools rmdup, and the deduplicated bam files were converted to bed files with bedtools v.2.26.0 bamtobed [6] and gzip compressed. The bed files were passed to the local version of Ginkgo master-build d7c7790 [7] for single-cell CNV calling at the 500k bin-size resolution. An estimated ploidy file based on FANS was provided for global CNV correction in Ginkgo. All the remaining parameters were set as default.

To cluster the single cells based on the CNV profile, we generated a pairwise distance matrix, where the distance between two cells was calculated as (1 – spearman correlation of copy number profiles). The distance matrix was then used for hierarchical clustering with ward.D2 linkage. To infer the time order, single cells were sorted by between-leaf or between-branch similarity based on the optimal leaf ordering method [8] implemented in R package seriation v.1.4.2. Fraction of genome altered is defined by the fraction of regions whose copy numbers deviated from the major copy number of the cell (chrY is excluded).

#### Mutational signature analysis

Mutational signature analysis was performed using the MutationalPatterns R package v3.4.1 [9] following its online tutorial.

#### Variant calling from Ultima ultra-deep bulk WGS

The 5' Truseq Read 1 adapter was trimmed using cutadapt v3.4 with the argument -g CTACACGACGCTCTTCCGATCT --discard-untrimmed --minimum-length 100, and 3' Truseq Read 2 adapter was trimmed using cutadapt v3.4 with the argument -a AGATCGGAAGAGCACACGTCTGAACTCCAGTCAC --minimum-length 50. After trimming, the UMI appeared to be the 3 bases in the beginning of the read, and the fastq file for each independent library was split into 64 files based on UMI. For each split, the reads were mapped to the hg19 reference genome using bwa-mem v0.7.13-r1126. PCR duplicates were removed by samtools markdup with argument -s --mode=s -r. The resulting deduplicated bam files were merged together, and variants were called using a low allele frequency variant caller LoFreq v2.1.5 [10] with argument --min-mq 60 --min-alt-bq 30. Variants called with strand bias score higher than 8 were filtered out. A variant is considered somatic if 1) the locus was covered by at least 10 reads in the matched normal bulk WGS with sequencing quality >=30, but zero read supports the variant call, and 2) the variant is not annotated in the dbSNP database.

#### Quantification of stem length mutation density based on ultra-deep bulk WGS

Based on the single tumor cell CNV patterns, we extract the genome regions with consensus copy number, where at least 80% of single cells demonstrated the same copy number. The consensus tetraploid (4N), triploid (3N), and diploid (2N) regions were kept for analysis if the total length exceeds $1 \times 10^8$ bp. Within each of these regions, we first plot out the variant allele frequency (VAF) distribution. Next, we manually select a VAF lower cutoff (normally ranging from 0.1 to 0.2) to exclude subclonal mutations, and fit the remaining VAF distribution to the sum of two or three normal distributions using the curve_fit function in the scipy Python library. The fitted normal distributions correspond to the mutations on different numbers of alleles. The final stem length mutation density is calculated as a weighted average of the quantification based on consensus 4N, 3N, and 2N regions. For tumor content estimation, we denote the tumor cell fraction as $f_t$. As a result, in the region of ploidy = $p$, the observed VAF is expected to be $\frac{f_t}{p \times f_t + 2 \times (1 - f_t)}$ for the peak corredponding to one allele. Therefore, the tumor cell fraction could be derived as $f_t = \frac{2 \times VAF_{1allele\_peak}}{1 + (2 - p) \times VAF_{1allele\_peak}}$.

Ultra-deep bulk WGS-guided single-cell evolutionary path analysis

For each single cell, we listed the genomic loci where somatic mutations were detected. Next, we used LoFreq call --plp-summary-only to pileup these loci in the matched ultra-deep bulk WGS bam file. Using a custom python code, we filtered out the bases with sequencing quality q<30, and counted the frequency of the four bases A, C, G, and T in the bulk WGS, respectively. To minimize the sequencing platform-dependent coverage bias, we excluded the loci with fewer than 100X depth at q30 sequencing quality. As a result, we defined the bulk VAF distribution of single cell-detected mutations (without deduplication of the same mutations called from different molecules) as the evolutionary path fingerprint of this exact single cell. Next, for each pairs of single cells, we performed Kolmogorov-Smirnov test on their evolutionary path fingerprints.

Simulation datasets for evolutionary path analysis

Tumor evolutionary simulation is performed using python 3.10. We defined a class for simulated single cells that independently divide and accumulate mutations. Each cell contains information including the mother of the cell, the time of generation, accumulated mutations, and tumor cell fitness (duplication rate). For each newly generated cell, its mutations were composed of inherited mutations from the parental cells and newly acquired mutations. The latter one were hypothesized to accumulate through a Poisson process. Newly acquired mutations are randomly sampled based on hg19 genomic size for the purpose of potential co-occurrence of mutations, but with no assumptions on the effect of mutation loci with cancer cell fitness. Tumor evolution is modeled in two different manners: branched evolution followed by neutral evolution, or continuous branched evolution. In the neutral mode, duplication rate of all cells is set to be constant one, regardless of number of mutations accumulated. In branched mode, after each division occurred, fitness of the two daughter cells is separately calculated based on the mutations accumulated, followed by a subsequent modification adjusting the fitness to magnify the imbalance of the branching process. The effect of mutations on cell fitness is determined by a concerted effect of neutral drift, cancer driver mutations, and tumor suppressor mutations. After the effects of all mutations are sampled for each cell, a collective effect of cellular fitness is calculated as a linear combination of all existing mutations. It is worth noting that if the same mutations occur in different cells, the same fitness effect is assigned to all the cells carrying the same mutation. The evolution simulation is initiated with three starting cells, namely, one normal cell with no somatic mutations (as the root) and two early branched tumor cells carrying 2,000 – 3,000 pre-lesion mutations. Evolution simulation is run for a user-determined period (experimental time for cells to evolve) to allow a full development of each experiment. 50 tumor cells were randomly selected for the WGS-guided single-cell evolutionary path analysis framework. We followed a Poission distribution to randomly sample an average of 125, 250, 500, 1000, or 2000 mutations per simulated tumor cell and then performed the pair-wise Kolmogorov-Smirnov test with their VAF in the bulk sample.

Estimation of whole tumor mutation burden

We first determined the number of mutations that is single cell-specific. The averaged terminal branch length $L$ derived from the CellPhy-inferred phylogenetic tree (corrected for the systematic bias) was taken as an approxiamation of the single cell-unique somatic mutation load. Next, based on the effectively exponential expansion model in the dominant clone, the effective tumor cell generation number $n$ is derived from log2 of the total tumor cell count generated at onset of neutral evolution. The tumor cell count is calculated as tumor weight multiplied by tumor cell content, with the estimation that 1 mg tissue corresponds to 1 million cells. The tumor cell content was estimated by FANS. With the averaged terminal branch length $L$ in the most recent clade (where effectively neutral evolution is reached), and effective cell generation number *n*, the effective mutation rate per cell doubling can be calculated as *L/n*. Therefore, the whole tumor mutation burden can be derived as $\sum_{i=0}^{n} 2^i \times \frac{L}{n} = (2^{n+1} - 1) \times \frac{L}{n}$ plus the stem mutation load identified.

### Supplementary Note 1. Mutation-based phylogenetic tree inference algorithms

The commonly used mutation-based phylogenetic tree inference algorithms mainly adopt a maximum likelihood approach, including SCITE[11], SiFit[12], and the more recent development of CellPhy[5]. It is worth noting that all three algorithms are based on a diploid assumption. Although SCITE and SiFit utilize an input genotype matrix in an absence/presence mode, both methods follow a Bayesian model to convert such observed absence/presence information of a variant into the probability of homozygous reference, heterozygous, or homozygous non-reference genotypes on the diploid assumption. The newly developed CellPhy algorithm is essentially on the same diploid assumption basis as SCITE or SiFit, but it implemented a genotype transition matrix of 16 diploid states and optimized the estimation of allele dropout rate to improve the robustness of inferred phylogenies and evolution distance.

Nevertheless, both earlier methods SCITE and SiFit indeed have been successfully performed on the tumor datasets in their original manuscripts, in which the lineage of single diploid cells and the lineages of single aneuploidy cells could be separated. With this observation, we reason that CellPhy could also be applied to aneuploidy cells to distinguish the single cell lineage structure. For aneuploidy genomes, we may concatenate the multiple copies of the same allele into one, thereby generating a pseudo-diploid genome. For example, in a 4N genomic locus with 3 reference alleles and 1 alternative allele (3REF+1ALT), it can be regarded as a diploid locus, where the detection rate of REF and ALT is unbalanced and the ALT detection rate is proportional to its mutation density. When we feed such a pseudo-diploid genotype that is reflective of mutation density, CellPhy algorithm essentially infer a phylogenetic tree based on mutation density. In our study, the accuracy of the CellPhy inferred phylogenetic tree is cross-validated by two independent methods, including CNV tree and bulk VAF evolutionary path fingerprint analysis.

**Supplementary Note 2. Estimating the number of loss-of-function mutations**

For each amino acid, consider the possibility that leads to stop codons (TAG, TAA, TGA) if one mutation occurs: the following sequences can mutate into a stop codon (A/C/G)AG, (A/C/G)AA, (A/C/G)GA, TCG, TGC, TTG, TTC with 1/3 chances, TGG has 2/3 chances to mutate into a stop codon. In total, 17/61 of codons can mutate into a stop codon. The human protein length distribution peaks at 320 amino acids[13]. Even if we only focus on the first 1/10 of a protein, which is equivalent to 32 amino acids, in total $32 \times 16/61 \times 1 + 32 \times 1/61 \times 2 = 9$ point mutations can lead to a premature stop codon, resulting in a truncated loss-of-function protein. With at least tens of genes being identified to be capable of endocrine therapy tolerance [14], we expect more than one hundred mutations that can boost drug tolerance in tumor cells.

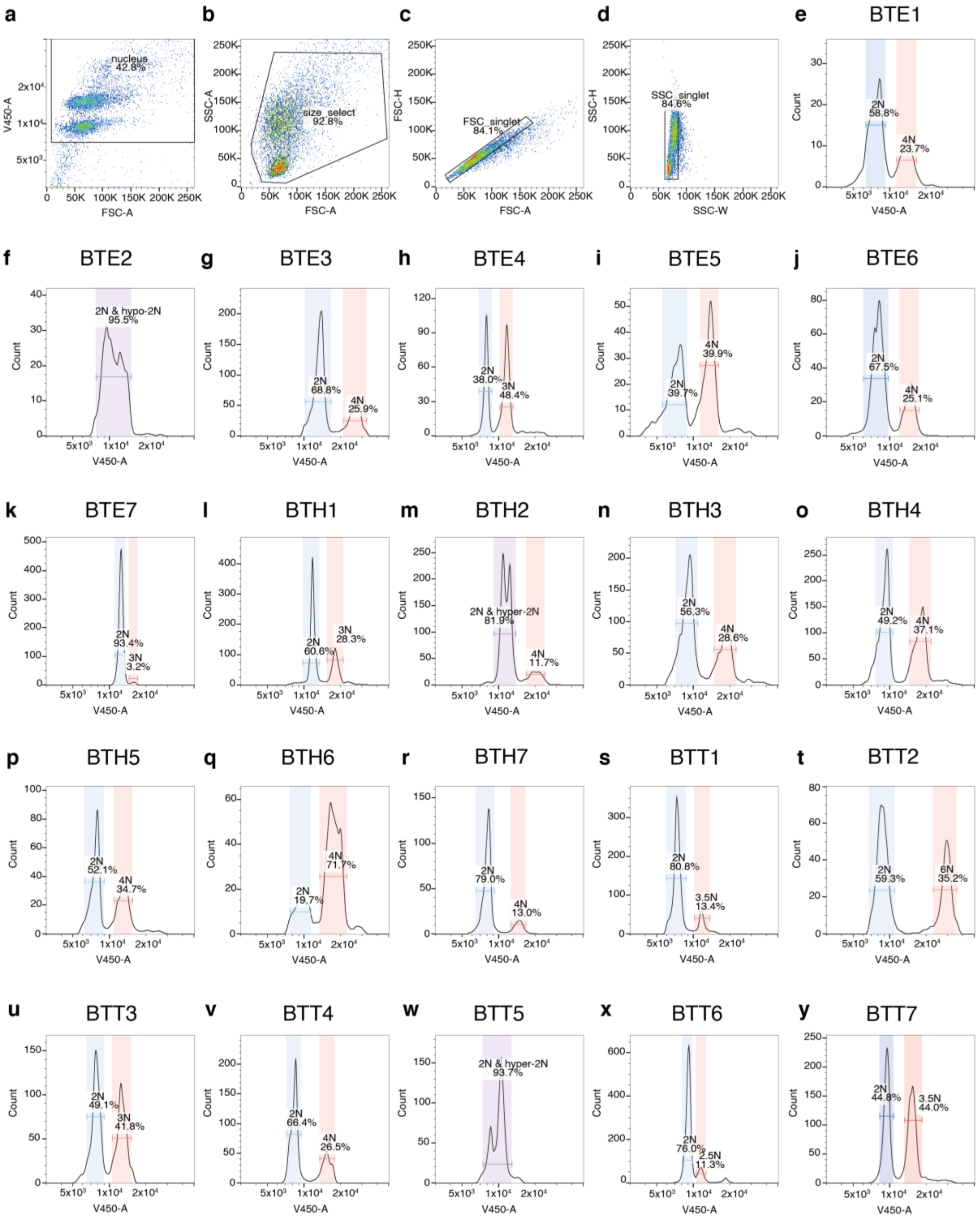
a
nucleus
42.8%
V450-A
FSC-A
b
size_select
92.8%
SSC-A
FSC-A
c
FSC_singlet
84.1%
FSC-H
FSC-A
d
SSC_singlet
84.6%
SSC-H
SSC-W
e BTE1
2N 58.8%
4N 23.7%
f BTE2
2N & hypo-2N 95.5%
g BTE3
2N 68.8%
4N 25.9%
h BTE4
2N 38.0%
3N 48.4%
i BTE5
2N 39.7%
4N 39.9%
j BTE6
2N 67.5%
4N 25.1%
k BTE7
2N 93.4%
3N 3.2%
l BTH1
2N 60.6%
3N 28.3%
m BTH2
2N & hyper-2N 81.9%
4N 11.7%
n BTH3
2N 56.3%
4N 28.6%
o BTH4
2N 49.2%
4N 37.1%
p BTH5
2N 52.1%
4N 34.7%
q BTH6
2N 19.7%
4N 71.7%
r BTH7
2N 79.0%
4N 13.0%
s BTT1
2N 80.8%
3.5N 13.4%
t BTT2
2N 59.3%
6N 35.2%
u BTT3
2N 49.1%
3N 41.8%
v BTT4
2N 66.4%
4N 26.5%
w BTT5
2N & hyper-2N 93.7%
x BTT6
2N 76.0%
2.5N 11.3%
y BTT7
2N 44.8%
3.5N 44.0%
Count
V450-A

**Supplementary Fig. 1. FANS gating strategy to isolate single candidate tumor nuclei and single diploid normal nuclei.** (**a**)-(**d**) Identification of single nuclei based on the forward scatter (FSC), side scatter (SSC), and Hoechst intensity (V450 channel). (**e**)-(**y**) Identification of single diploid normal nuclei and nondiploid candidate tumor nuclei based on Hoechst intensity in 21 breast cancer samples.

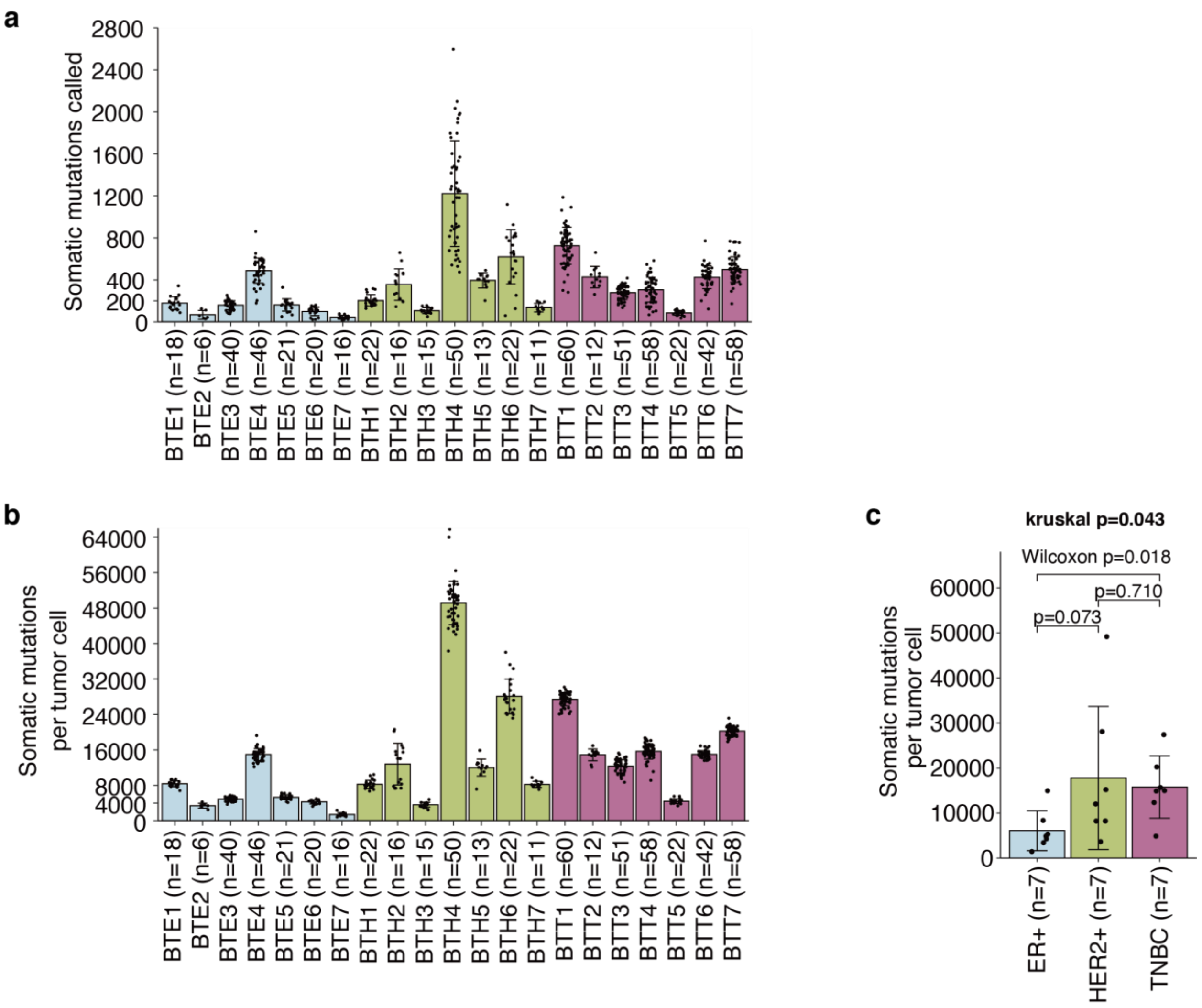


**Supplementary Fig. 2. Somatic mutation load in single tumor cells.** (**a**) Bar plot demonstrating the number of detected somatic mutations per tumor cell. (**b**) Bar plot demonstrating the estimated number of somatic mutations per tumor cell. (**c**) Bar plot demonstrating the average number of somatic mutations per tumor cell in individual tumor samples, grouped by breast cancer subtypes. Kruskal-Wallis test was performed among the three breast cancer subtypes. A two-sided Wilcoxon ranksum test was performed for pairwise comparison. For panels (**a**)-(**c**), error bars indicate standard deviation

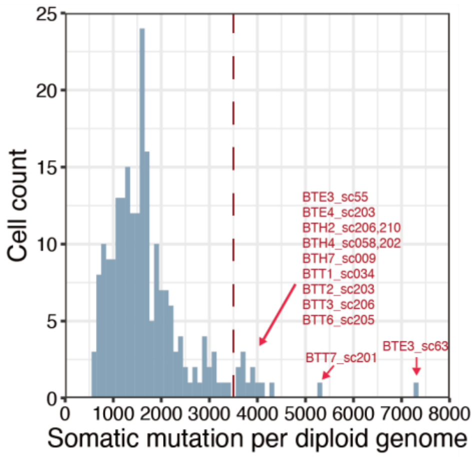


**Supplementary Fig. 3. Histogram showing the distribution of somatic mutation load in normal cells.** 13 out of 217 normal lineage cells were identified as hypermutators with somatic mutation load higher than 3,500 (dashed line).

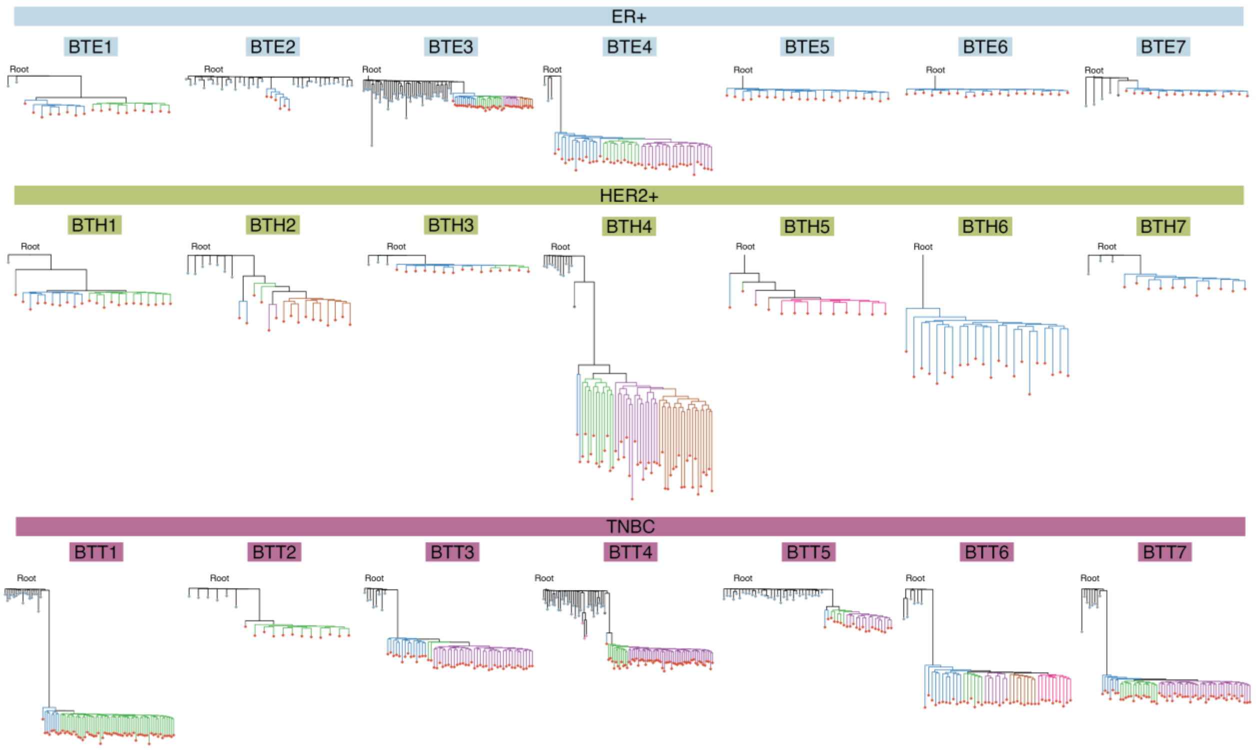


**Supplementary Fig. 4. The mutation-based phylogenetic trees of 21 breast cancer samples.** Length scaled to mutation load across all 21 samples.

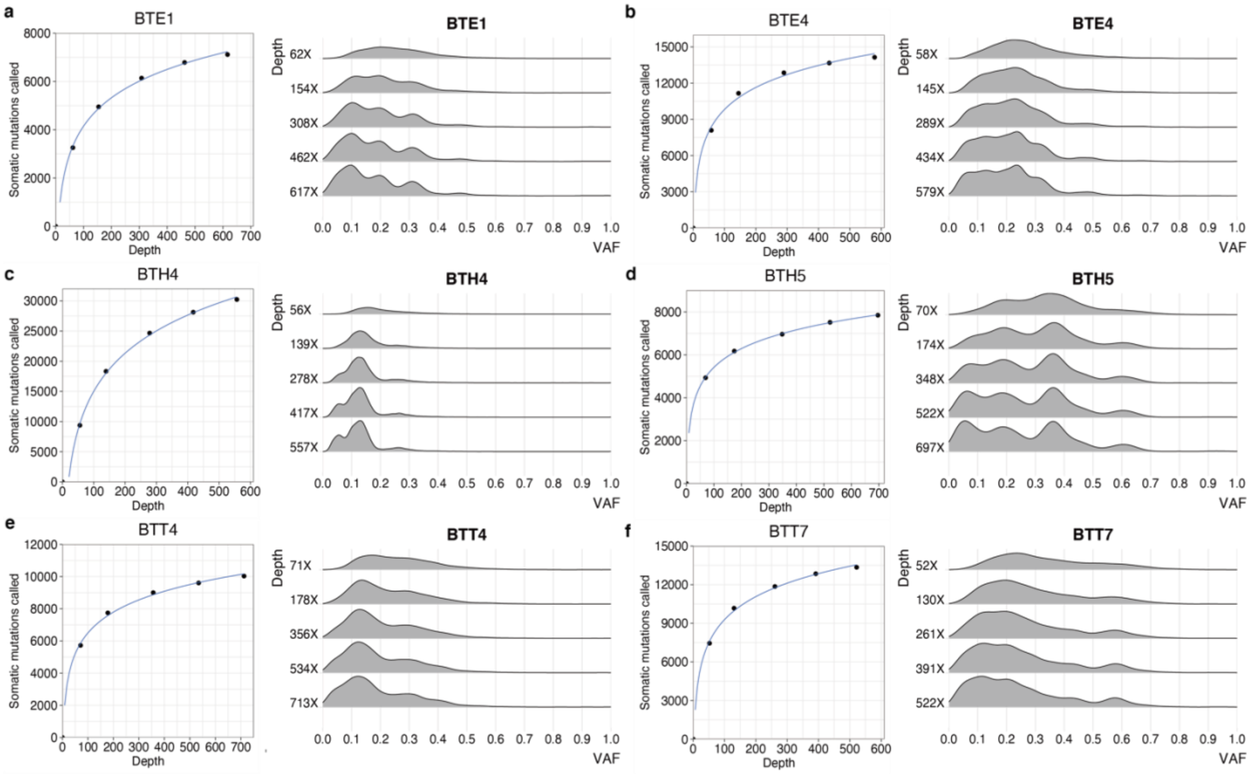


**Supplementary Fig. 5. Somatic mutations detected in ultra-deep tumor bulk WGS.** The tumor bulk WGS is subsampled at different depth (mean coverage per locus with sequencing quality >=30) for somatic mutation calling using LoFreq. (**a-f**) For different tumor samples (BTE1, BTE4, BTH4, BTH5, BTT4, and BTT7), the scatter plots demonstrated the saturation curve showing the number of called somatic mutations at different depths, and the ridgeplots display the variant allele frequency (VAF) distribution of called somatic mutations.

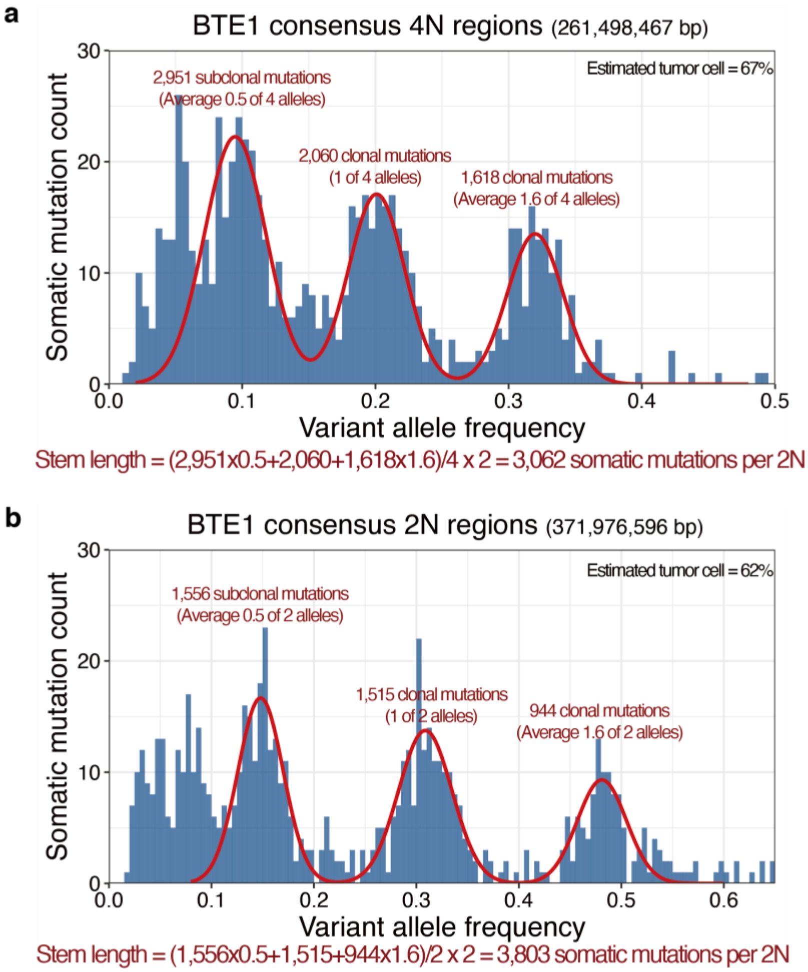


**Supplementary Fig. 6. Quantification of stem length mutation density in BTE1 ultra-deep bulk WGS.** (**a**) Quantification of clonal and subclonal mutations in consensus tetraploid regions. (**b**) Quantification of clonal and subclonal mutations in consensus diploid regions.

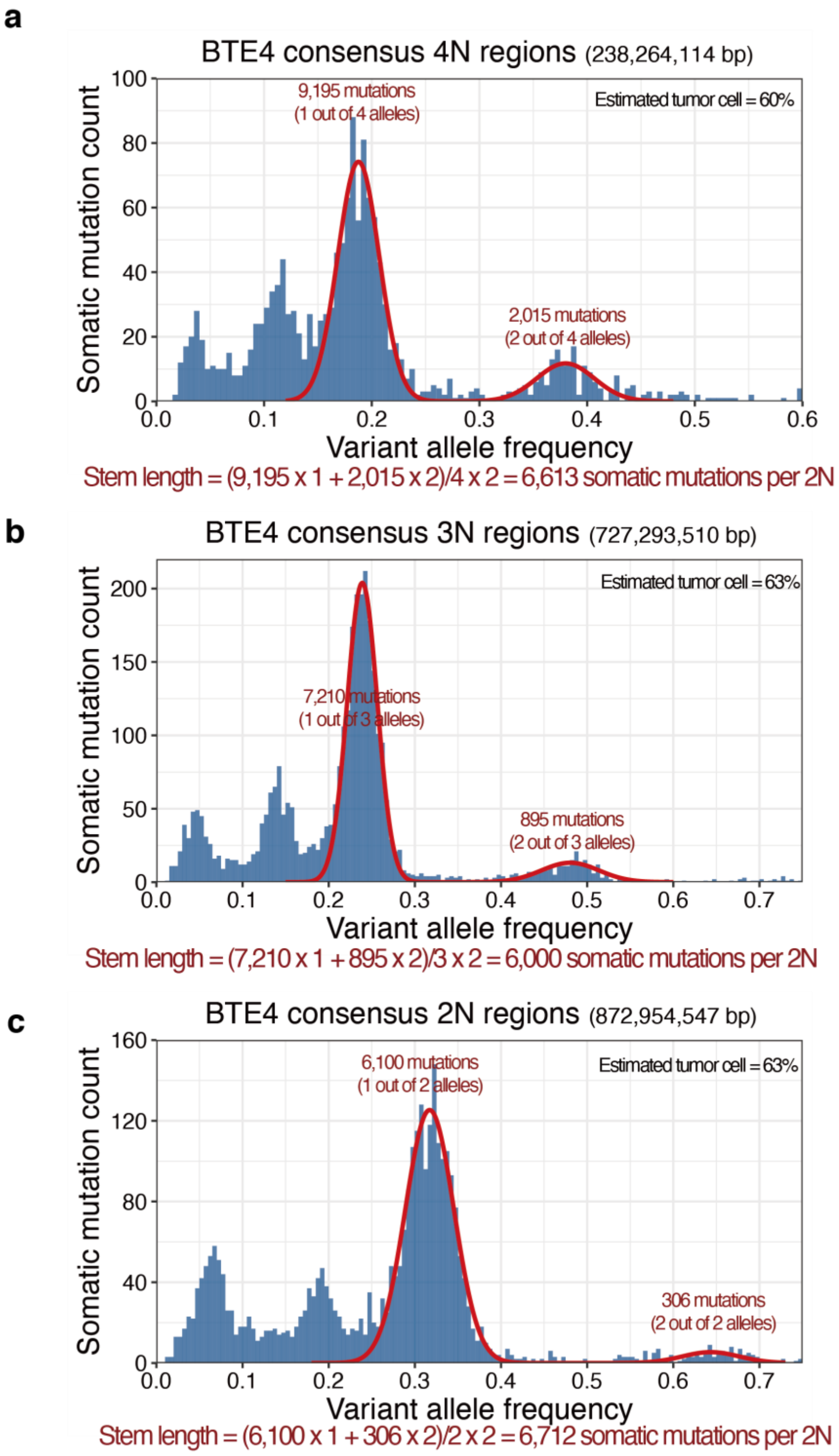


**Supplementary Fig. 7. Quantification of stem length mutation density in BTE4 ultra-deep bulk WGS.** (**a**) Quantification of clonal and subclonal mutations in consensus tetraploid regions. (**b**) Quantification of clonal and subclonal mutations in consensus triploid regions. (**c**) Quantification of clonal and subclonal mutations in consensus diploid regions.

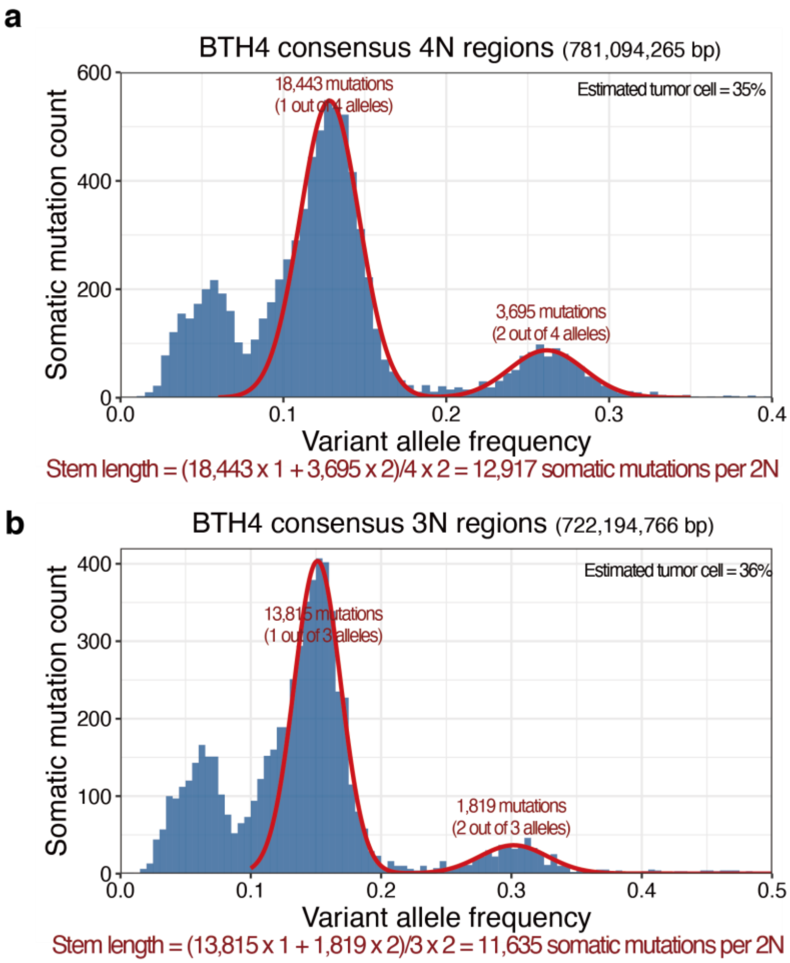


**Supplementary Fig. 8. Quantification of stem length mutation density in BTH4 ultra-deep bulk WGS.** (**a**) Quantification of clonal and subclonal mutations in consensus tetraploid regions. (**b**) Quantification of clonal and subclonal mutations in consensus triploid regions.

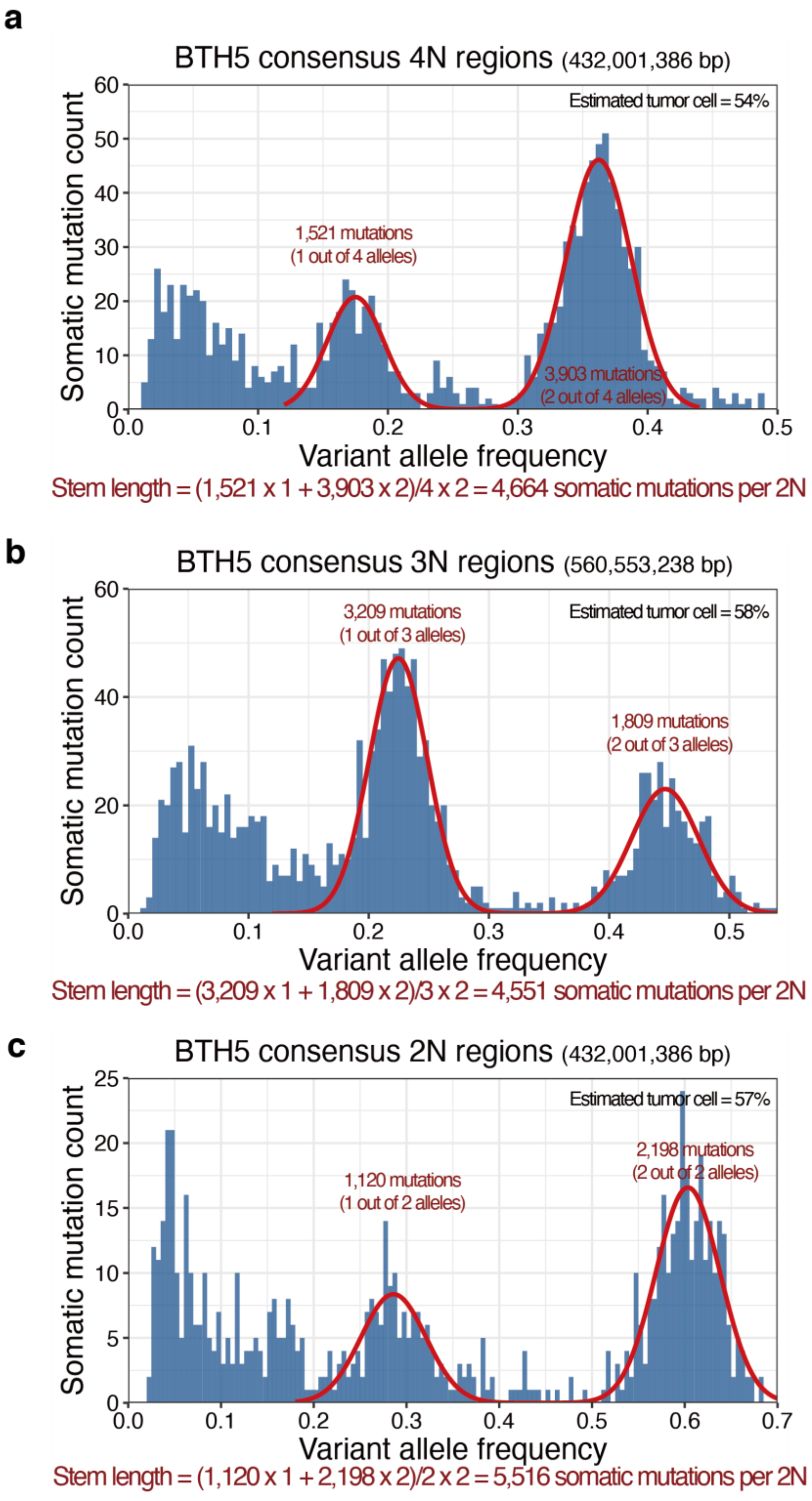


**Supplementary Fig. 9. Quantification of stem length mutation density in BTH5 ultra-deep bulk WGS.** (**a**) Quantification of clonal and subclonal mutations in consensus tetraploid regions. (**b**) Quantification of clonal and subclonal mutations in consensus triploid regions. (**c**) Quantification of clonal and subclonal mutations in consensus diploid regions.

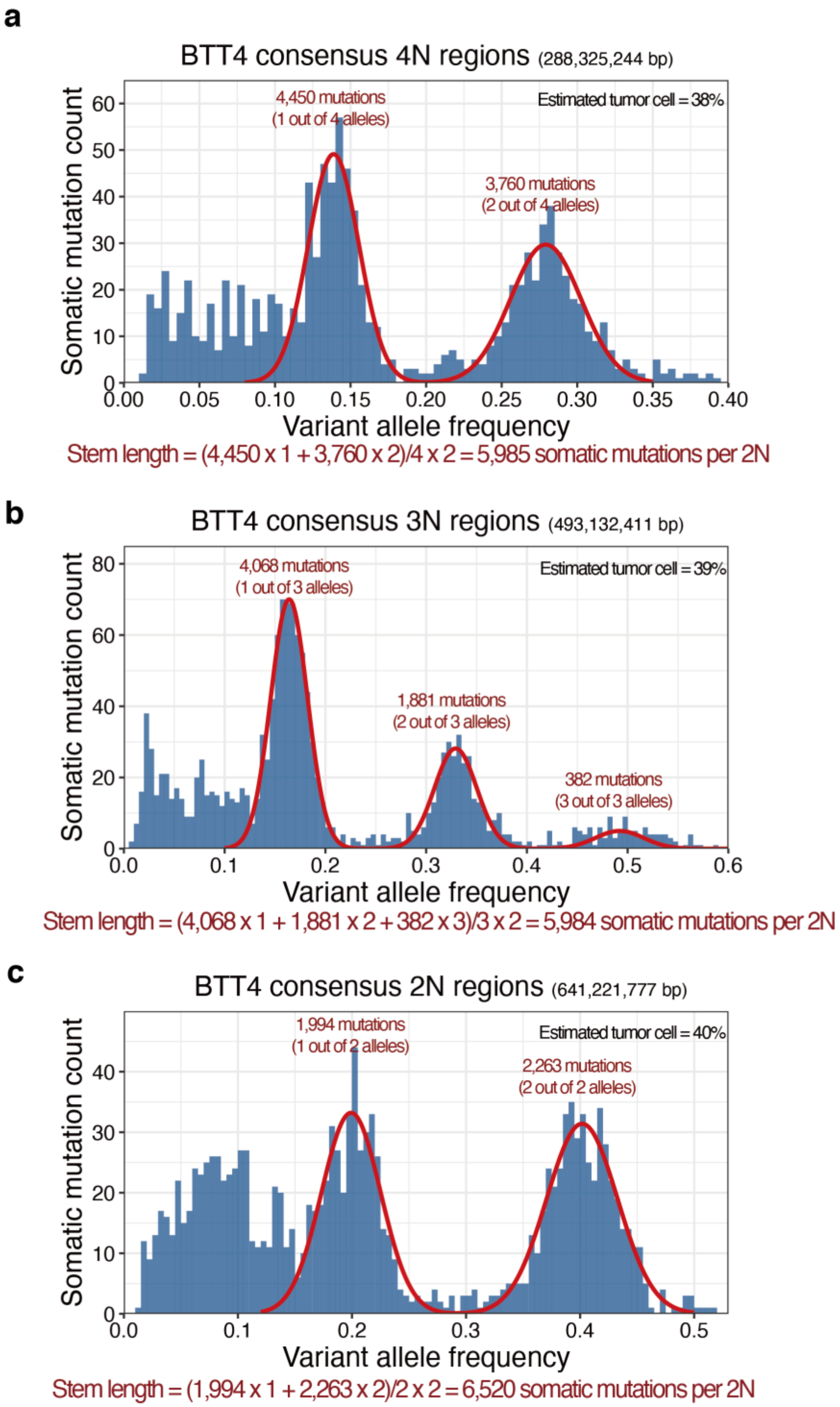


**Supplementary Fig. 10. Quantification of stem length mutation density in BTT4 ultra-deep bulk WGS.** (**a**) Quantification of clonal and subclonal mutations in consensus tetraploid regions. (**b**) Quantification of clonal and subclonal mutations in consensus triploid regions. (**c**) Quantification of clonal and subclonal mutations in consensus diploid regions.

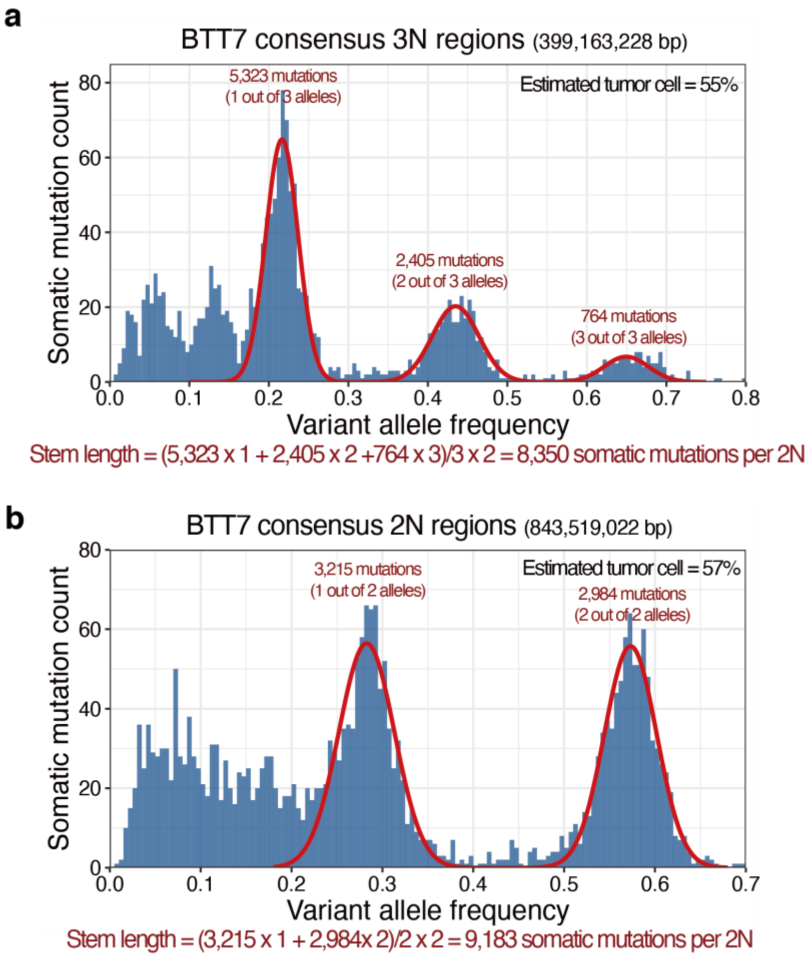


**Supplementary Fig. 11. Quantification of stem length mutation density in BTT7 ultra-deep bulk WGS.** (**a**) Quantification of clonal and subclonal mutations in consensus triploid regions. (**b**) Quantification of clonal and subclonal mutations in consensus diploid regions.

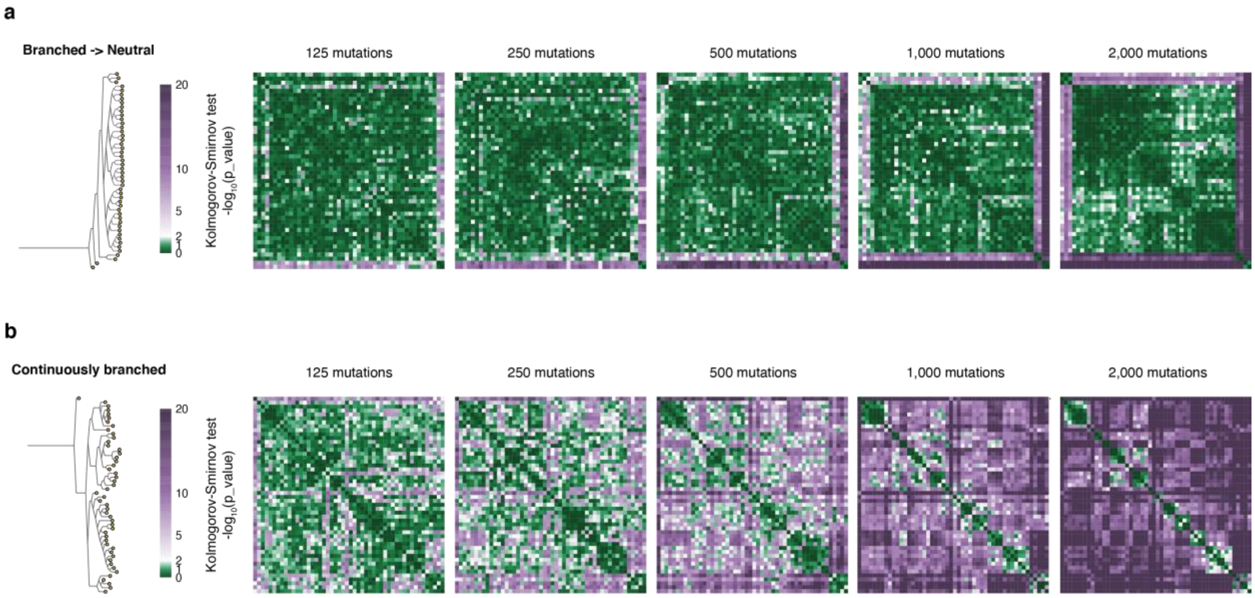


**Supplementary Fig. 12. Evaluation of the bulk-guided single-cell evolutionary path analysis framework on simulated datasets.** For each single cell, different number of mutations were randomly sampled for pairwise Kolmogorov-Smirnov test, with the pairwise Kolmogorov-Smirnov p values displayed in the heatmaps. (**a**) The simulated datasets under the assumption of branched evolution followed by neutral evolution. (**b**) The simulated datasets under the assumption of continuously branched evolution.

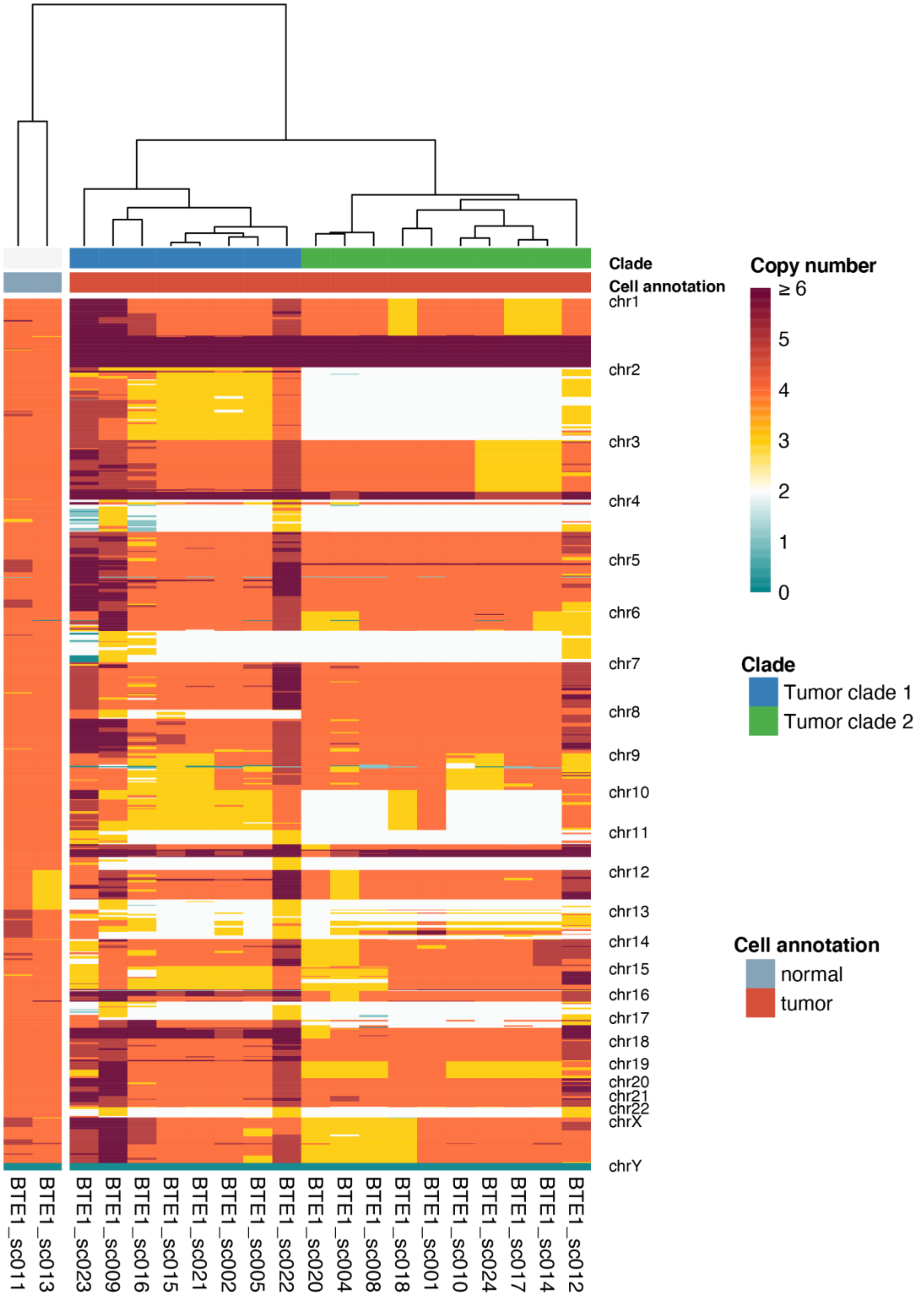


**Supplementary Fig. 13. Hierarchical clustering of CNV profiles of tumor BTE1.**

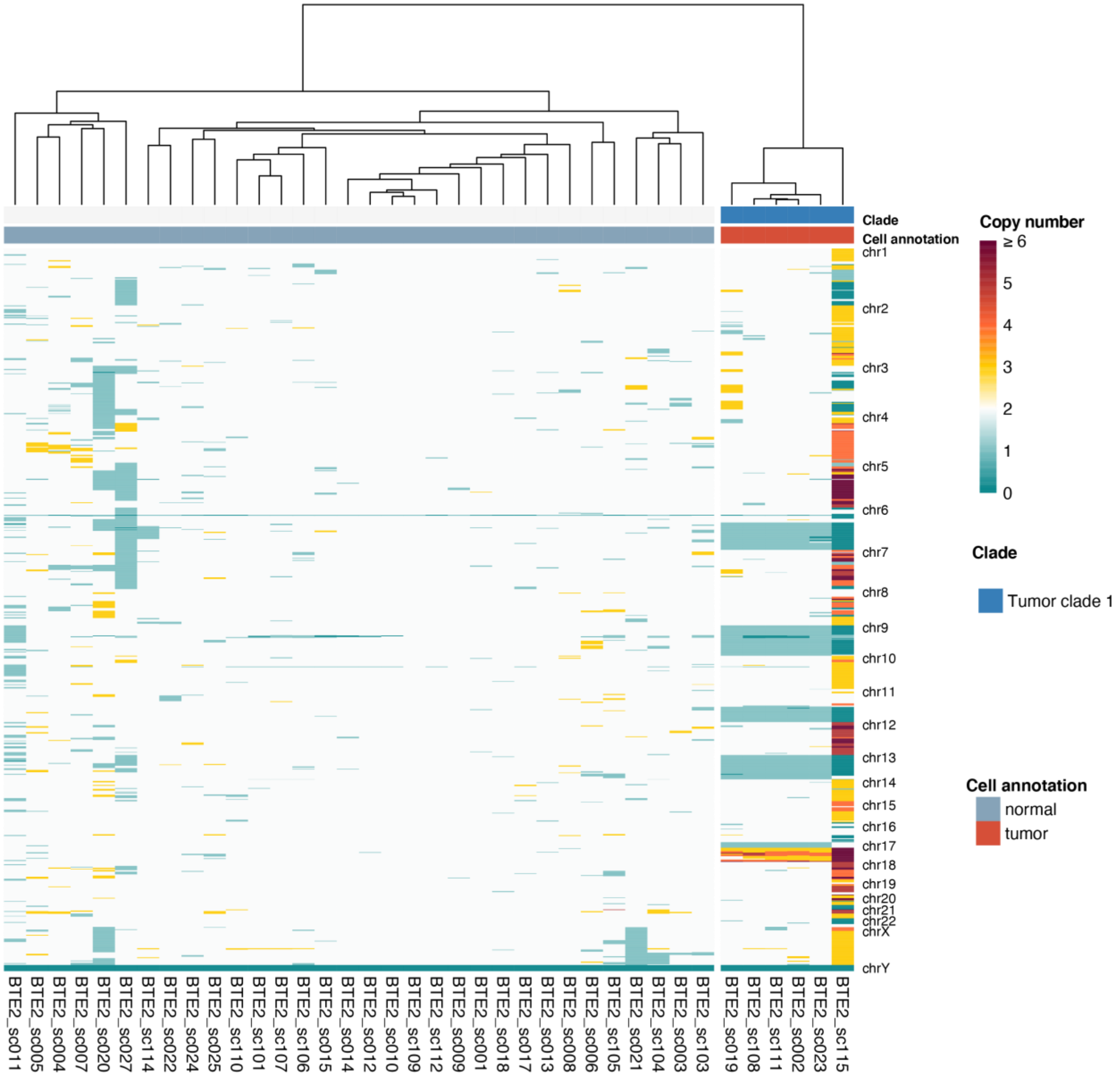


**Supplementary Fig. 14. Hierarchical clustering of CNV profiles of tumor BTE2.**

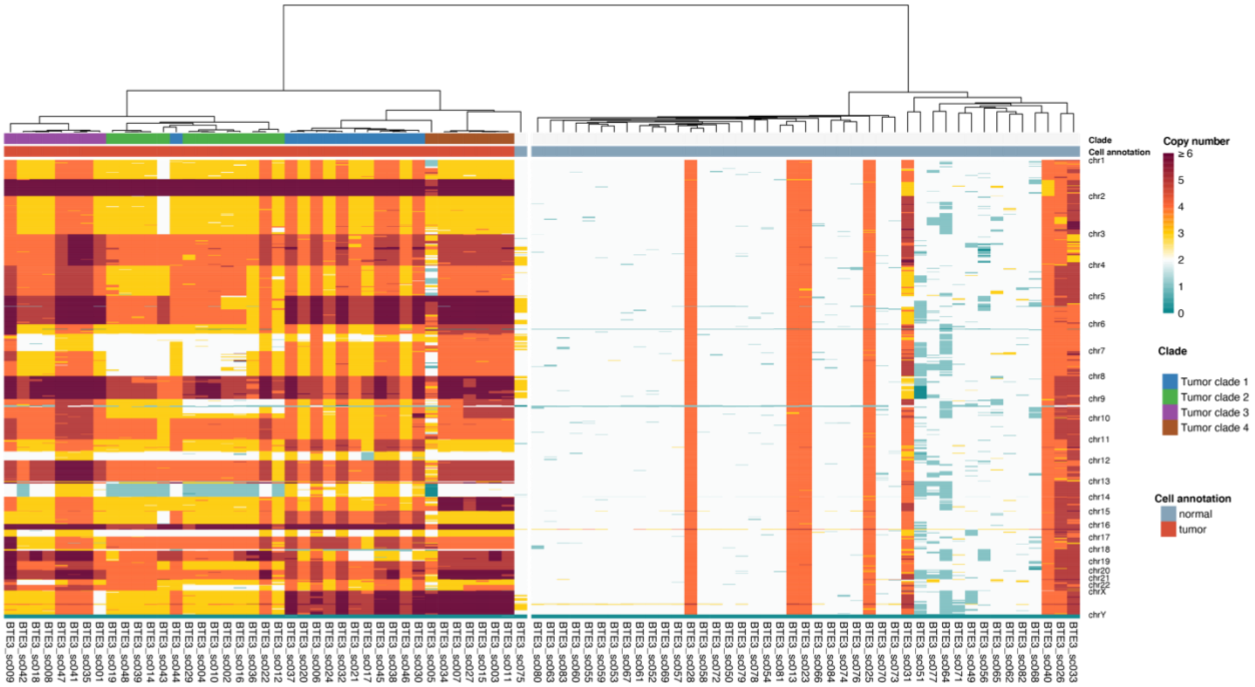


**Supplementary Fig. 15. Hierarchical clustering of CNV profiles of tumor BTE3.**

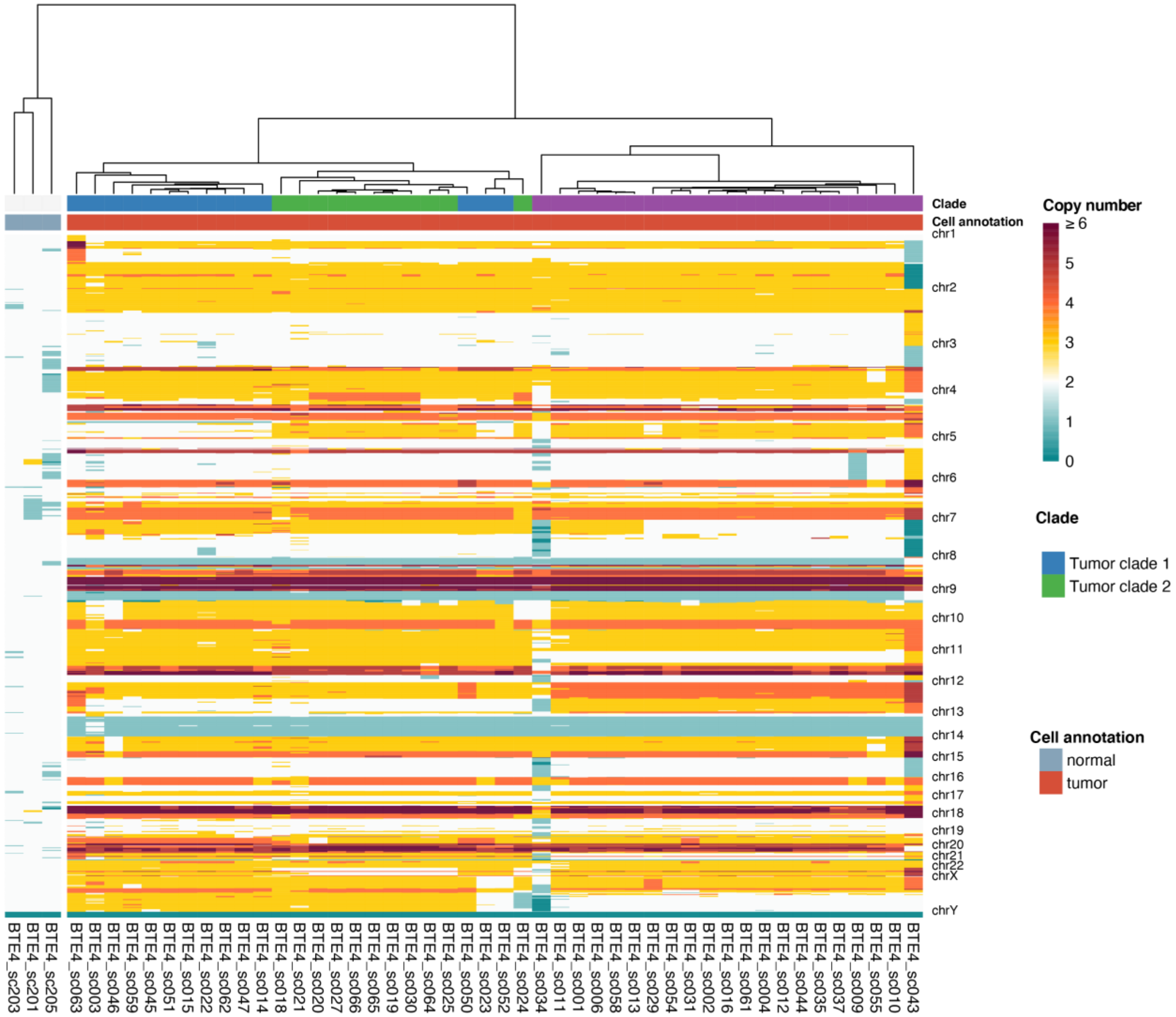


**Supplementary Fig. 16. Hierarchical clustering of CNV profiles of tumor BTE4.**

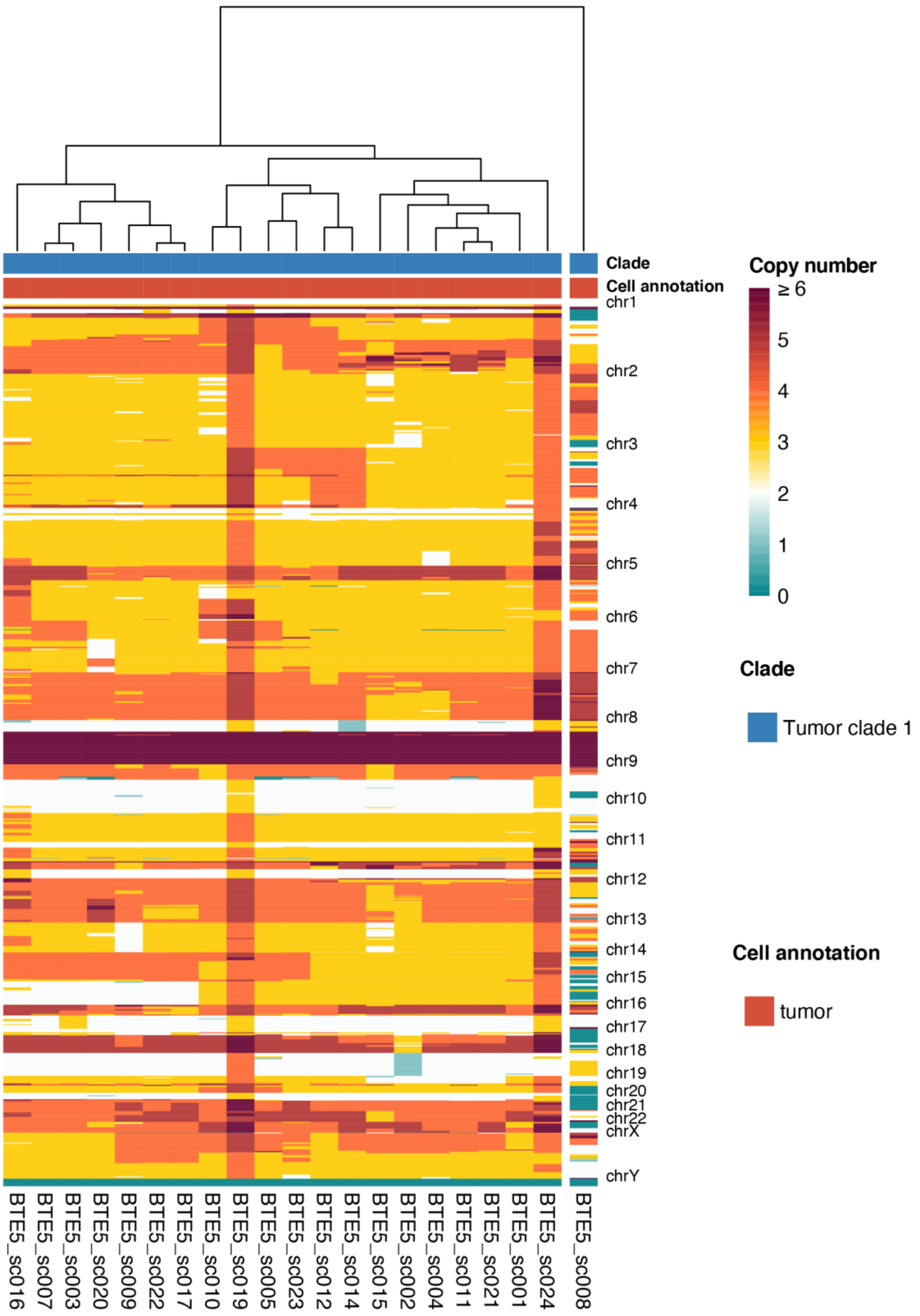


**Supplementary Fig. 17. Hierarchical clustering of CNV profiles of tumor BTE5.**

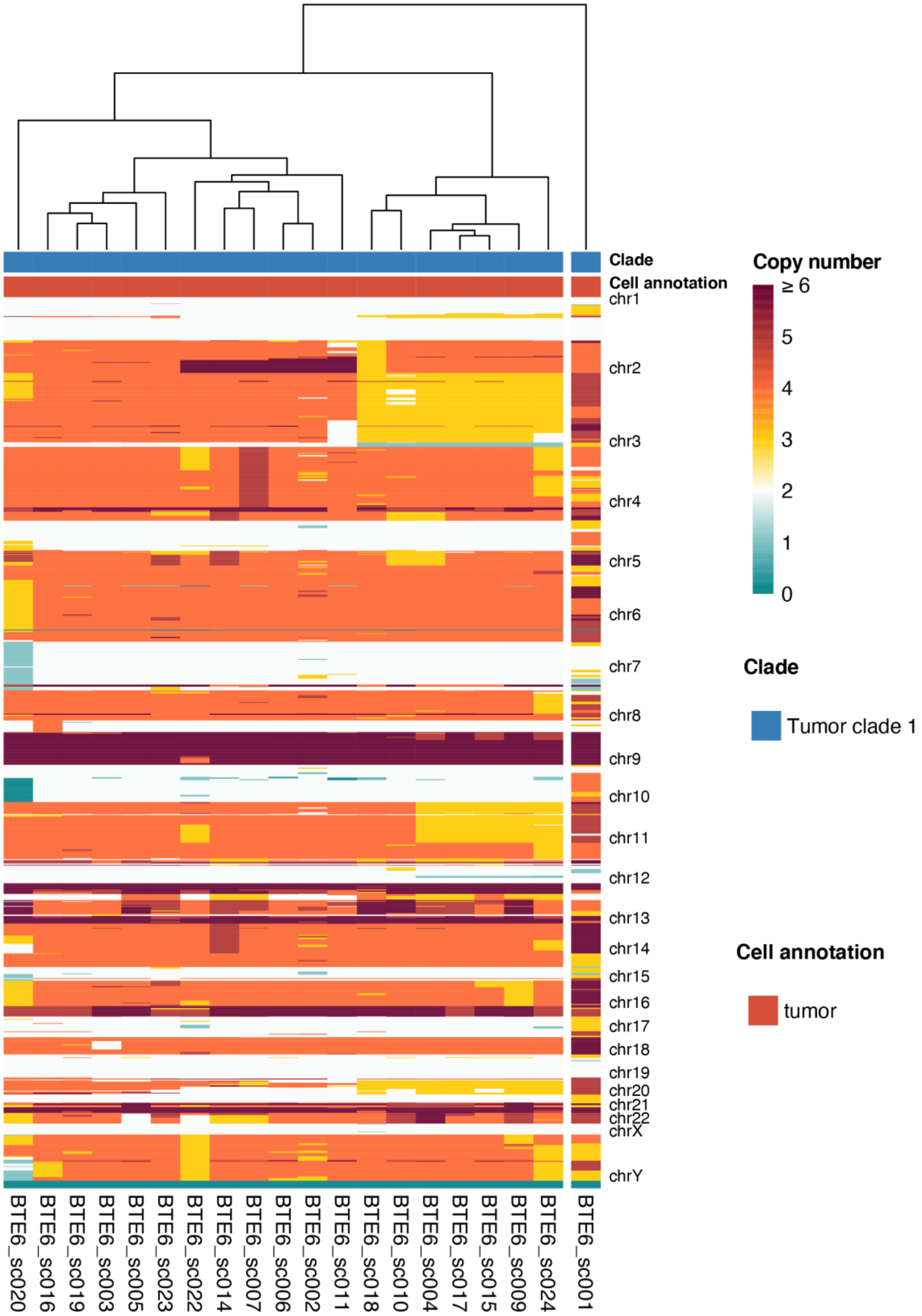

**Supplementary Fig. 18. Hierarchical clustering of CNV profiles of tumor BTE6.**

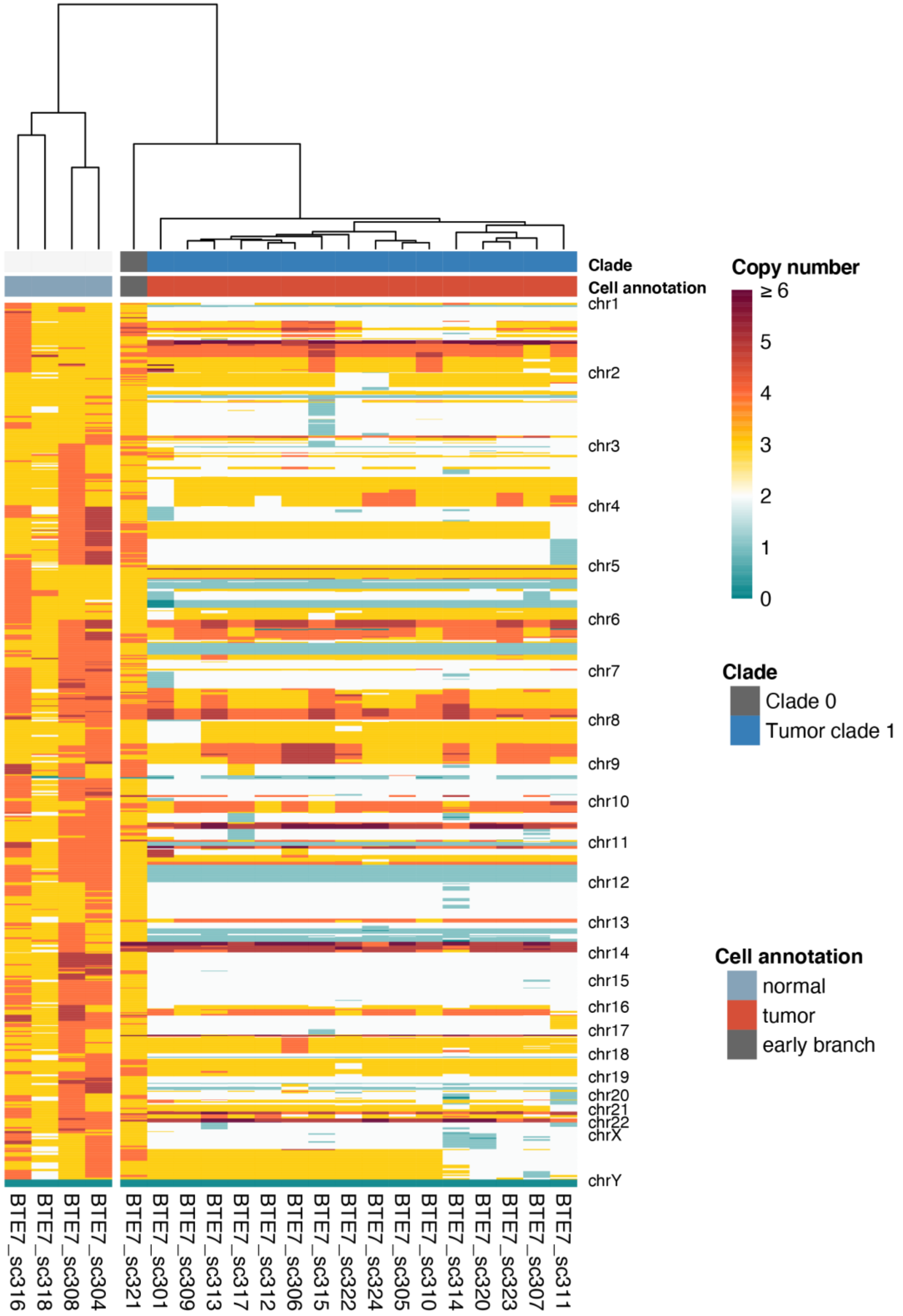


**Supplementary Fig. 19. Hierarchical clustering of CNV profiles of tumor BTE7.**

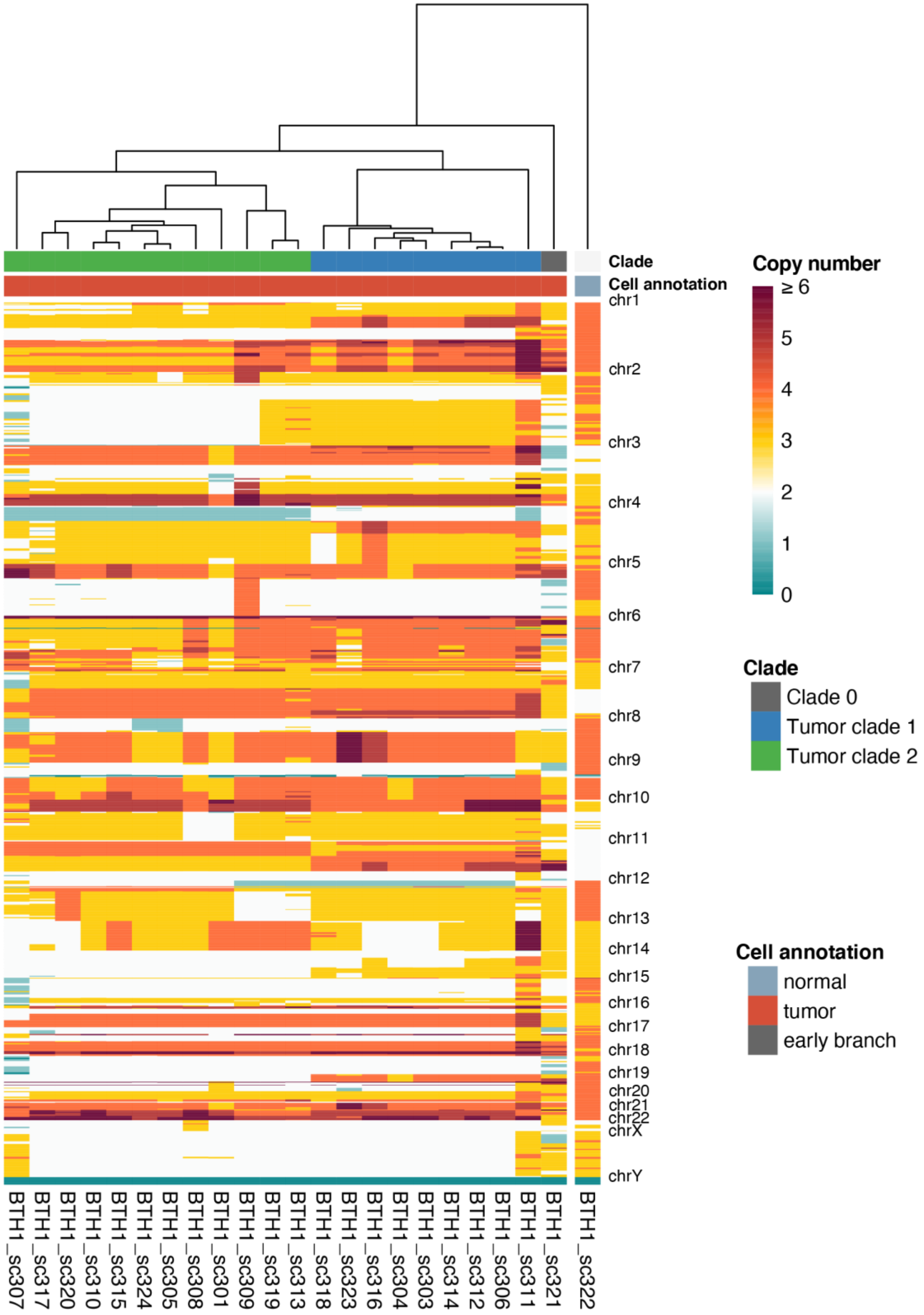


**Supplementary Fig. 20. Hierarchical clustering of CNV profiles of tumor BTH1.**

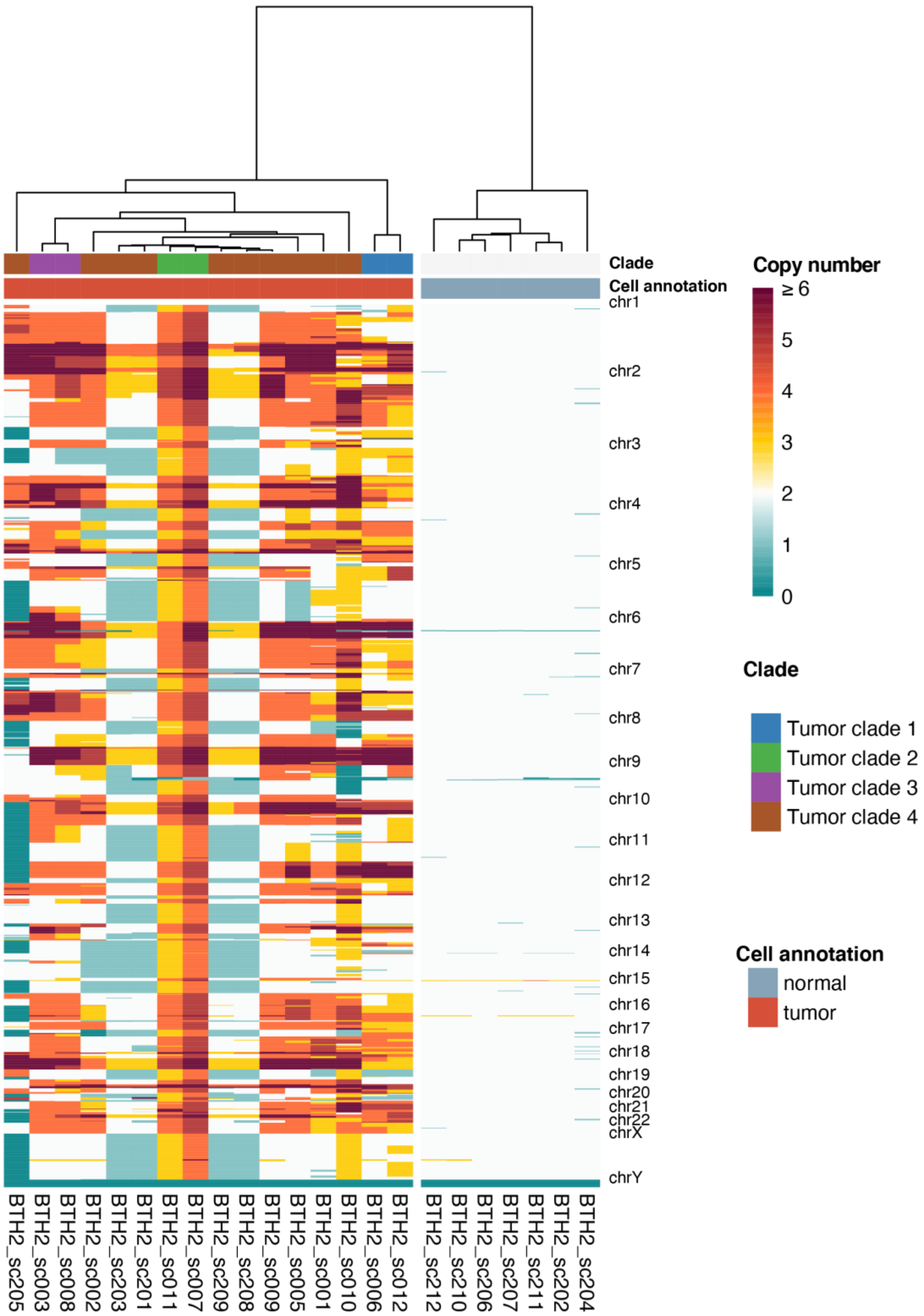


**Supplementary Fig. 21. Hierarchical clustering of CNV profiles of tumor BTH2.**

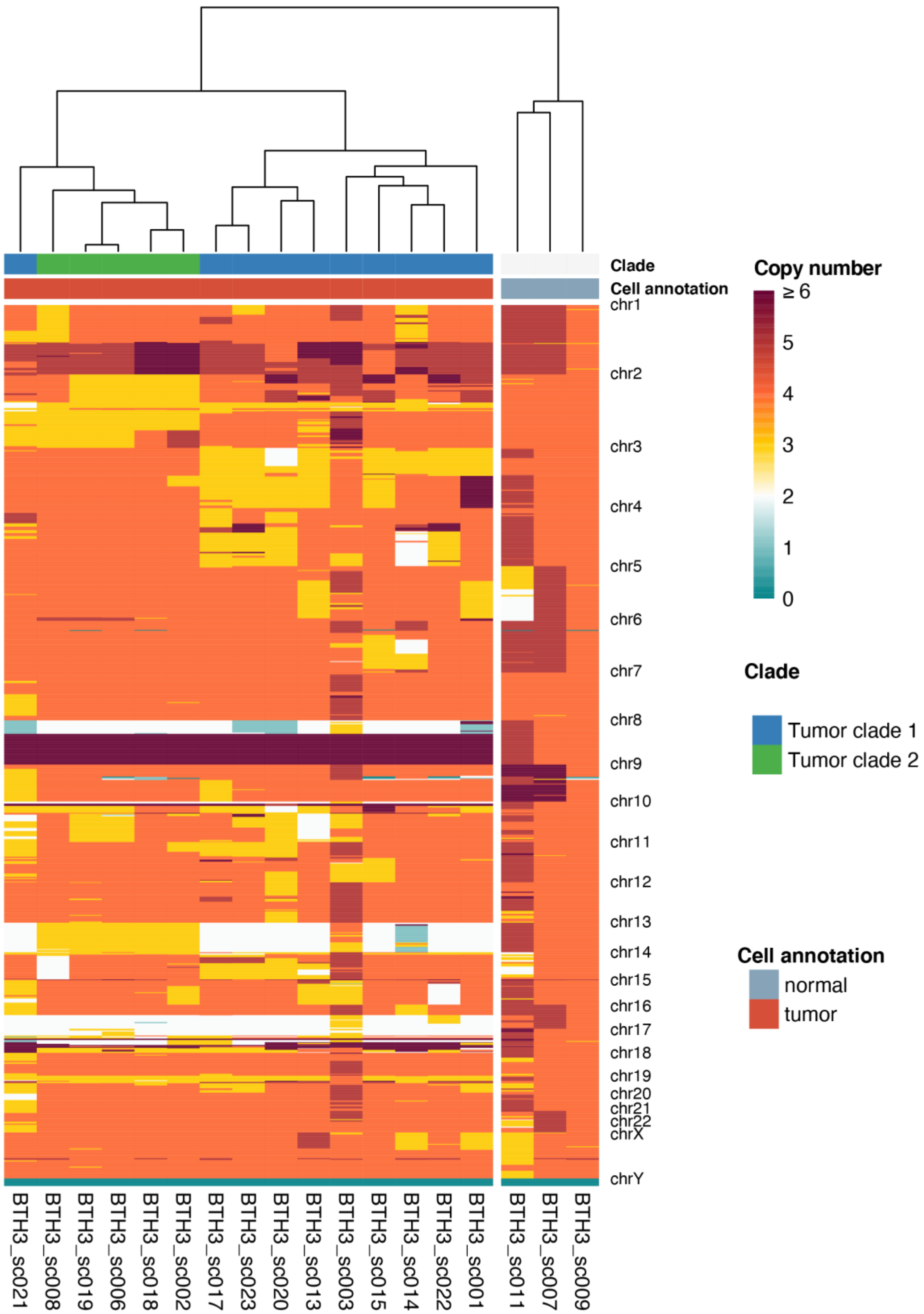


**Supplementary Fig. 22. Hierarchical clustering of CNV profiles of tumor BTH3.**

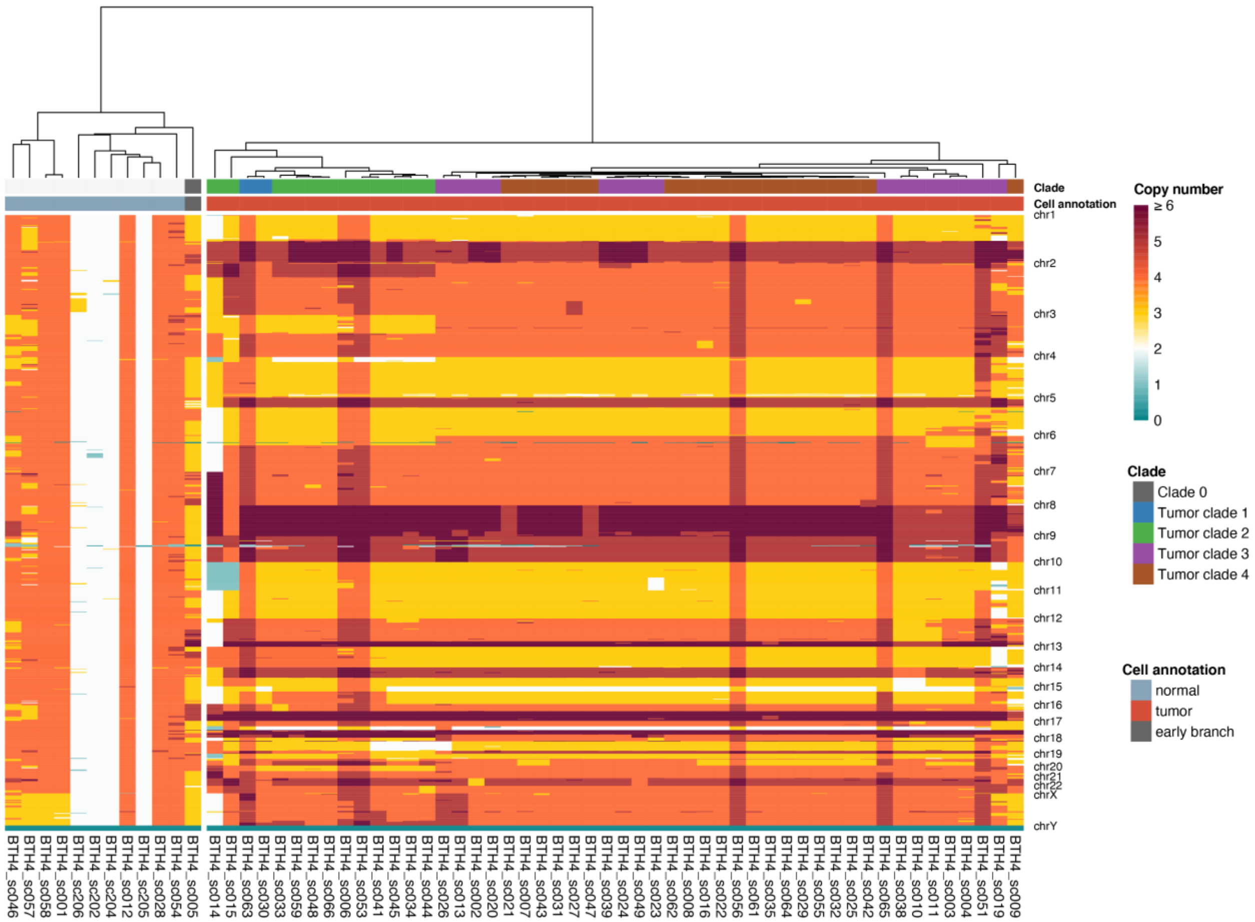


**Supplementary Fig. 23. Hierarchical clustering of CNV profiles of tumor BTH4.**

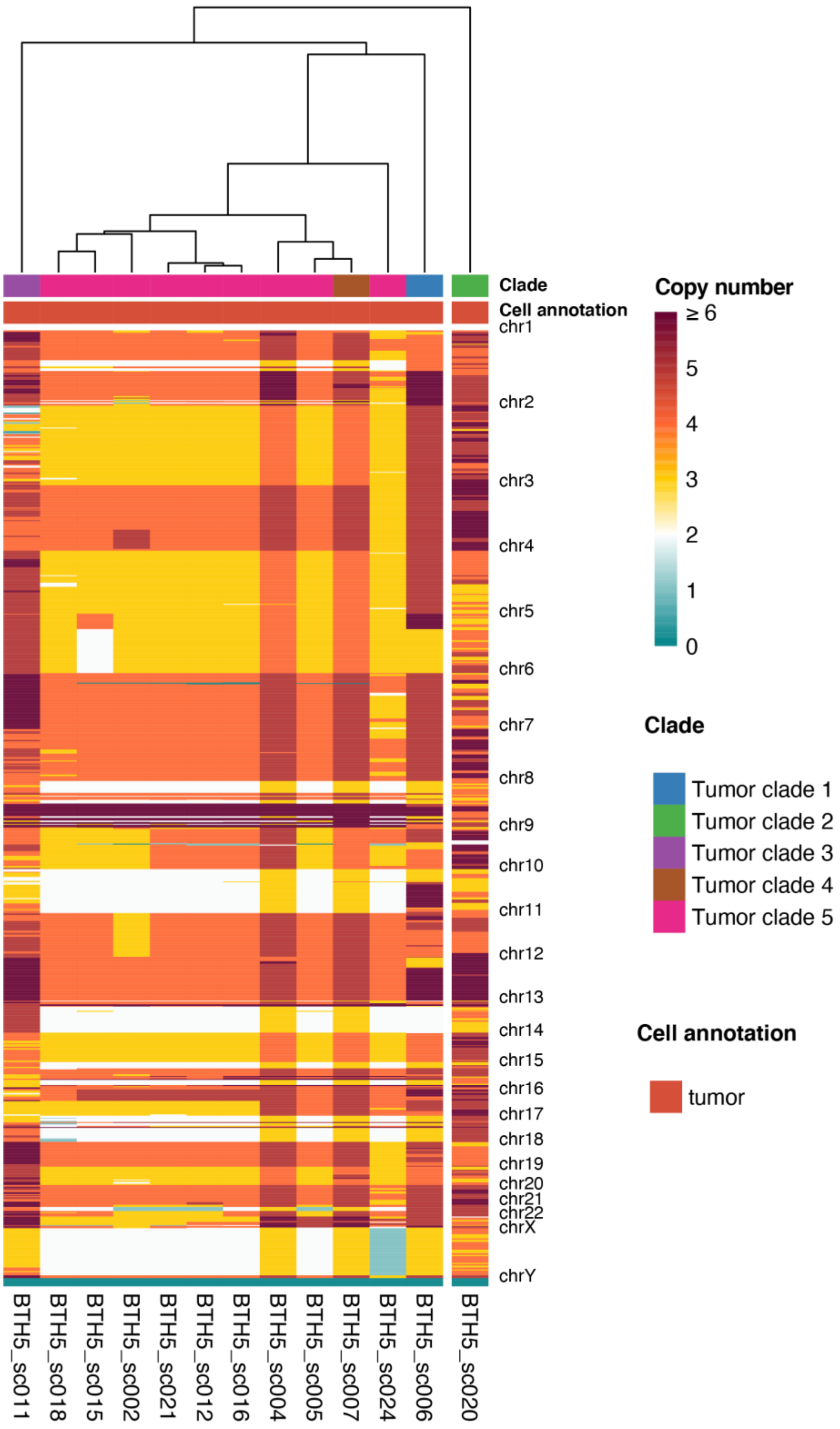


**Supplementary Fig. 24. Hierarchical clustering of CNV profiles of tumor BTH5.**

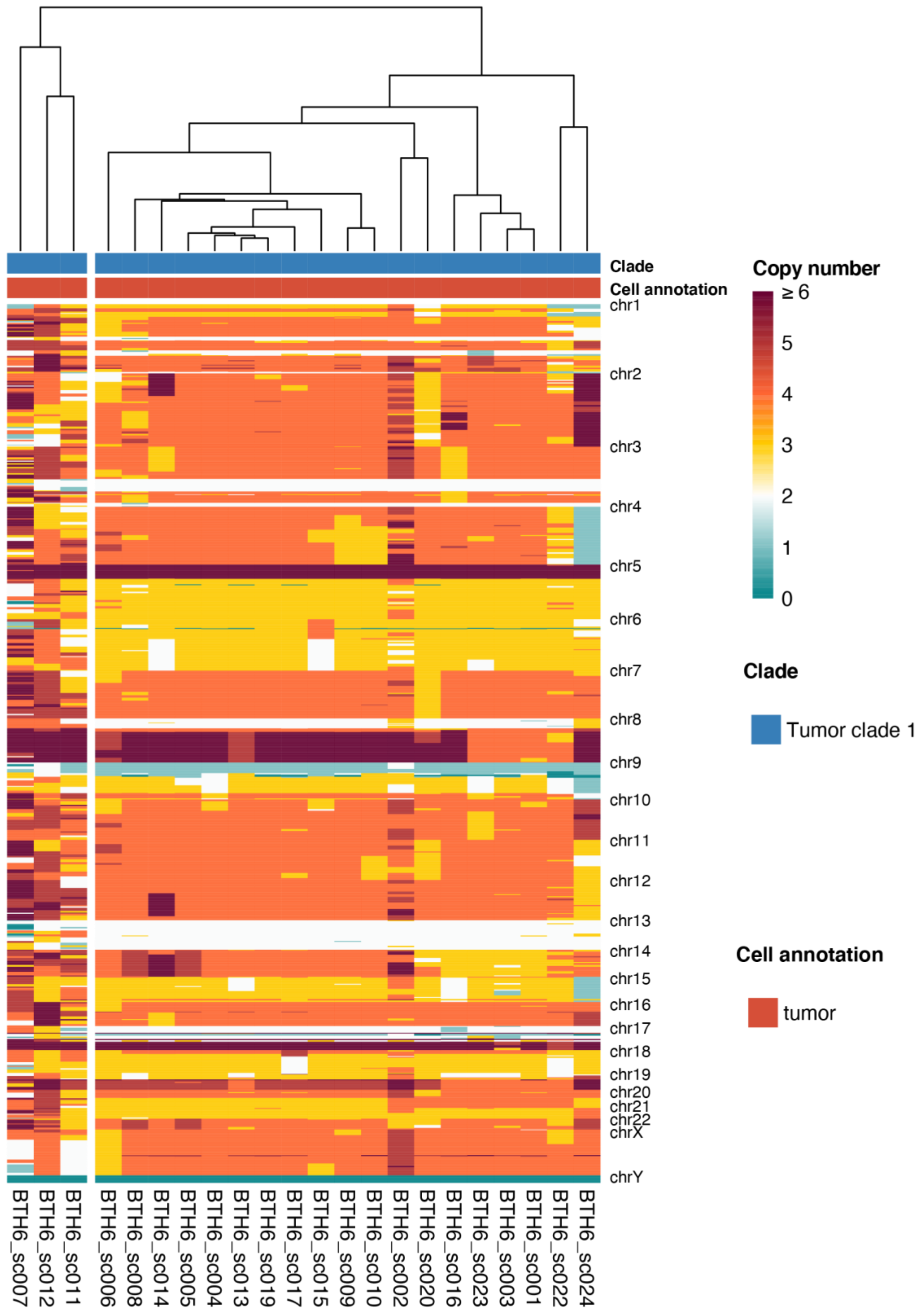


**Supplementary Fig. 25. Hierarchical clustering of CNV profiles of tumor BTH6.**

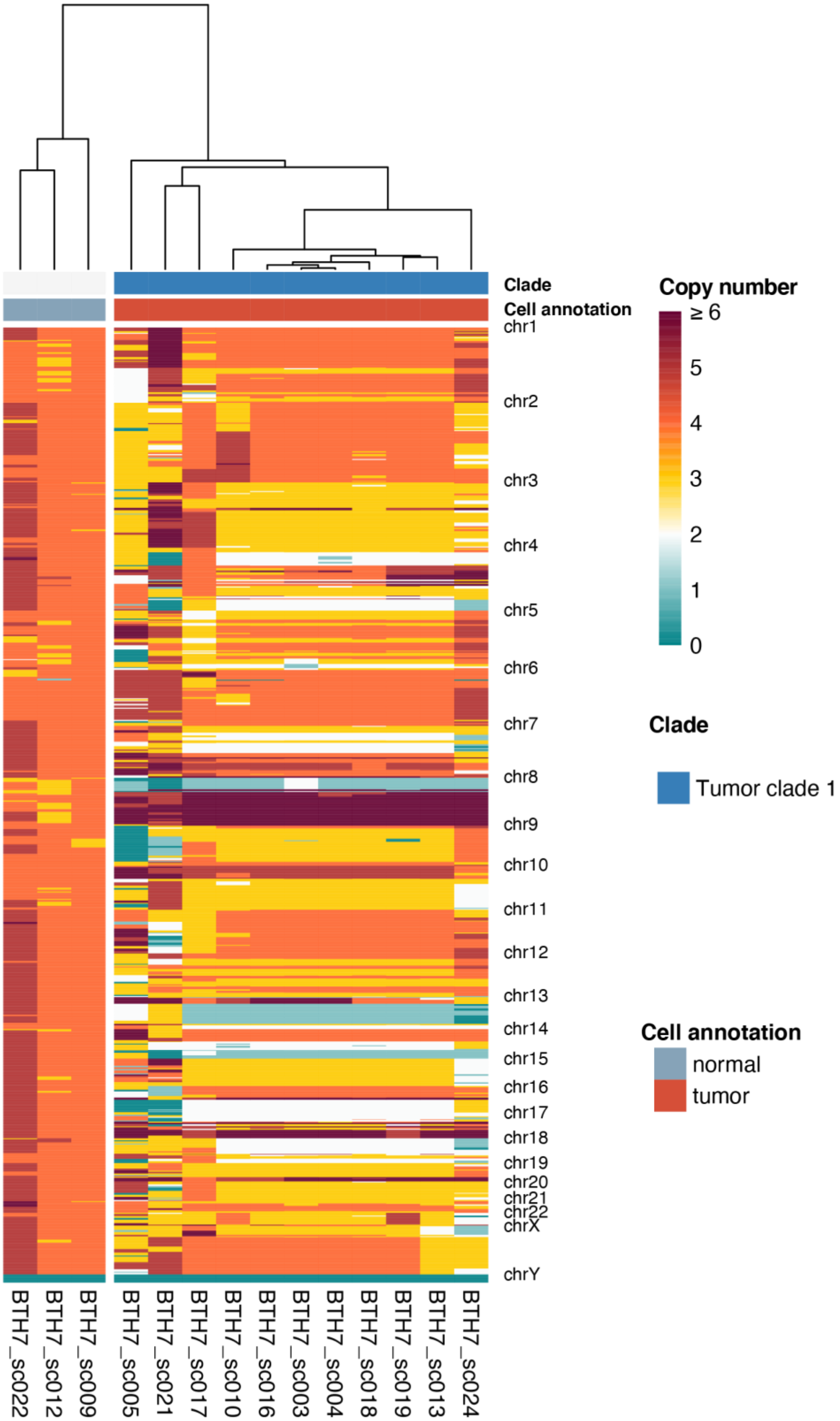


**Supplementary Fig. 26. Hierarchical clustering of CNV profiles of tumor BTH7.**

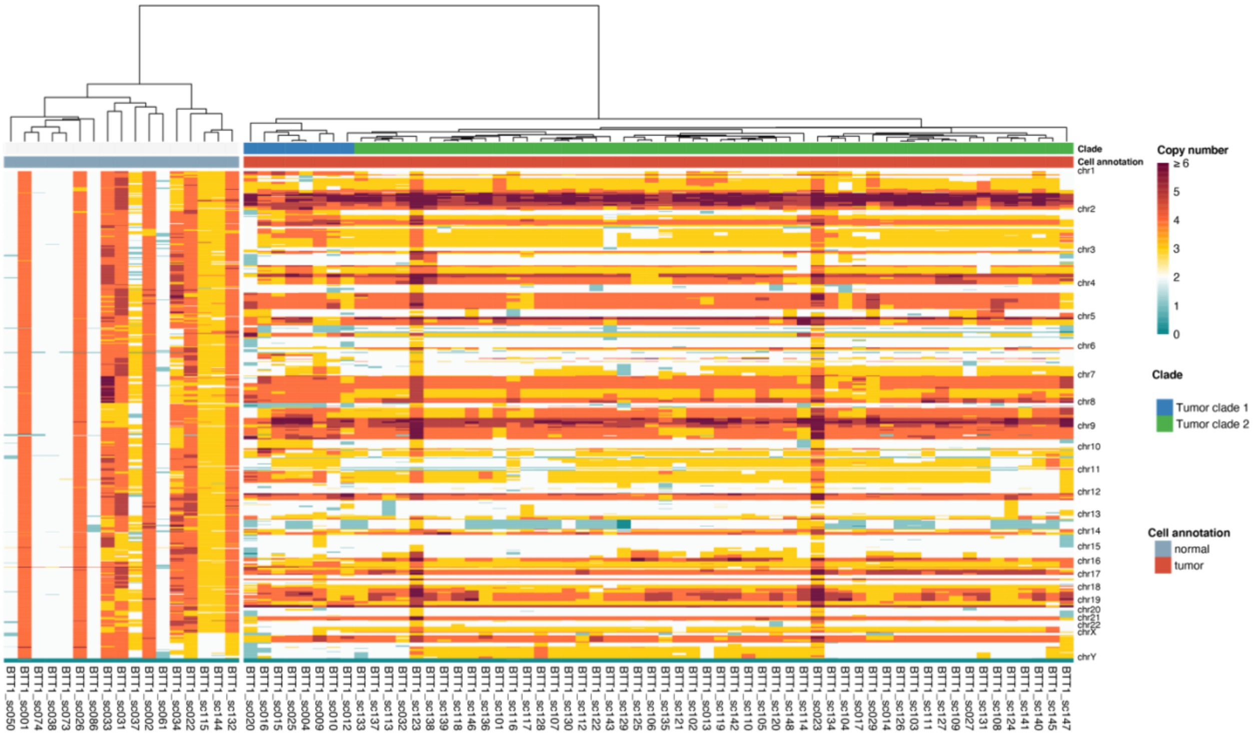

**Supplementary Fig. 27. Hierarchical clustering of CNV profiles of tumor BTT1.**

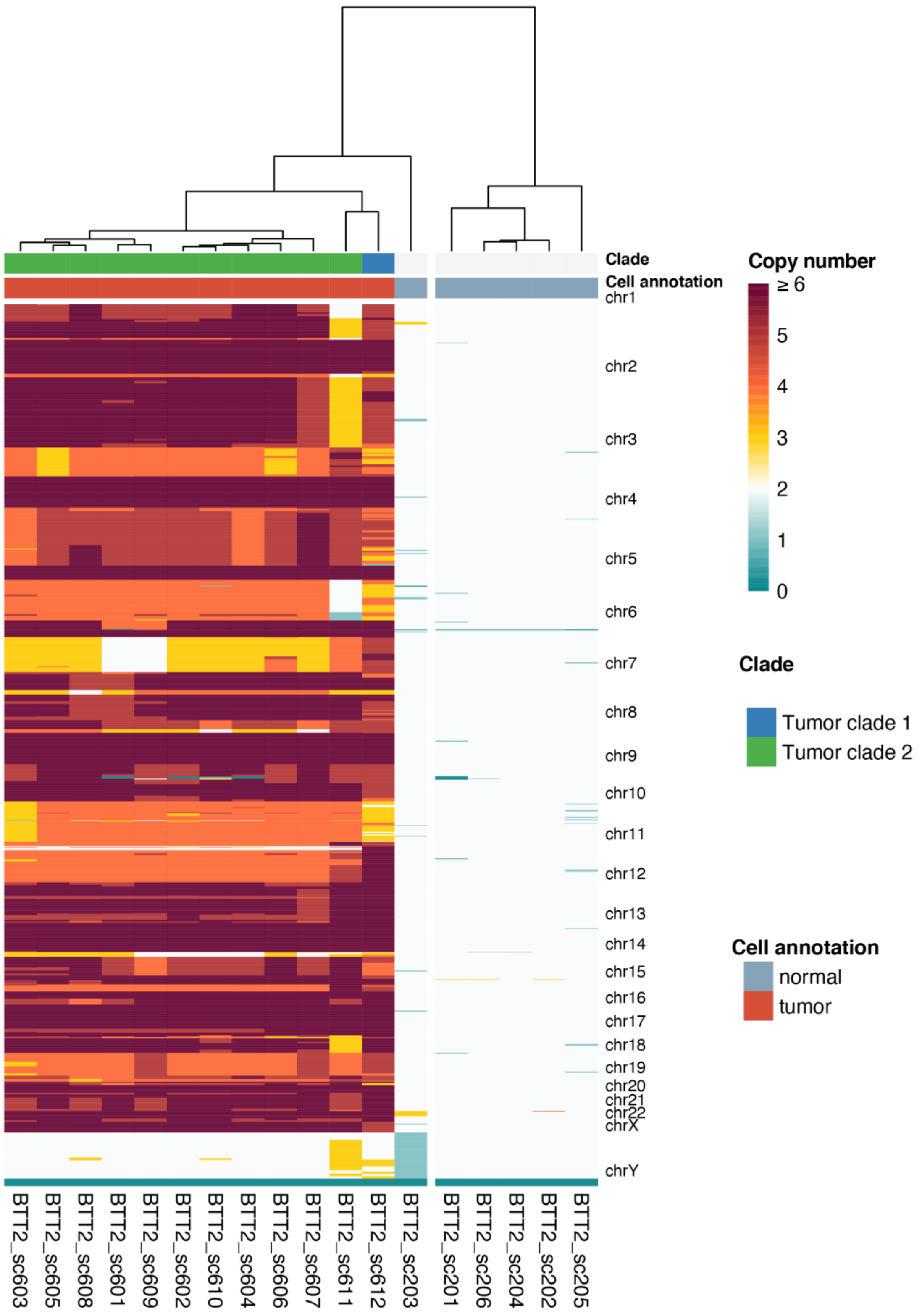


**Supplementary Fig. 28. Hierarchical clustering of CNV profiles of tumor BTT2.**

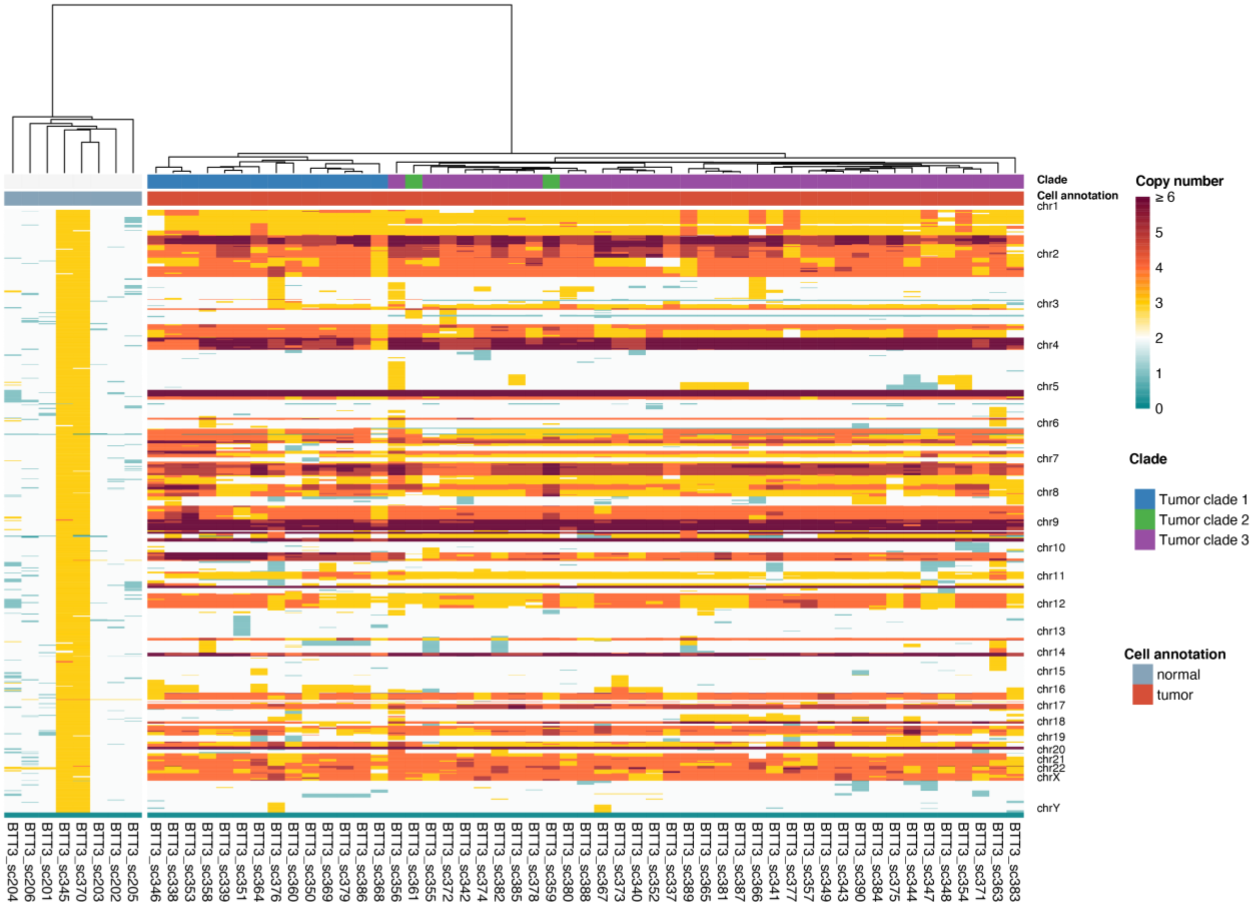


**Supplementary Fig. 29. Hierarchical clustering of CNV profiles of tumor BTT3.**

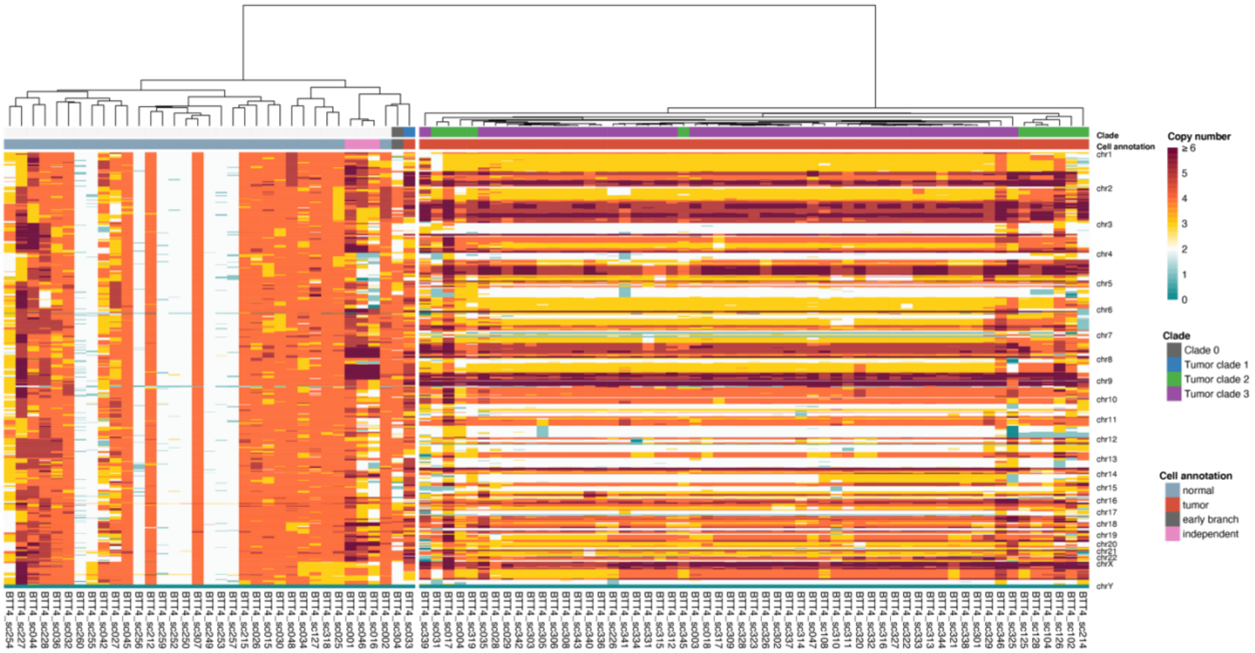


**Supplementary Fig. 30. Hierarchical clustering of CNV profiles of tumor BTT4.**

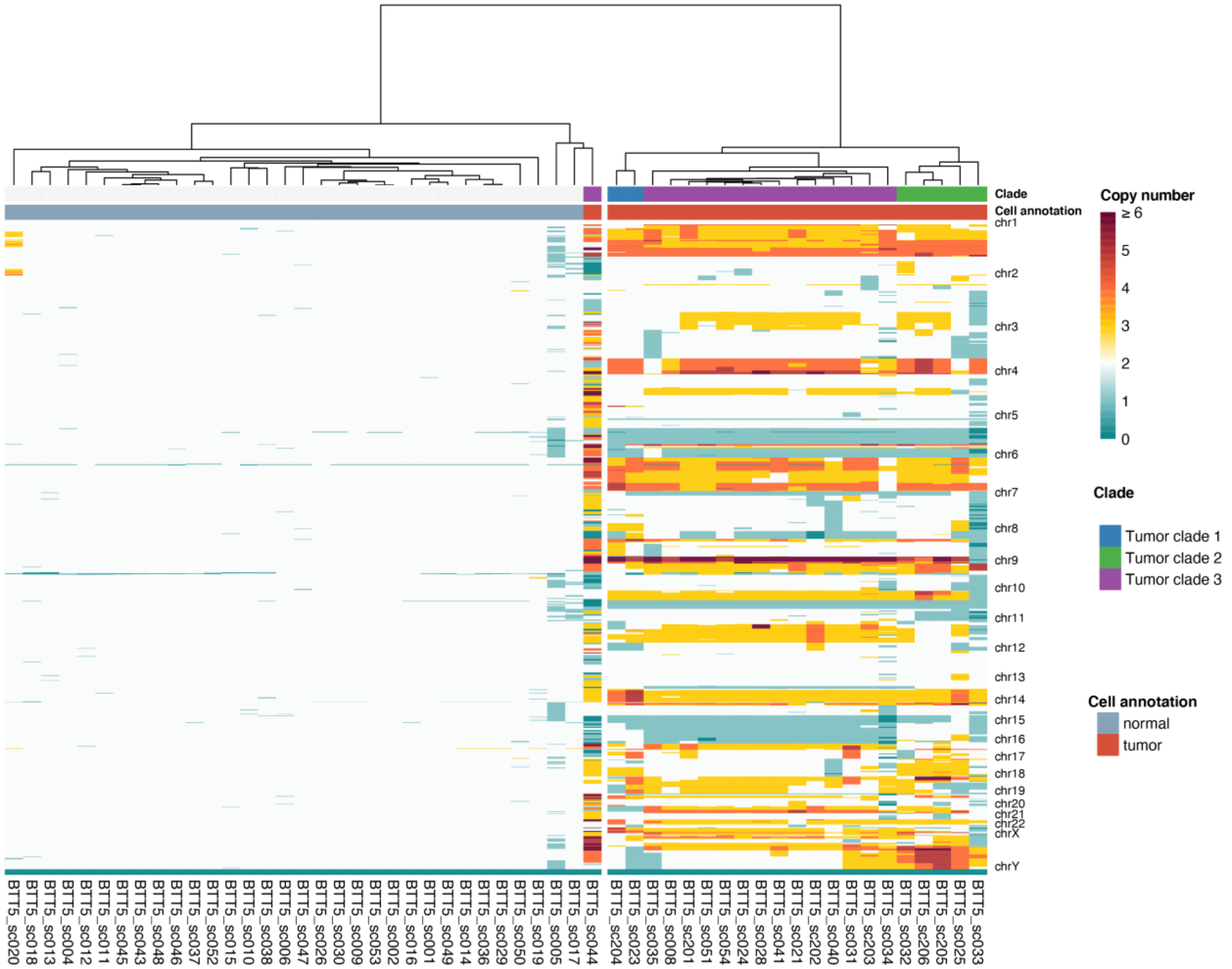


**Supplementary Fig. 31. Hierarchical clustering of CNV profiles of tumor BTT5.**

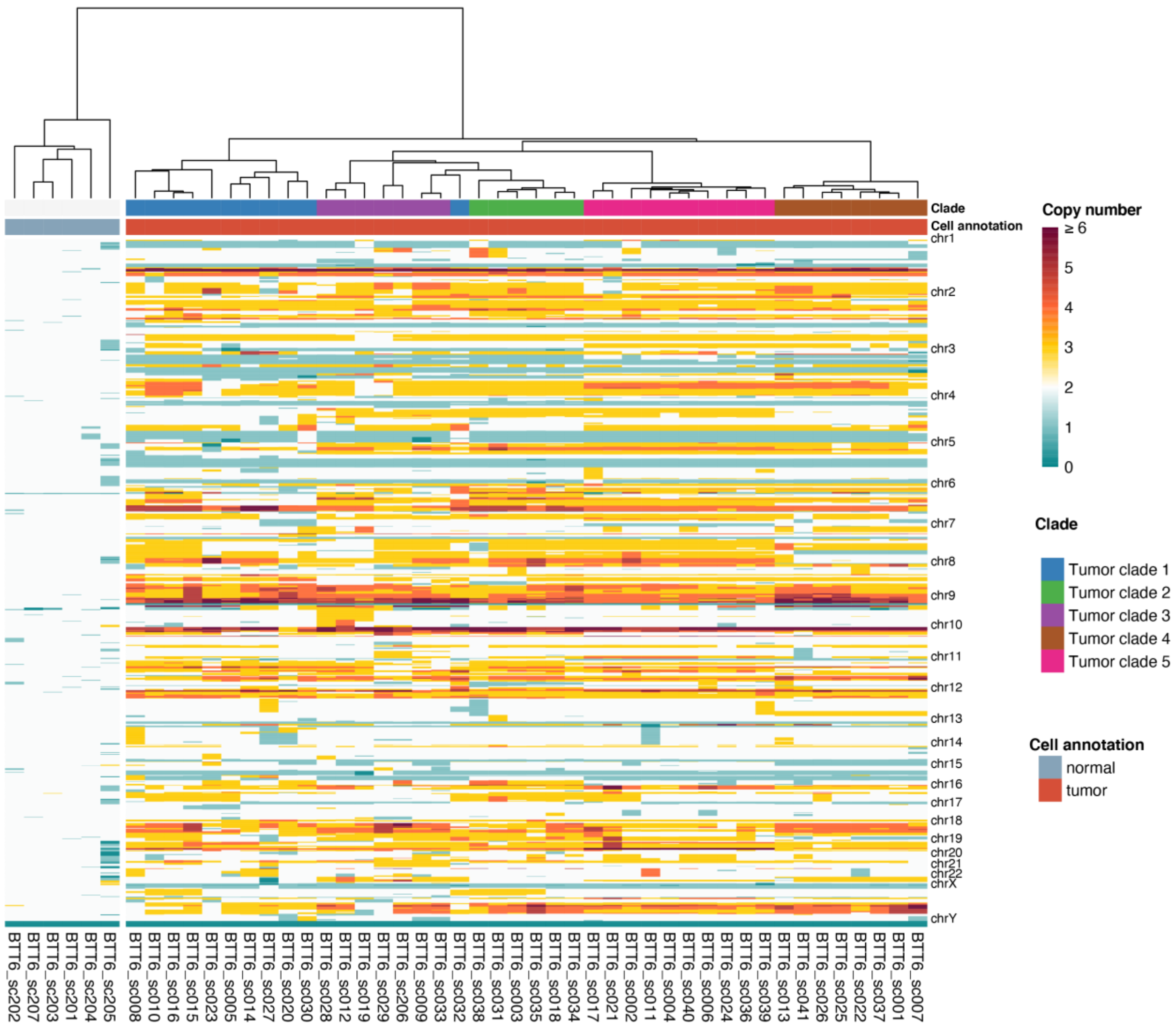


**Supplementary Fig. 32. Hierarchical clustering of CNV profiles of tumor BTT6.**

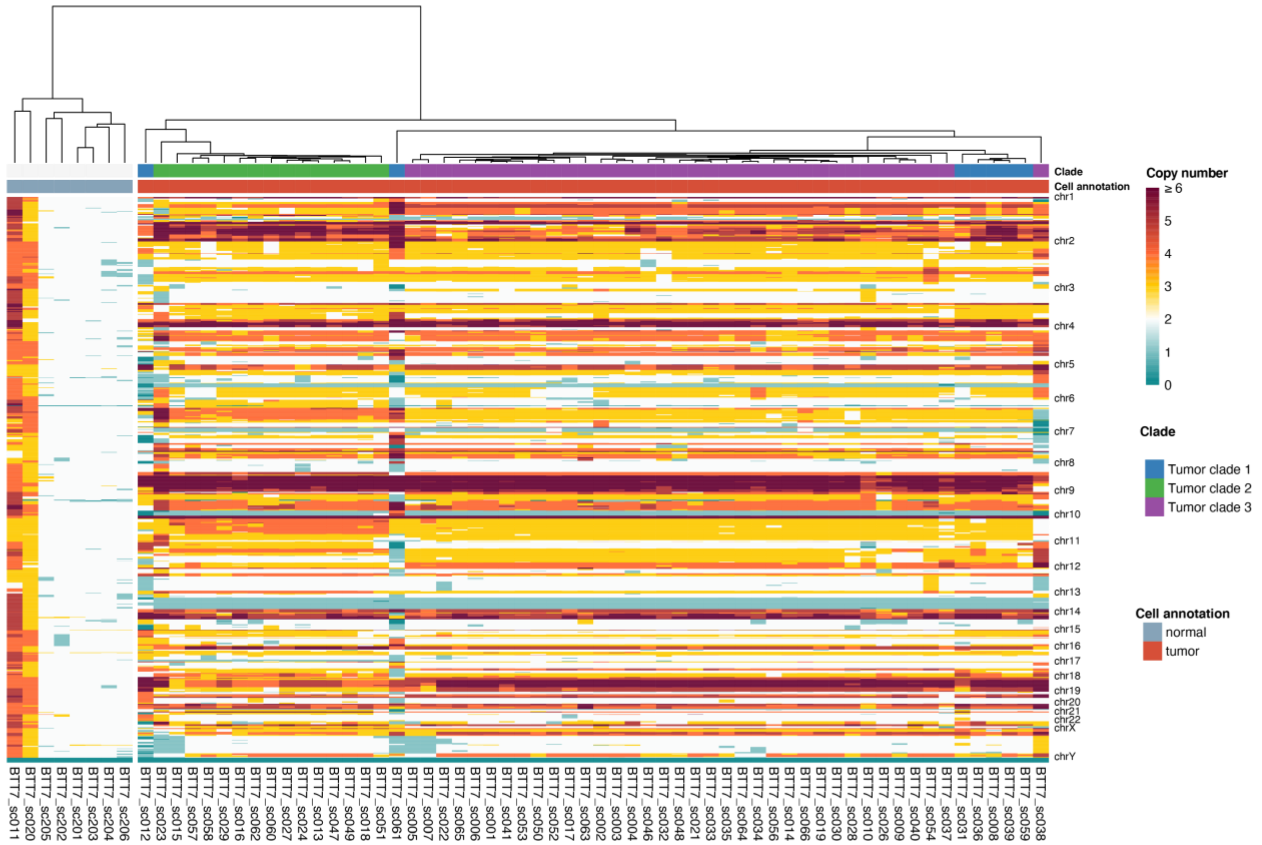


**Supplementary Fig. 33. Hierarchical clustering of CNV profiles of tumor BTT7.**

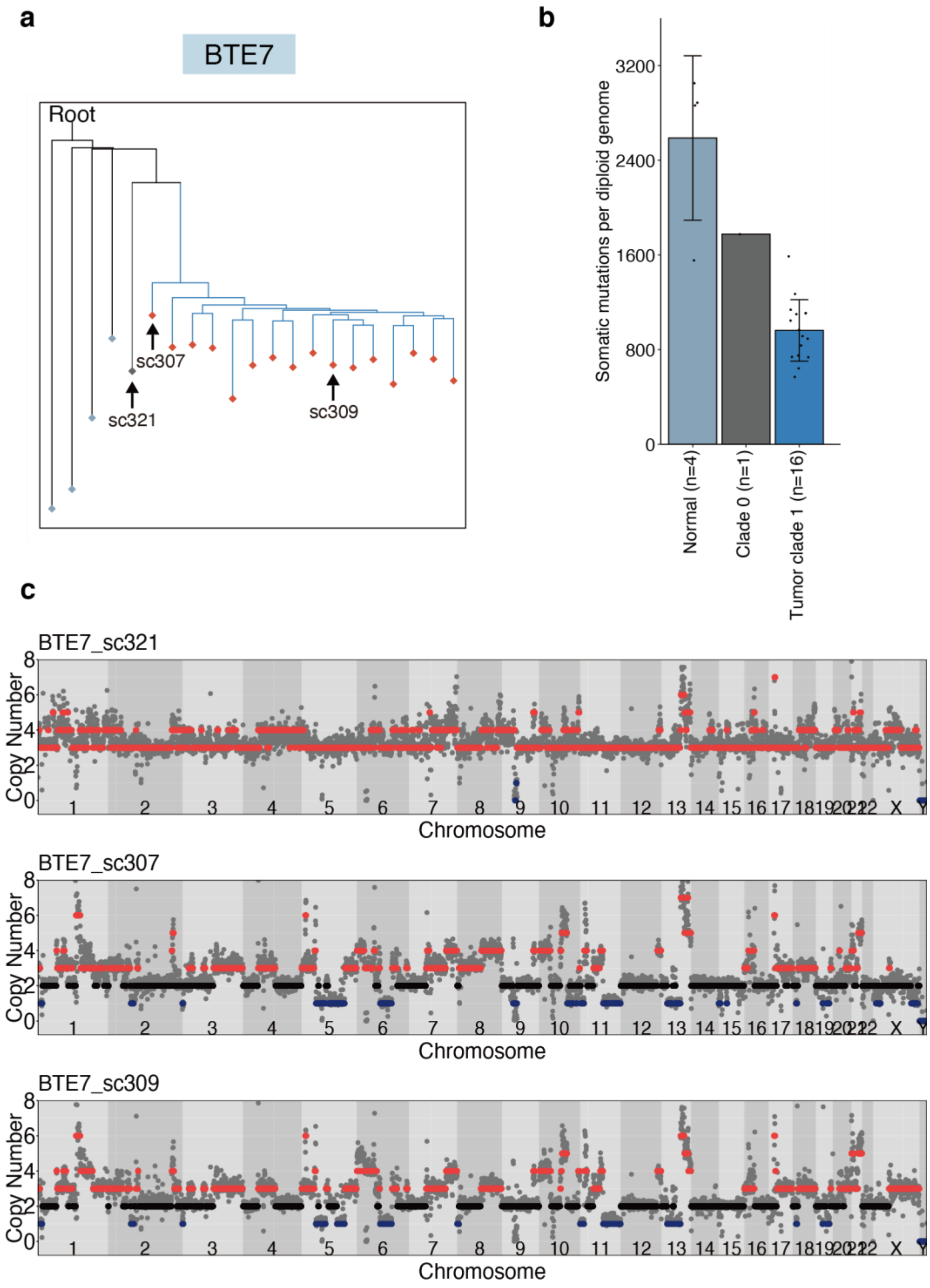


**Supplementary Fig. 34. The evolution tree of BTE7.** (**a**) The evolution tree of BTE7, with selected cells labeled. (**b**) Bar plot showing the somatic mutation load per diploid genome in different clades. (**c**) Copy number profiles of selected cells. For panel (**b**), error bars indicate standard deviation.

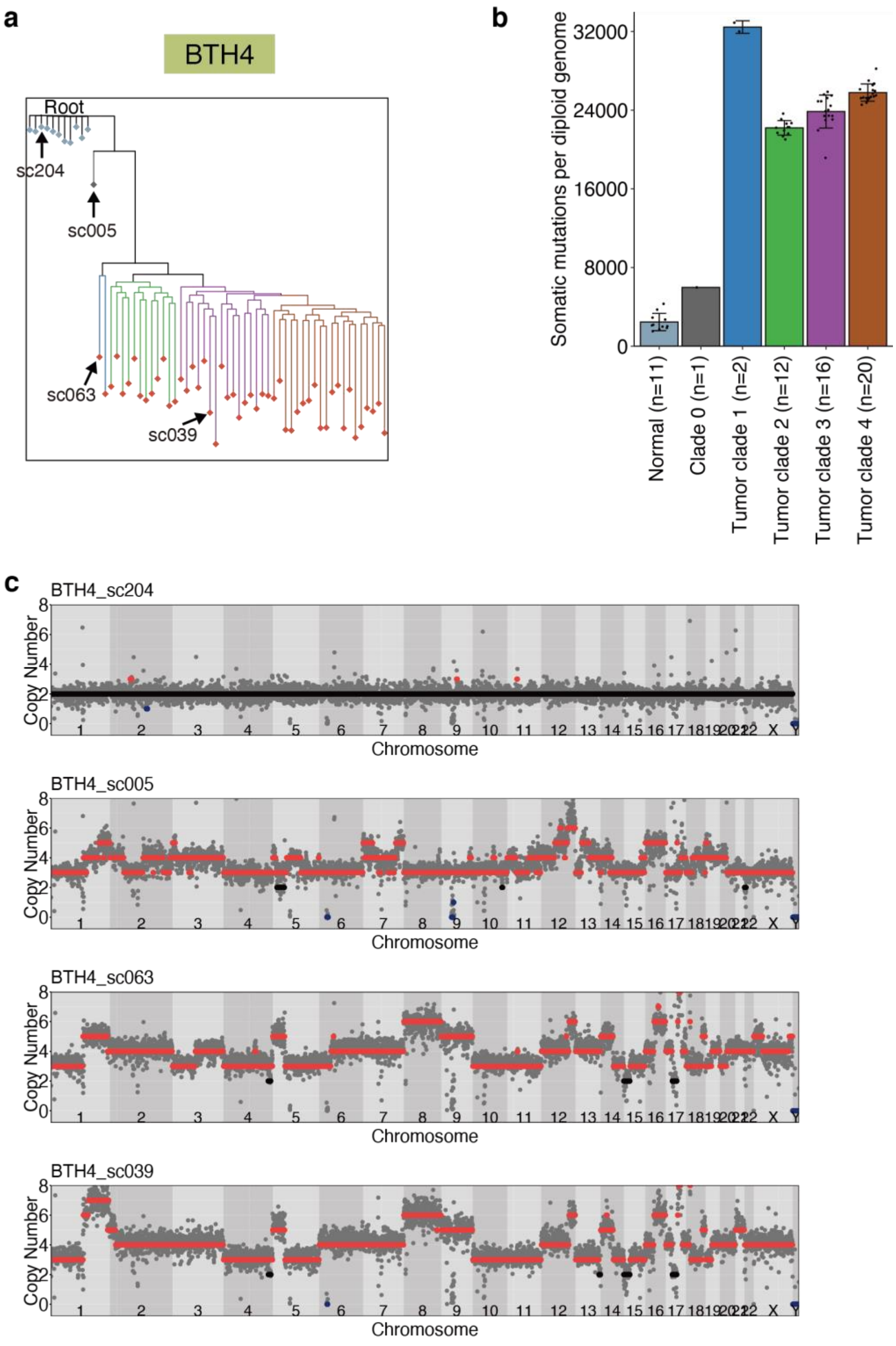


**Supplementary Fig. 35. The evolution tree of BTH4.** (**a**) The evolution tree of BTH4, with selected cells labeled. (**b**) Bar plot showing the somatic mutation load per diploid genome in different clades. (**c**) Copy number profiles of selected cells. For panel (**b**), error bars indicate standard deviation.

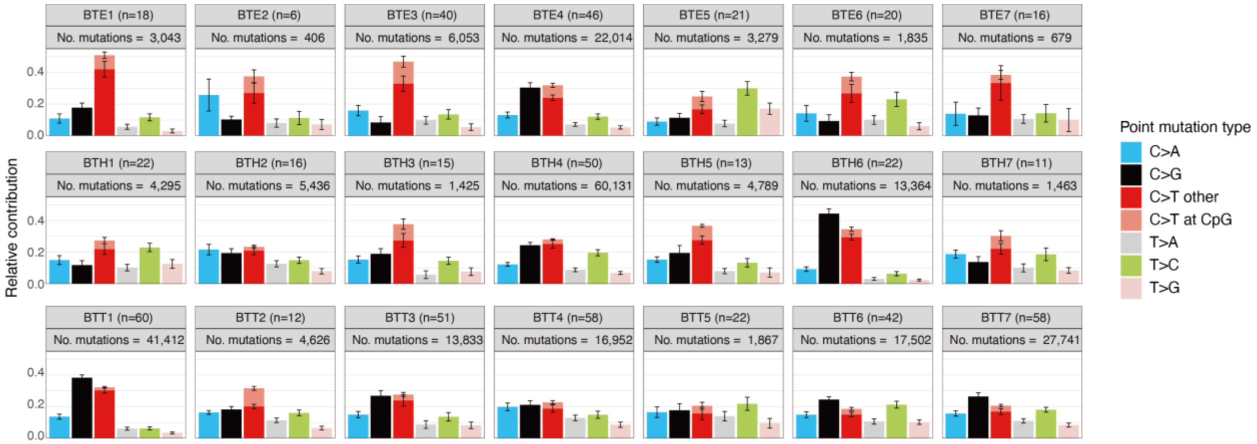


**Supplementary Fig. 36. Six-type mutation spectrum of the 21 breast cancers.** Error bars indicate the standard error of the mean.

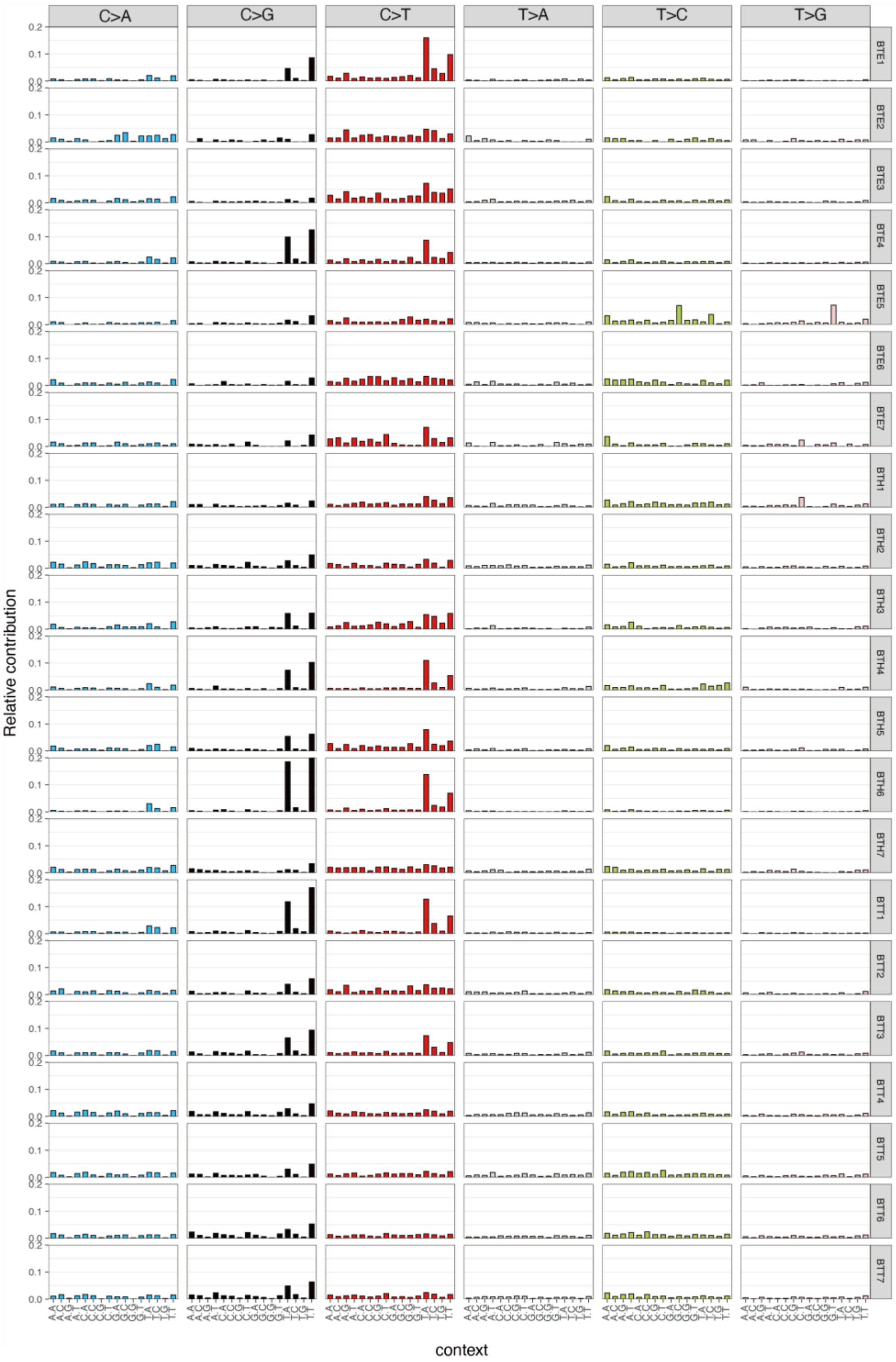


**Supplementary Fig. 37. 96-type SBS mutational profiles of the 21 breast cancers.**

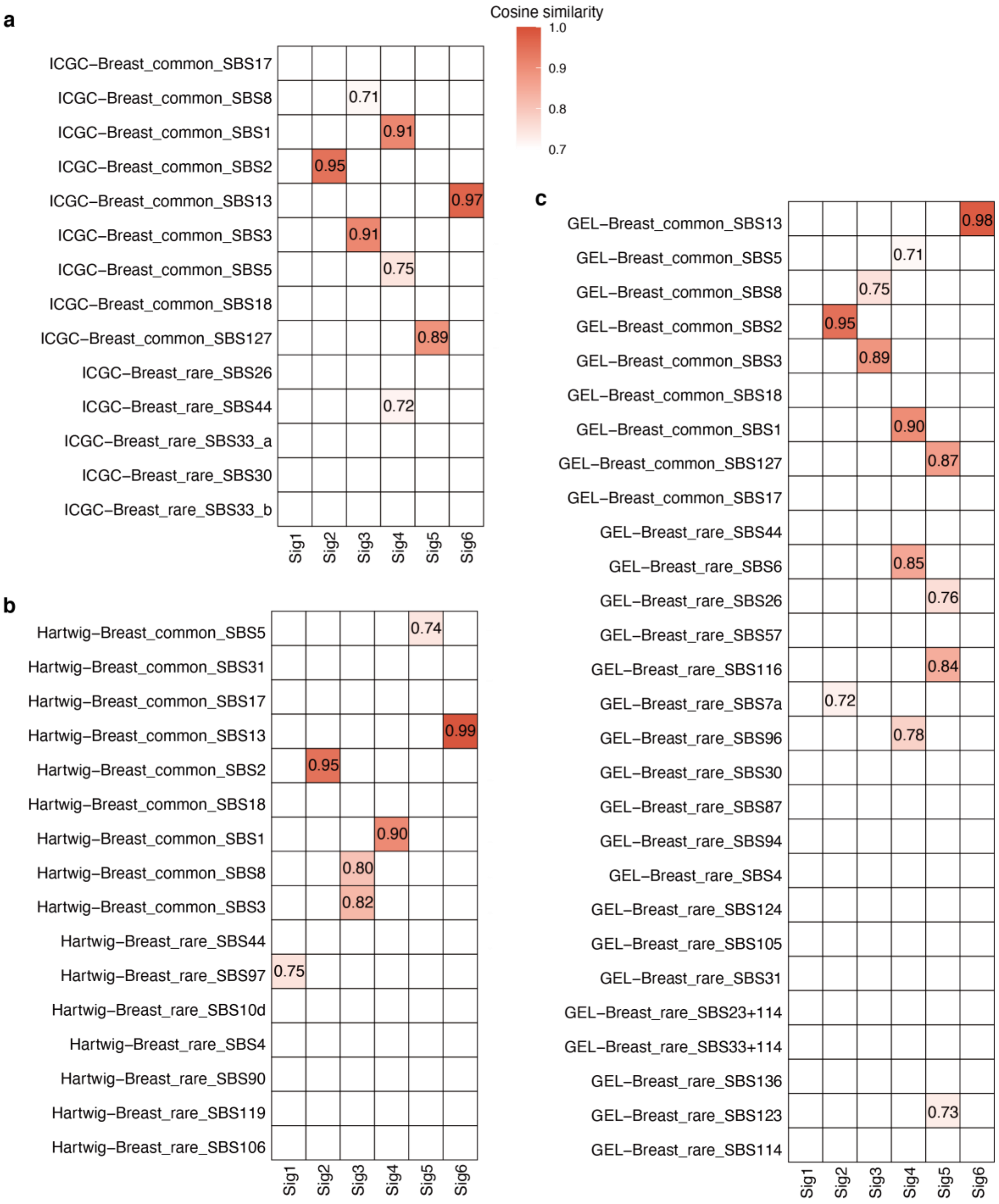


**Supplementary Fig. 38. Mapping the 6 extracted mutational signatures to Signal cancer signature database.** (**a-c**) Heatmap showing the cosine similarity between the 6 extracted signatures versus the reference signatures based on three independent breast cancer cohorts: (**a**) International cancer genome consortium (ICGC), (**b**) Hartwig, and (**c**) Genomics England (GEL).

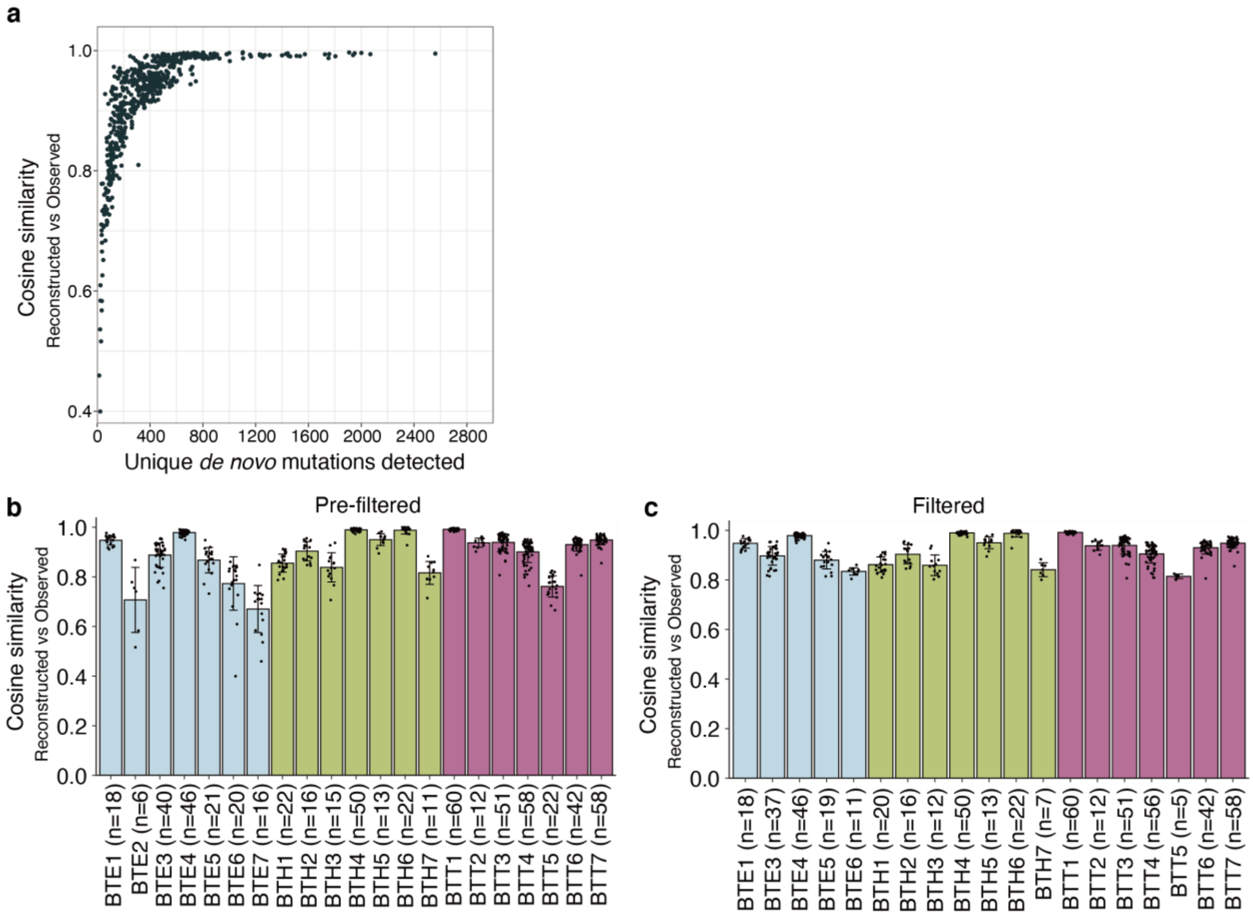


**Supplementary Fig. 39. Reconstruction of mutational signatures.** (**a**) Scatter plot showing the correlation between the signature reconstruction accuracy versus the number of somatic mutations detected in single cells. (**b**) Bar plot demonstrating the cosine similarity between reconstructed signatures versus observed signatures. (**c**) Bar plot demonstrating the cosine similarity between reconstructed signatures versus observed signatures after filtering out cells with low-quality reconstruction. For panels (**b**) and (**c**), error bars indicate standard deviation.

**Supplementary Table 1. Clinical information of the 21 breast cancer samples.**

**Supplementary Table 2. Summary information for single-cell somatic mutation load.**

**Supplementary Table 3. Somatic mutation calling by subsampling ultra-deep bulk WGS.**

**Supplementary Table 4. Quantification of stem length mutation density.**

**Supplementary Table 5. Approximation of whole tumor mutation burden by CellPhy.**

**Supplementary Table 6. Single-cell copy number profiles.**

**Supplementary Table 7. Mutational signatures identified by NMF.**

**Supplementary Table 8. Cosine similarity of the six extracted signatures compared to Signal cancer signature database.**

**Supplementary Table 9. Relative contribution of 6 mutational signatures in single tumor cells.**
